# Glioblastoma-instructed microglia transition to heterogeneous phenotypic states with phagocytic and dendritic cell-like features in patient tumors and patient-derived orthotopic xenografts

**DOI:** 10.1101/2023.03.05.531162

**Authors:** Yahaya A. Yabo, Pilar M. Moreno-Sanchez, Yolanda Pires-Afonso, Tony Kaoma, Bakhtiyor Nosirov, Andrea Scafidi, Luca Ermini, Anuja Lipsa, Anaïs Oudin, Dimitrios Kyriakis, Kamil Grzyb, Suresh K. Poovathingal, Aurélie Poli, Arnaud Muller, Reka Toth, Barbara Klink, Guy Berchem, Christophe Berthold, Frank Hertel, Michel Mittelbronn, Dieter H. Heiland, Alexander Skupin, Petr V. Nazarov, Simone P. Niclou, Alessandro Michelucci, Anna Golebiewska

## Abstract

**Background:** A major contributing factor to glioblastoma (GBM) development and progression is its ability to evade the immune system by creating an immune-suppressive environment, where GBM-associated myeloid cells, including resident microglia and peripheral monocyte-derived macrophages, play critical pro-tumoral roles. However, it is unclear whether recruited myeloid cells are phenotypically and functionally identical in GBM patients and whether this heterogeneity is recapitulated in patient-derived orthotopic xenografts (PDOXs). A thorough understanding of the GBM ecosystem and its recapitulation in preclinical models is currently missing, leading to inaccurate results and failures of clinical trials.

**Methods:** Here, we report systematic characterization of the tumor microenvironment (TME) in GBM PDOXs and patient tumors at the single-cell and spatial levels. We applied single-cell RNA-sequencing, spatial transcriptomics, multicolor flow cytometry, immunohistochemistry and functional studies to examine the heterogeneous TME instructed by GBM cells. GBM PDOXs representing different tumor phenotypes were compared to glioma mouse GL261 syngeneic model and patient tumors.

**Results:** We show that GBM tumor cells reciprocally interact with host cells to create a GBM patient-specific TME in PDOXs. We detected the most prominent transcriptomic adaptations in myeloid cells, with brain-resident microglia representing the main population in the cellular tumor, while peripheral-derived myeloid cells infiltrated the brain at sites of blood-brain barrier disruption. More specifically, we show that GBM-educated microglia undergo transition to diverse phenotypic states across distinct GBM landscapes and tumor niches. GBM-educated microglia subsets display phagocytic and dendritic cell-like gene expression programs. Additionally, we found novel microglial states expressing cell cycle programs, astrocytic or endothelial markers. Lastly, we show that temozolomide treatment leads to transcriptomic plasticity and altered crosstalk between GBM tumor cells and adjacent TME components.

**Conclusions:** Our data provide novel insights into the phenotypic adaptation of the heterogeneous TME instructed by GBM tumors. We show the key role of microglial phenotypic states in supporting GBM tumor growth and response to treatment. Our data place PDOXs as relevant models to assess the functionality of the TME and changes in the GBM ecosystem upon treatment.

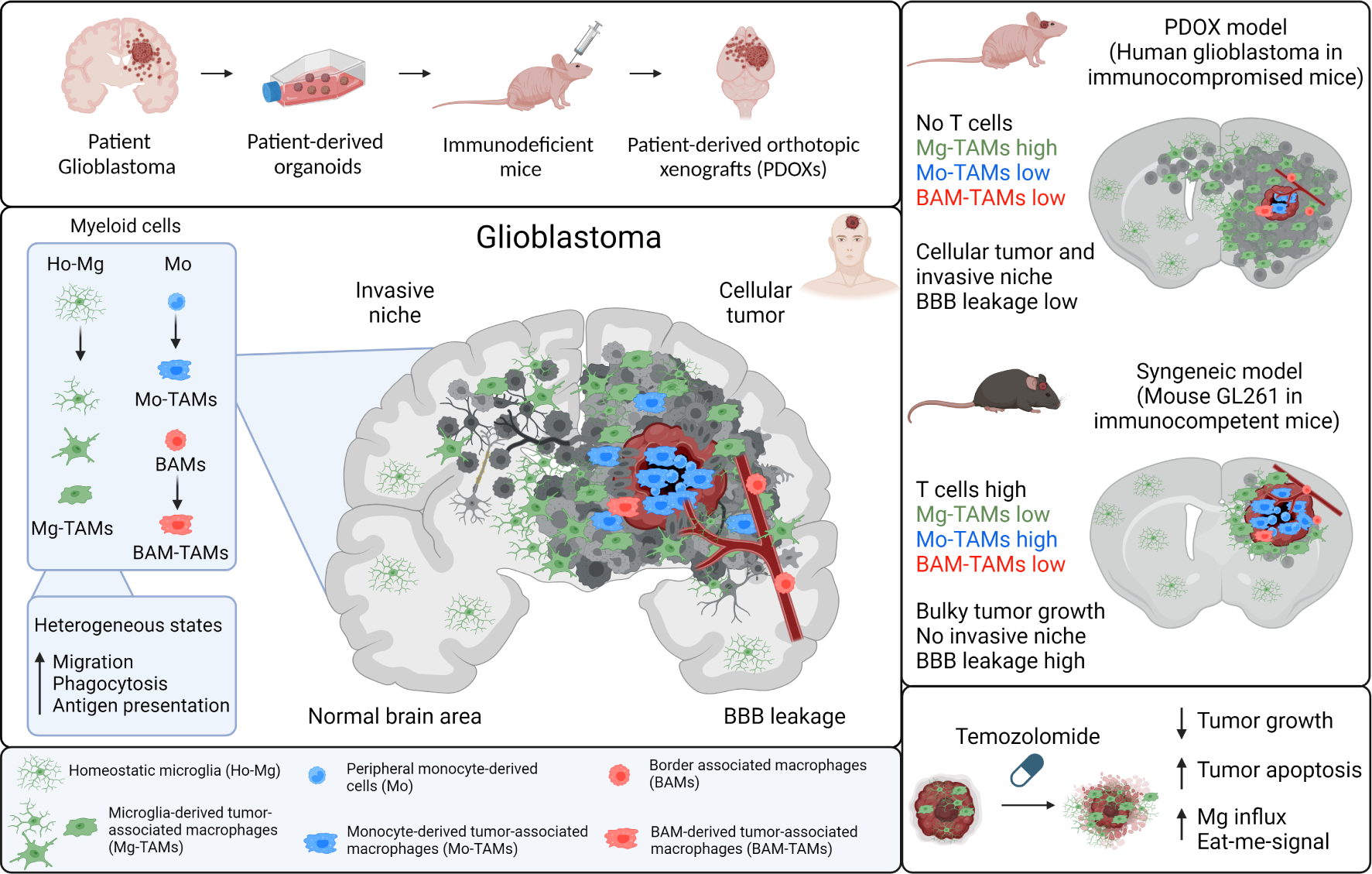

## BACKGROUND

Tumor microenvironment (TME) components, encompassing immune and non-immune non-malignant cells, have been recognized as key players in tumor initiation, progression and treatment resistance in all aggressive cancers^1^. Glioblastomas (GBMs), the most aggressive and incurable primary brain tumors, form a very dynamic ecosystem, in which heterogeneous tumor cells reciprocally interact with various cells of the TME^2^. The brain TME includes endothelial cells, pericytes, astrocytes, neurons, oligodendrocytes and immune cells. Although GBMs are identified as ‘cold tumors’ with very little lymphocytic infiltration^3^, the GBM TME contains up to 40% of myeloid cells, which, as a whole, are referred to as tumor-associated macrophages (TAMs) and are known to create a supportive environment facilitating tumor proliferation, survival and migration^4,5^. TAMs are mainly composed of cells arising from brain-resident parenchymal microglia (Mg), peripheral monocyte-derived cells (Mo) and perivascular macrophages, known as border-associated macrophages (BAMs)^6–8^. The proportions and functions of TAMs of different origins are not yet clearly defined due to the striking phenotypical and functional adaptation in the TME, lack of stable identifying markers and adequate preclinical models. Recent studies have shown that TAMs in GBM are different from classical pro-inflammatory activated (immune-permissive) M1 or alternatively activated (immune-suppressive) M2 reactive profiles^9,10^. Notably, it has been proposed that TAMs acquire different gene expression programs depending on the GBM subtype and upon GBM recurrence^11–13^. However, it is not yet well understood to what extent TAMs acquire diverse functional phenotypes *in vivo* depending on their origin, specific tumor niches, or along tumor development and progression^14^. Therefore, a better understanding of functional TAM heterogeneity will pave the way for new therapeutic strategies targeting the myeloid compartment.

To achieve this critical aim, reliable patient-derived brain tumor models are needed. The lack of in-depth comprehension of preclinical models is regularly stated as one of the main challenges in curing cancer, including brain tumors^15^. Despite being used for functional investigations of the TME, the commonly used syngeneic and genetically engineered mouse models suffer from their limited resemblance to human disease. In this context, while GBM patient-derived organoids preserve certain TME components only during initial days, and *ex vivo* co-culture protocols are still immature^16^, patient-derived xenografts allow for propagation of primary patient tumors in less selective conditions than in *in vitro* cultures^17^. Additionally, as subcutaneous xenografts do not recapitulate the natural TME, patient-derived orthotopic xenografts (PDOXs) implanted in the brain are certainly more adequate for modeling gliomas. However, although showing an excellent recapitulation of GBM tumor features, PDOXs are characterized by an immunocompromised environment and the replacement of the human TME with mouse counterparts. It is therefore important to assess to what extent PDOXs can mimic major TME features observed in GBM patient tumors.

We have previously reported that PDOX models recapitulate the genetic, epigenetic and transcriptomic features of human tumors^18^. We have also shown that mouse cells interact with human GBM cells in the brain. In particular, endothelial cells forming blood vessels adapt their morphology and molecular features analogous to the aberrant vasculature observed in patients^19^. We have further shown that mouse endothelial cells respond to anti-angiogenic treatment, as observed in GBM patients, leading to normalized blood vessels and treatment escape mechanisms toward more invasive tumors^20^. Here, we further inferred the heterogeneity of the TME compartment in PDOXs across genetically and phenotypically diverse GBM landscapes as well as upon temozolomide (TMZ) treatment. Focusing on TAMs, we distinguish their origin across GBM landscapes and identify distinct molecular programs. We show that Mg are a key component of the TME in GBM and transition toward heterogeneous phenotypic states in tumors of diverse genetic backgrounds. By interrogating single-cell RNA-sequencing (scRNA-seq) and spatial transcriptomic profiles of GBM patient tumors, we confirm that the identified Mg states in PDOXs are also abundant in patients and localize variably across spatial TME niches. Notably, high proportions of these Mg-TAMs present dendritic cell-like features, including enhanced phagocytic and antigen-presenting cell characteristics. Finally, we show that TMZ modulates the molecular features of the tumor and TME components, leading to a differential network between various components of the GBM ecosystem.

## MATERIALS AND METHODS

### Clinical GBM samples and PDOXs

We collected GBM tissues from the Centre Hospitalier de Luxembourg (CHL, Neurosurgical Department) or the Haukeland University Hospital (Bergen, Norway) from patients who had given their informed consent. PDOX derivation and maintenance via serial transplantation of primary organoids in NOD/SCID (NOD. CB17-Prkdc^scid^/J) or NSG (NOD. Cg-Prkdc^scid^ Il2rg^tm1WjI^/SzJ) was previously described^18,19,21^ (**Table S1**). See **Supplementary material** for PDOX characterization at multiomics level. For specific flow cytometry-based experiments, patient-derived organoids were implanted into eGFP-expressing NOD/Scid mice as described previously^22^. For the mouse-derived TME analysis, tumor organoids were implanted (6 per brain) into the right frontal cortex of female nude mice (athymic nude mice, Charles River Laboratories, France). Animals were housed in a specific pathogen-free (SPF) facility under a controlled environment (temperature, humidity, and light) with free access to water and food. Animals were sacrificed at the endpoint defined in the scoresheet. For the time point experiment, P3 PDOXs in nude mice were sacrificed also 25 and 35 days post implantation, following tumor growth evaluation by MRI. Mice were deeply anesthetized with a mixture of ketamine (100 mg/kg) and xylazine (10 mg/kg) and transcardially perfused with ice-cold PBS.

For TMZ treatment, P3 PDOXs were evaluated daily. Tumor growth was monitored by MRI (3T MRI system, MR Solutions). At day 30, mice were randomized into 2 groups: from day 33 control PDOXs were administered NaCl 0.9% + 10% DMSO, treatment group received 40 mg/kg TMZ in NaCl 0.9% + 10% DMSO corresponding to 120 mg/m^2^ in humans. Treatment was administered by oral gavage 5x per week with a total of 8 doses, where the last treatment dose was given shortly before the endpoint. Tumor growth was followed by MRI at days 37 and 42 and quantified as previously described^21^. Statistical differences were assessed with two-tailed Student’s t-test.

### scRNA-seq in PDOXs

We extracted tumor tissues from mouse brains and dissociated with the MACS Neural Dissociation kit (Miltenyi Biotec) according to the manufacturer’s instructions. Single cells were purified with the Myelin Removal Beads II kit (MACS Miltenyi Biotec) as previously described^21^. To separate human GBM cells from the mouse TME, PDOX-derived cells were FACS-sorted (P8, nude control brain)^23^ or MACS-purified (remaining PDOXs) with a Mouse Cell Depletion kit (Miltenyi Biotec)^21^. Except for tissue dissociation, all steps were performed on ice. Mouse-derived TME was processed via Drop-seq. See **Supplementary material** for human tumor cells.

Drop-seq and data preprocessing were performed as previously described^18,24^. Briefly, scRNA-seq analysis was performed in R (v4.1.1) with the Seurat package (v4.0.5)^25^. Human and mouse cells were separated by mapping the scRNA-seq reads to human g38 and mouse mm10 reference genomes. The distributions of UMI counts and features expressed allowed for clear cell separation. Mouse TME samples were merged with the published Drop-seq dataset of the GL261 TME^24^. QC thresholds were empirically applied per sample, and only genes expressed in at least 5 cells, cells expressing at least 200 features and cells with 30% or fewer mitochondrial reads were selected. Potential doublets were predicted and removed using DoubletFinder (v2.0.3)^26^. Counts were normalized using the Seurat-based ‘NormalizeData’ function, batch correction was performed by Harmony (v0.1.0)^27^. Clustering was performed on harmony embeddings using the default parameters of the Seurat package. Dimensionality reduction was performed using the Uniform Manifold Approximation and Projection (UMAP) of the Seurat package. Differential expression analysis was performed using the Wilcoxon rank sum test, and the false discovery rate (FDR) was calculated using the Benjamini-Hochberg method. Cell clusters were identified based on the expression of known marker genes and differentially expressed genes (DEGs) were determined by the ‘FindAllMarkers’ function. To assess the identity of cells identified as a cycling state, the ‘Cycling’ cluster was extracted as a separate Seurat object. We used UCell method^28^ to score each single cell by the gene signatures of other cell types (i.e. clusters) identified in the study and we labelled each cell with the cell type that had max UCell modular score calculated.

The myeloid cluster was extracted using the ‘Subset’ function. Gene ontology analysis was performed by METASCAPE (https://metascape.org/). Single-cell trajectory inference analyses were performed on Mg cells with the TSCAN R packages using default parameters (doi:10.18129/B9.bioc.TSCAN) to obtain minimum spanning tree and pseudotime ordering of cells. The z-score of genes was calculated by subtracting the mean of expression from the raw expression of each gene and normalization by the corresponding standard deviation. Gene expression was displayed as heatmap of z-scores. Single-cell gene set signature scores were calculated using the Seurat ‘AddModuleScore’^29^. Identification of master transcriptional regulators was performed using normalized counts from subsetted myeloid cells. Gene regulatory network inference was performed according to the standard SCENIC workflow^30^.

### Reference Based Mapping

Myeloid cells were identified and extracted from publicly available GL261 scRNA-seq datasets based on the expression of key myeloid cell markers (“*Itgam*”,“*P2ry12*”,“*Csf1r*”,“*Tgfbi*”, “*Ptprc*”, “*Hexb*”,“*Mrc1*”, “*Ly6c2*”)^7,8,24^. Dendritic cells (DCs) were excluded by identifying clusters expressing key DC markers reported in Pombo et al.. The Ochocka et al. dataset was used as the reference. Other datasets were projected onto the reference UMAP structure. The reference principal component analysis (PCA) space was computed using 2000 most variable genes, and the first 30 PCs were used to calculate the UMAP model. Next, we determined the common features of the reference and each of the query datasets by Seurat’s ‘FindTransferAnchors’ function with the reduction method ‘pcaproject’ and the parameter ‘dims = 1:50’. Finally, we called ‘MapQuery()’ to transfer cell type labels and project the query data onto the UMAP structure of the reference.

### Cell-to-cell communication analyses

Cell-to-cell communications were inferred using CellChat(v1.6.0)^31^. Human genes in GBM tumor cells from PDOXs were converted to mouse homologs using the ‘convert_human_to_mouse_symbols’ function in the nichenetr package (v1.1.1)^32^ and merged with the TME matrix from the corresponding PDOX. The subpopulation was considered for the analysis if at least 10 cells per subpopulation were present in the sample(s) considered. Crosstalk inference analyses were performed using the ‘CellChatDB.mouse’ database on each of the conditions before they were merged for comparison. The exploration, analysis and visualization of inferred networks were performed using default parameters of relevant CellChat functions. Other visualization packages of R, such as ggplot2 and patchwork, were used to improve the quality of plots and plot annotations.

### Human GBM scRNA-seq and bulk RNA-seq analyses

Where applicable, mouse MGI gene symbols were automatically converted to human HGNC symbols in R using the capital letter (*toupper*) or the ‘getLDS’ function in the biomaRt ^33^. Only genes with human homologs available in http://www.informatics.jax.org/downloads/reports/HOM_MouseHumanSequence.rpt were applied.

#### scRNA-seq

Analysis of the TME in human GBM was performed using the Darmanis et al., dataset^34^. Analysis of myeloid cells was performed using publicly available 10X Genomics GBM datasets^8,35–37^ and an annotated GBmap database^38^. 10x Genomics GBM datasets were obtained as preprocessed gene expression matrices (DEMs) from a total of 36 IDH wild-type tumors including newly diagnosed (n = 27) and recurrent (n = 9) GBMs. GBmap data were obtained from a total of 110 IDH wild-type GBM tumors, 103 tumors were annotated as newly diagnosed or recurrent and contain sufficient amount of myeloid cells for the analysis. Each dataset was analyzed separately to extract myeloid cells. Myeloid cells extracted as follows: (1) For GBmap and Friedrich et al., dataset, cells were identified according to the author’s annotation; (2) For Wang et al., Johnson et al., and Pombo-Antunes et al., datasets, we used an approach that combines overexpression (OE) and clustering analysis. First, genes with zero count in all cells were filtered out, and the ‘NormalizeData’ function was applied to LogNormalize each cell with a scale factor of 10,000. UMAP was used to visualize the cells and clusters in 2 dimensions. Cluster identity was determined according to overrepresentation of a cell type within the cluster as called by OE analysis. Myeloid cells were extracted for further analysis. Overall, 51,302 myeloid cells were extracted and combined into one Seurat object. Genes with zero counts in all cells were removed, and cells were log-normalized with a scale of 10,000. The Harmony (v0.1.0) package was used to remove variation due to batch effects. All PCs were used and *theta* was set to 1. Harmony embedding was used for clustering analysis. A subset of myeloid cells including Monocytes, TAM-BDM and TAM-MG cells from the GBmap Seurat object’s ‘annotation_level_3’ were scored using our Human Mo and Human Mg signature gene lists. Myeloid cells were assigned to either Mo or Mg based on the highest single-cell gene set signature scores between Mo or Mg gene sets for each cell. A heatmap showing the overexpression scores (OES) of cell type markers, functional signatures and cluster markers in the Pombo-Antunes et al., dataset was generated using pHeatmap (v1.0.12). OES were calculated using Seurat’s ‘addModuleScore’ function. Cells (columns) were ordered according to hierarchical clustering analysis based on OES of Mo and Mg gene signature lists.

#### Bulk RNA-seq

The Ivy Glioblastoma Atlas Project (IVY-GAP) bulk RNA-seq dataset was obtained from 279 patients tumor fragments in total (anatomic structure cohort: 122 samples from 10 tumors; Cancer Stem Cell cohort: 157 samples from 34 tumors)^39^. To generate the heatmap of the OES of signatures, OES of the gene lists were calculated using the normalized z-score as done in Jerby-Arnon et al.^40^ The OES table was then split according to the tumor location, and heatmaps of OES were generated for each tumor location using ComplexHeatmap (v2.12.1)^41^. Bulk RNA-seq profiles of CD49d^+^ (MG) and CD49d^−^ (MDM) myeloid cells purified from normal human brain (n=7) and GBM patient tumors (n=15) were analysed via https://joycelab.shinyapps.io/braintime^42^.

#### Survival analysis

For the survival analyses, IDHwt GBMs were selected from the CGGA dataset (n= 249 tumors in batch 1 and 109 tumors in batch 2 combined)^43^. IDH mutant and unknown status tumors were removed. IDHwt GBMs were stratified into three distinct transcriptomic subtypes: classical, mesenchymal and proneural following previously reported signatures^12^. Additional clinical parameters included treatment status of the patient and *MGMT* promoter methylation status. Scores for the signatures used were computed for each sample using the ssGSEA method from the package GSVA (v.1.38.2)^44^ in a purrr tidyverse environment leading to a matrix of enrichment score of each signature (rows) and for each sample (columns). The samples have been stratified into “high-score” or “low-score” by splitting the top 25% highest ssGSEA scores and the top 25% lowest ssGSEA score respectively. The stratified samples were subjected to the Kaplan-Meier analysis to estimate the survival with the package *survival* (v3.2-11).

### Spatial transcriptomics

Spatial transcriptomic profiles of 16 GBM patient tumors obtained from 16 individual patients were obtained as described recently^45^. For spatial data analysis, we acquired spatially resolved RNA-seq datasets using the SPATAData package (https://github.com/theMILOlab/SPATAData). For annotation of the scRNA-seq dataset to spatial transcriptomic data, we humanized the genes using the mice2human database (http://www.informatics.jax.org/downloads/reports/HOM_ MouseHumanSequence.rpt) and rejected all genes that failed to map to the human transcriptome. Spatial correlation analysis was performed by either a spatial Lag model or a Canonical Correlation Analysis (CCA) using the ‘runSpatialRegression’ function from the SPATAwrappers package. Cell type deconvolution of each spot was performed by Robust Cell Type Decomposition (RCTD) a well-validated toolbox. The deconvolution was performed by the SPATAwrapper (https://github.com/heilandd-/SPATAwrappers) package using the function runRCTD. Visualization of surface plots or correlation analysis was performed by the SPATA2 toolbox.

### Reference mapping to GBmap dataset

To map the scRNA-seq dataset to the human GBmap reference dataset (110 GBM patient tumors), we humanized the genes using the mice2human database (http://www.informatics.jax.org/downloads/reports/ HOM_MouseHumanSequence.rpt) and rejected all genes that failed to map to the human transcriptome. Next, we used an optimized version of azimuth (’modified_azimuth.R’) to map the query scRNA-seq dataset to GBmap^38^ as described recently. The mapping results were visualized in ref.umap.

### Immunohistochemistry

The regular histological analysis of PDOX models (H&E, human Nestin/Vimentin, mouse CD31, Ki67) was performed as described previously^18,19^ at the endpoint of tumor growth, unless specified otherwise. Antibodies are listed in **Table S2.** Mouse Iba1 staining was performed on coronal 4 - 8 µm sections from paraffin-embedded brains. Sections were incubated for 30 min at 95°C in retrieval solution (Dako). Primary antibodies were incubated overnight at 4°C or 3 hours at room temperature, followed by 30 min incubation with secondary antibodies. The signal was developed with the Envision+ System/HRP Kit in 5–20 min (K4007, Agilent/Dako). Iba1^+^ cells were quantified based on the ImageJ plugin ^46^. The quantification of Iba1^+^ cells was performed using several technical and biological replicates. Number of images per mouse (technical replicates) depended on the tumor size. Each biological replicate was calculated as a mean of technical replicates obtained across the normal brain/tumor tissue in the same mouse. Human Iba1 staining was performed according to standardized protocols using Discovery XT automated immunostainer (Ventana Medical Systems, München, Germany) as previously published^10^. For immunofluorescence, brains were perfused and post-fixed with 4% paraformaldehyde (PFA)/sucrose for 48 hours. Coronal sections (4 - 12 µm) were permeabilised with PBS with 1.5% Triton X-100, blocked with 5% BSA and incubated with the primary antibodies. Secondary antibodies were incubated for 2 h. Alternatively, an Opal 3-Plex Manual Detection Kit (Akoya Biosciences) was used following the manufacturer’s guidelines. Cell nuclei were counterstained with Hoechst (1 mg/ml; Sigma). Sections were mounted on glass slides cover slipped using Fluoromount^TM^ Aqueous Mounting Medium (Sigma). Images were obtained using a Nikon Ni-E or Zeiss LSM880 confocal microscope. Z-stacks were performed with 0.5 μm steps in the Z direction, with a XY resolution of 1.024 x 1.024 pixels.

### Multicolor flow cytometry

Animals were collected at the endpoint of tumor growth. Animals were perfused with ice-cold PBS. PDOX brains were dissected into separate zones when specified: tumor core (cellular tumor, including pseudopalisading/hypoxic zone if present), invasive zone (corpus callosum and front left hemisphere, P3) and distant zone (back left hemisphere, P3 & P13). P8 PDOX was not dissected into distinct tumor zones due to its very invasive nature and similar tumor cell density in the left and right hemispheres. Patient tumors, PDOX tumors and control mouse brains were dissociated with a MACS Neural Tissue Dissociation Kit (P) (Miltenyi) following the manufacturers’ instructions. Analysis of human cells in eGFP^+^ mice was performed as described previously^22^. For analysis of mouse-derived TME in nude mice, single cells were resuspended in ice-cold HBSS, 2% FBS, 10 mM HEPES buffer (100 µl/test). Fc receptors were blocked with CD16/CD32 antibody for 30 min at 4°C. Cells were incubated with the appropriate pre-conjugated antibodies for 30 min at 4°C in the dark (**Table S2**). Non-viable cells were stained with Hoechst (0.1 µg/ml, Sigma). Data acquisition was performed at 4°C on a FACS Aria^TM^ SORP cytometer (BD Biosciences) fitted with a 640 nm (40 mW) red laser, a 355 (20 mW) UV laser, a 405 nm (50 mW) violet laser and, a 561 nm (50mW) yellow/green laser, and a 488 nm blue laser (50 mW). Data were analyzed with FlowJo software (version 10.8.1). Data were quantified as percentages of representative subpopulations, technical replicates (n=2-4) obtained from the same mice were used to calculate the mean for a biological replicate. Biological replicates, corresponding to individual mice, were applied for statistical analysis and displayed on the figures (Nude normal brain: n= 6; PDOX P8 n=6 (n=2-5 for activation markers); PDOX P3 n=3, PDOX P13 n=3)

### CD11b^+^ myeloid cell isolation and functional assays

Mice were perfused with ice-cold PBS. Tumor tissue was dissected from mouse brains and dissociated with the MACS Neural Dissociation kit (Miltenyi Biotec). PDOX brains were dissected into separate zones when specified: tumor core (cellular tumor, including pseudopalisading/hypoxic zone if present) and distant zone (back left hemisphere, PDOXs P3 and P13).

*Ex vivo* migratory abilities were assessed using 8 μm pore size Boyden chambers (ThinCert cell culture inserts, Greiner), fitted into 24-well plates. Myeloid cells were enriched using CD11b^+^ beads (MACS Miltenyi Biotec). Experiments were conducted in cells isolated from 3 distinct mice per model (biological replicates), each with 2 technical replicates (2 wells/ mouse). A total of 100,000 cells were seeded in the upper chambers in DMEM-F12 medium. After 48 hours, the cells were fixed in 4% PFA for 15 minutes and stained with DAPI for 15 minutes. Migratory cells were quantified by counting the number of cells on the lower side of the membrane under a light microscope with a 20x magnifying objective (5 fields/membrane). The data were normalized according to the proliferation index and are represented as the percentage of cells that migrated relative to the initial number of cells. Each biological replicate was calculated as a mean of technical replicates (number of cells per field of view followed by mean number of cells/view in technical replicates).

*Ex vivo* phagocytic abilities were measured using pHrodo Red E.coli bioparticles (Essen Bioscience, MI USA). Biological replicates represent 3 (Nu-NB, PDOX P3, PDOX P8) or 4 (PDOX P3) mice per model. A total of 100,000 freshly isolated CD11b^+^ cells were plated into 96 well plates in 100 μl and left for 2 h to adhere. pHrodo Red E.coli bioparticles were added at 10 μg/ml, and the plates were transferred into the IncuCyte ZOOM (Essen Bioscience, MI USA) platform. 4 images/well from at least 3 technical replicates (3 wells/mouse) were taken every hour for a duration of 44 h. The red fluorescence signal was quantified by applying a mask, and the parameter red object area was extracted for data analysis and visualization. Each biological replicate represents a mean of technical replicates. For flow cytometry, Percoll-purified freshly isolated myeloid cells were incubated with 1 µg of pHrodo™ Red E. coli BioParticles™ Conjugate for Phagocytosis (Invitrogen) at 37°C for 1 h followed by multicolor cell membrane marker staining at 4°C. Data acquisition was performed on a NovoCyte Quanteon Flow Cytometer (Agilent) fitted with a 405nm Violet laser, a 488 nm blue laser, a 561 nm yellow/green laser and a 637 nm red laser.

### Statistical analyses

Statistical analysis details for each experiment are reported in respective material and methods section and in the figure legends. For non-RNAseq data we applied parametric tests: (i) Students’ t-test (unpaired, two-tailed) for comparing two distinct groups; a Bonferroni multiple-significance-test correction for the number of conditions was applied to compare the proportions (ii) One-way Analysis of Variance (Anova) with Tukey’s Honest Significant Difference (HSD) correction for multiple comparisons for scenarios where we compared more than two biological groups. We employed parametric tests based on their high sensitivity, which helps mitigate the potential for false negatives that arise when non-normally distributed values are subjected to parametric tests. All data points for biological replicates are represented by dots, error bars represent standard error.

## RESULTS

### Human TME components are depleted in GBM PDOXs upon *in vivo* passaging

To characterize the recapitulation of TME in preclinical models *in vivo* we studied PDOXs derived from aggressive high grade gliomas. Up to date, we have derived a cohort of 46 PDOX models by intracranial implantation of patient-derived organoids (**Fig 1A, Table S1**). Our protocol is based on short term cultures of patient tumor tissue fragments that form 3D organoids *ex vivo*^16^. Organoids are further implanted intracranially to form orthotopic xenografts in the brain of immunodeficient mice. While use of NOD/SCID and NSG mice allows for higher engraftment rate, well-established models can be recapitulated also in nude mice with less immunodeficient background. We have previously shown that this protocol allows for an efficient recapitulation of histopathological and molecular features of patient tumors (**Fig S1A-B)**. Our current cohort comprises 42 IDH wild-type GBMs and 4 IDH mutant high grade astrocytomas. In depth assessment of genetic and DNA methylation features in 39 models revealed diverse GBM profiles regularly observed in patients (**Fig 1B, Table S1**). While human tumor cells can be expanded *in vivo* by serial passaging, we hypothesized that human components of the TME are depleted upon orthotopic xenografting, and are replaced by the equivalent mouse counterparts. As expected, flow cytometry confirmed depletion of these TME components already in the first PDOX generation (**Fig 1C**). These results were consistent with our prior profiling of human and mouse cells in PDOXs^18,19,22^. They also underscore the challenges associated with preserving long-term human TME components within preclinical models.

**Figure 1.**
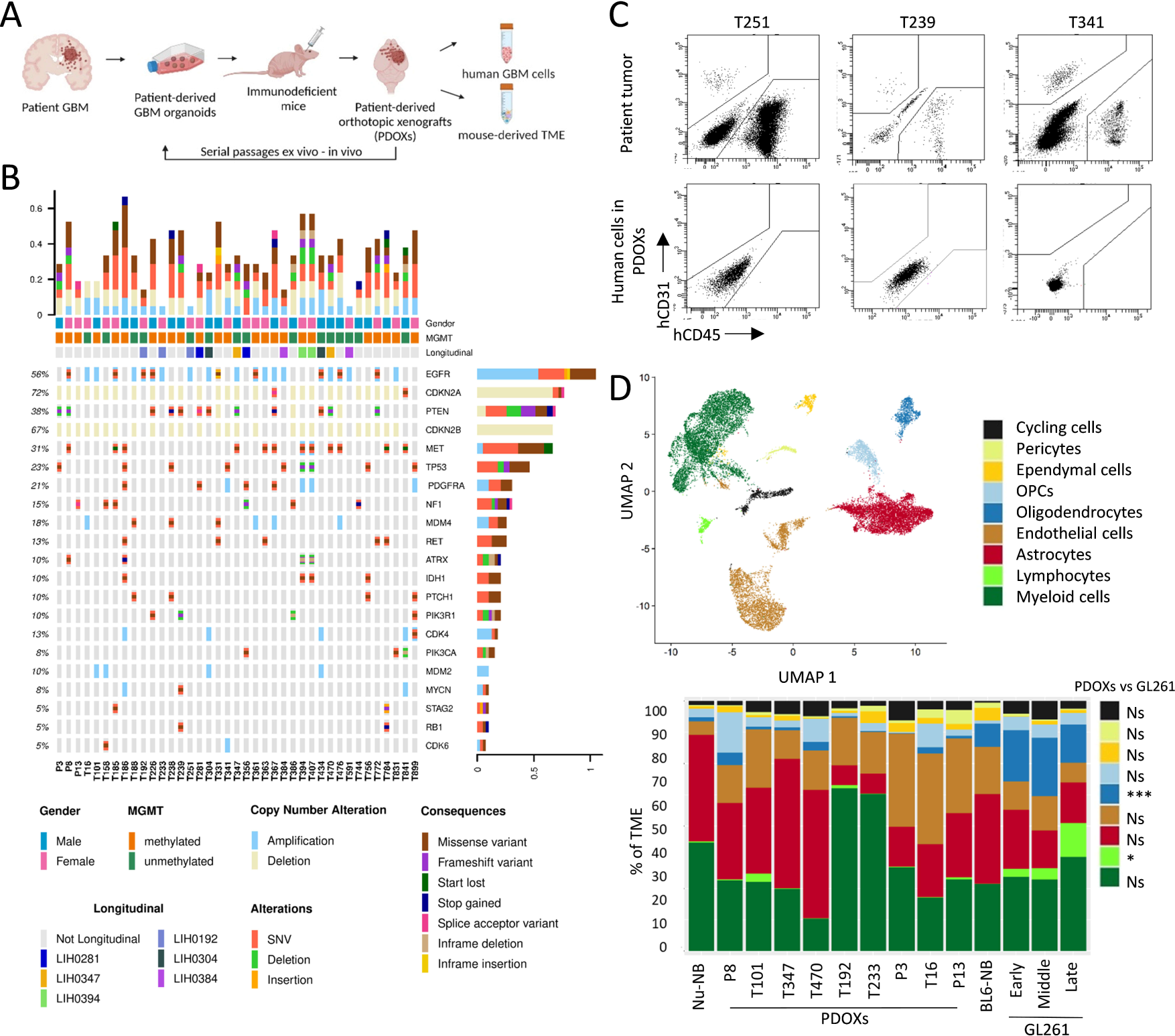
Composition of the mouse-derived TME in GBM PDOXs. **(A)** Schematic of the preclinical modeling of GBM tumors in PDOXs. See PDOX characteristics in **Fig. S1** and **Table S1. (B)** Oncoplot of glioma specific somatic mutations, gene amplifications and deep deletions in the PDOX cohort. Longitudinal PDOXs are highlighted with color. *MGMT* promoter methylation status in PDOX models in depicted. **(C)** Flow cytometric analysis showing depletion of human CD31+ endothelial cells and CD45^+^ immune cells upon xenografting. Examples are shown for 3 GBM patient tumors (single viable cells) and respective PDOXs models at the first passage (single viable human cells, characterized as GFP^neg^ population in GFP^+^ NOD/SCID mice). **(D)** Top: UMAP projection of scRNA-seq data showing the overall gene expression profile of TME cell types. scRNA-seq data combined the biological groups: nude mouse normal brain (Nu-NB), PDOXs (9 models), C57BL6/N mouse normal brain (BL6-NB), GL261 tumor (3 collection time points: early, middle, late). Bottom: Proportions of TME cell types across different tumors and normal brains; Statistical difference between PDOXs (n=9) and GL261 (n=3) was evaluated with two-tailed Student’s t-test with Bonferroni correction (***p<0.001, *p<0.1); Cell types are color-coded; OPCs: oligodendrocyte progenitor cells.

### scRNA-seq analyses identify major TME components in GBM PDOXs

Due to the depletion of human TME upon xenografting, we hypothesized that growth of human GBM cells in mouse brains is supported by diverse cell types of mouse origin forming the TME in GBM PDOX tumors. To assess TME composition in an unbiased manner, we performed scRNA-seq on mouse-derived cells. PDOXs were derived by intracranial implantation of GBM organoids into nude mice, which have the least immunocompromised background compared to NOD/SCID and NSG strains. We selected nine genetically and phenotypically diverse models, which were derived from treatment-naïve and recurrent IDH wild-type GBMs (**Fig 1B, Table S1, Fig S1A**)^18^. Four models represented longitudinal tumors derived from GBM patients prior and after standard-of-care treatment, which included radiotherapy and TMZ (LIH0347: T347/T470, LIH0192: T192/T233)^18^. All tumors were collected at endpoint, when tumors were fully developed. Tumor tissue were microdissected, following the previously evaluated MRI and histopathological features of each model, to ensure minimal contamination of healthy mouse brain. Mouse-derived cells of the TME were purified and processed by Drop-seq. In total we obtained 15,366 cells from nine PDOXs. The data were combined with the Pires-Afonso et al. dataset^24^ of the TME of the GL261 syngeneic orthotopic GBM mouse model derived in C57BL6/N (BL6/N) wild-type mice (3 time points, 2,492 cells in total). Normal brain controls were included for both mouse strains (1,692 cells/nude brain, 1,972 cells/BL6/N brain). Unsupervised clustering and uniform manifold approximation and projection (UMAP) analysis based on 21,522 cells and 24,067 genes in total, revealed nine major cellular clusters present in all samples (**Fig 1D, Table S3**). Cell clusters were identified based on the expression of cell type-specific markers (**Fig S2A**) and included well-known components of the normal brain and GBM TME, such as astrocytes, endothelial and ependymal cells, pericytes, oligodendrocytes and oligodendrocyte progenitor cells (OPCs), as well as immune cells **(Fig 1D**). All major cellular subpopulations were present in PDOXs, GL261 and normal brain controls. Similar to patients, myeloid cells constituted the major immune component in the PDOX and GL261 models. As expected, T lymphocytes were largely depleted and few functional B and NK cells were detected in PDOXs (**Fig S2B**), while the majority of infiltrated lymphocytes in the GL261 TME were T and NK cells, with increased proportions upon tumor development (3.3-13.5%, **Fig 1D**). GL261 also displayed higher proportions of oligodendrocytes (15-23%) than PDOXs (0.15-3.3%). In accordance with our previous report^19^, PDOXs with stronger angiogenic features (P13, T16, P3) had higher proportions of endothelial cells than more invasive PDOXs (**Fig 1D**). No correlation between histopathological features and the abundance of myeloid cells was observed (**Fig 1D**, **Fig S1B**).

The TME composition exhibited a patient-specific trend, e.g., longitudinal models (LIH0347: T347/T470, LIH0192: T192/T233) showed similar cellular proportions, where PDOXs derived from the LIH0192 patient showed a high percentage of myeloid cells, whereas PDOXs of the LIH0347 patient were particularly abundant in astrocytes. This suggests a potential influence of the genetic background of tumor cells on the TME composition, as has been suggested in human GBMs^13^. Since mesenchymal GBMs were described to contain the highest proportions of myeloid cells, we examined transcriptomic heterogeneity of human GBM tumor cells in PDOXs. PDOXs with high myeloid content did not show an increased abundance of mesenchymal-like GBM tumor cells (**Fig S2C**). We also did not observe major differences in the TME composition between PDOXs derived from treatment-naïve and recurrent GBMs. This can be explained by the fact that these models show similar genetic profiles at recurrence (**Table S1**)^18^ and do not display transcriptomic evolution towards the mesenchymal subtype (**Fig S2C**).

We further assessed the ontogeny of cells with active cell cycle programs. Assessment of the expression of key cell cycle marker genes (e.g. *Top2a*, *Mki67*, **Fig S2D**) revealed that cells with an active cell cycle gene expression program were in general clustered together in the ‘cycling cells’ cluster, with an exception of cycling myeloid cells that were also identified in the original ‘myeloid cells’ cluster. The ‘cycling cells’ cluster was identified in all conditions ranging from 0.5-1.6% for normal brains and 1.7-7.5% for PDOXs and GL261 tumors (**Fig 1D**) and was composed of different cellular entities (**Fig S2E**). Importantly, cells exhibiting active cell cycle gene expression profiles constituted a minor proportion within the cells in each cell type (**Fig S2F**). The highest proportion of cycling cells was observed for pericytes (28%), ependymal cells (19.5%) and oligodendrocytes (14.5%). Cycling myeloid cells were investigated in the follow up analyses. In summary, the mouse-derived TME in PDOXs is composed of cellular types relevant to human GBM^4^.

### TME subpopulations in PDOXs show transcriptional adaptation toward GBM-specific phenotypic states

As all relevant cell types were detected in the TME, we further tested to what extent murine TME cells in PDOXs are instructed by human GBM cells and acquire GBM-specific molecular profiles. To investigate the transcriptomic changes of each cell type, we compared identified cell population in GBM tumors with the corresponding cells in the naïve nude mouse brain. We detected pronounced transcriptomic differences across all the populations of the TME that were generally stronger than the changes observed in GL261 tumors when compared to normal BL6/N brain (**Fig 2A, Fig S3A**), thus indicating effective crosstalk between human GBM tumor cells and mouse-derived TME. The more prominent changes may result from the different tumor tissue structures between PDOXs and GL261. While PDOXs well recapitulate the cellular tumor niche with large areas of tumor-TME crosstalk, GL261 creates circumscribed tumors mostly representing the angiogenic niche with limited infiltration of brain-derived components of the TME, which remain high in the surrounding normal brain structures (**Fig S1A**). Transcriptomic adaptation was most pronounced in myeloid cells, endothelial cells, astrocytes, and OPCs (**Table S4**). Myeloid cells in PDOXs displayed transcriptomic programs linked to cell migration, inflammation and cytokine production (**Fig 2B**). Furthermore, key “homeostatic” Mg genes including *P2ry12*, *Tmem119* and *Gpr34*^47,48^ were downregulated (**Fig 2B**). In parallel, myeloid cells overexpressed GBM-specific TAM markers such as *Spp1* (Osteopontin), *Fn1*, *Cst7* and *Ch25h* pointing toward reciprocal crosstalk with GBM cells and transition to TAMs. We confirmed myeloid cell adaptation across the PDOX and GL261 models by qPCR of FACS-sorted CD11b^+^ cells (**Fig. S3B**).

**Figure 2.**
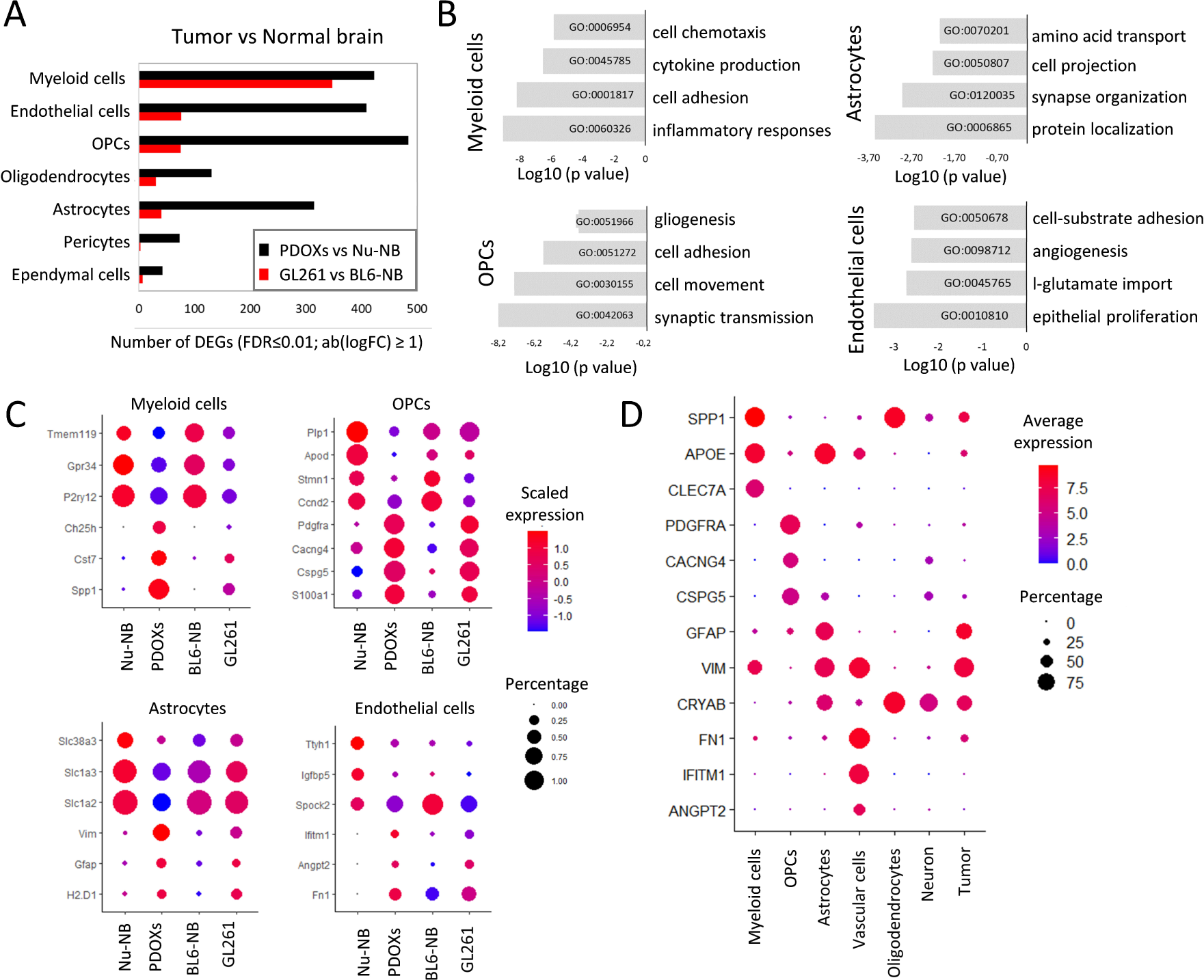
Transcriptomic adaptation of GBM-educated TME subpopulations in PDOXs. **(A)** Differentially expressed genes (DEGs) between the TME of PDOX and GL261 versus corresponding normal brains in identified cell types (FDR ≤0.01, |log_2_FC| ≥1, Wilcoxon rank sum test with Benjamini-Hochberg correction). **(B)** Top four gene ontology terms characterizing DEGs in PDOX versus Nu-NB. **(C)** Gene expression levels of exemplary DEGs for distinct cell types in four biological groups: nude mouse brain (Nu-NB), PDOXs (9 models combined), C57BL6/N mouse normal brain (BL6-NB), GL261 tumor (3 time points combined). **(D)** Expression levels of exemplary markers in distinct cell types detected and annotated in human GBM tumors by Darmanis et al.,^34^.

Interestingly, other cell types within the PDOX TME also activated biological processes linked to phenotypic states in GBM. For example, OPCs activated programs of tissue inflammation and regeneration (e.g., *Pdgfra, Cspg4, Cspg5, Cacng4,* **Fig 2B-C, Fig S3C**), while astrocytes expressed genes linked to metabolic processes and cellular shape, suggesting ongoing reactive gliosis (e.g., upregulated *Gfap* and *Vim*, downregulated *Slc1a2* and *Slc1a3,* **Fig 2B-C, Fig S3D**). In agreement with our previous study^19^, endothelial cells displayed an activated and proliferative phenotype associated with angiogenesis (**Fig 2B-C, Fig S1B**). We detected similar profiles in corresponding TME subpopulations of human GBM tumors (n = 3589 cells from 4 IDHwt GBMs^34^, **Fig 2D**). Altogether, these results point toward GBM-specific transcriptomic adaptations of myeloid cells and the main TME components in PDOXs.

### GBM-educated myeloid cells in PDOXs are largely of microglial origin

The abundance of myeloid cells was further confirmed by Iba1^+^ staining in PDOX tumors at the endpoint of tumor growth (**Fig 3A, Fig S1B**). While normal brains of nude mice showed Iba1^+^ myeloid cells with morphology corresponding to ‘surveilling’ ramified Mg, GBM tumors in PDOXs displayed TAMs with different morphologies. The cellular tumor was in general occupied by Iba1^+^ TAMs showing amoeboid or hyper-ramified morphology. Tumors displaying less invasive growth showed a gradient of TAM phenotypes, from ramified and hyper-ramified phenotypes at the invasive front toward amoeboid TAMs in the tumor center. Myeloid cells with macrophagocytic morphology were especially present in areas of pseudopalisading necrosis (PDOXs P13, T16). In these models we observed a notable accumulation of TAMs at the tumor border. A pronounced accumulation of TAMs was also detectable at a much sharper delineated GL261 tumor border (**Fig 3A**). Tumor showing invasive growth without a distinct tumor border showed more uniform, diffuse infiltration and activation of myeloid cells toward amoeboid states. Notably, certain resting and hyper-ramified morphologies remained detectable, particularly at the invasive front, in areas marked by lower tumor cells density (e.g. PDOXs P8 **Fig 3A**; PDOXs T101, T192 **Fig S1B**).This was in agreement with increased presence of amoeboid Iba1^+^ cells concomitant with increased tumor cell density upon tumor growth over time in mice (**Fig 3B**).

**Figure 3.**
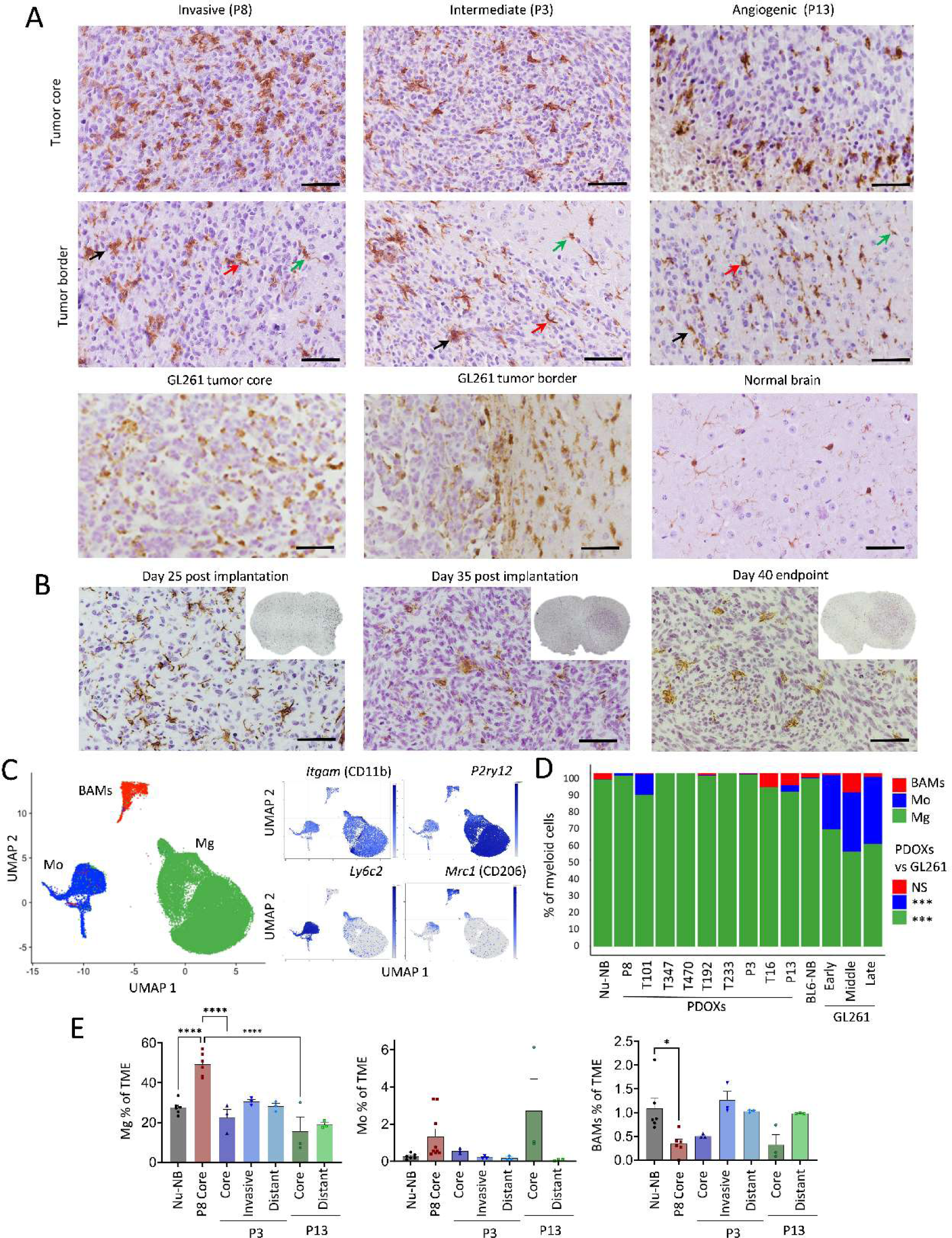
Ontogeny of GBM-educated myeloid cells. **(A)** Representative Iba1 staining in PDOXs depicting myeloid cells in invasive (PDOX P8), intermediate (PDOX P3) and angiogenic (PDOX P13) tumor growth, normal nude brain and GL261 tumor. Tumor core and tumor border zones are highlighted. Arrows indicate examples of ramified (green), hyper-ramified (red) and amoeboid (black) myeloid cells. Scale bar: 50 µm. Sections were co-stained with hematoxylin. See more examples in **Fig S1B**. **(B)** Representative Iba1 staining representing myeloid cells in PDOX P3 at different stages of tumor growth. Inserts represent sections of the entire mouse brains. Sections were co-stained with hematoxylin to visualize tumor cell density. **(C)** UMAP projection of reference-based mapping of myeloid cells from TME of GBM PDOXs and GL261 tumors and respective normal brains. Three myeloid cell entities were identified: microglia (Mg), peripheral monocyte-derived cells (Mo) and border-associated macrophages (BAMs). Inserts show expression of marker genes: pan-myeloid: *Itgam* (CD11b), Mg: *P2ry12*, Mo: *Ly6c2*, BAMs: *Mrc1* (CD206). The color gradient represents expression levels. **(D)** Proportions of myeloid cell subpopulations in nude mouse normal brain (Nu-NB), PDOXs (9 models), C57BL6/N mouse normal brain (BL6-NB), GL261 tumors (3 collection time points: early, middle, late). Statistical difference between PDOXs (n=9) and GL261 (n=3) was evaluated with two-tailed Student’s t-test with Bonferroni correction (***p<0.001). **(E)** Flow cytometry-based quantification of CD45^+^CD11b^+^Ly6G^−^Ly6C^−^CD206^−^ Mg, CD45^+^CD11b^+^Ly6G^−^Ly6C^+^CD206^−^ Mo and CD45^+^CD11b^+^Ly6G^−^Ly6C^−^CD206^+^ BAMs in the Nu-NB and PDOX P3, P8 and P13 TME. For PDOXs P3 and P13 invasive zone and distant normal brain areas were also collected (n≥3 mice/condition, mean ± SEM, one-way ANOVA with Tukey’s HSD correction, ****p<0.0001, *p<0.05). See gating strategy in **Fig S4D.**

Due to pronounced heterogeneity within the TME, we hypothesized that myeloid cells in PDOXs are composed of cells of different ontogeny and phenotypic states. To examine the ontogeny of myeloid cells, we combined our dataset with previously published scRNA-seq data of myeloid cells in GL261 tumors, which assigned the ontogeny of TAMs to Mg, Mo and BAMs based on transcriptomic profiles^7,8,24^ (**Fig 3C-D, Fig S4A-C**). Referencing PDOX myeloid cells to Ochocka et al.^7^ dataset confirmed a high abundance of Mg (81-100%) and a low proportion of Mo (0-19%) and BAMs (0-8%). P13 PDOX with pronounced angiogenic features showed a higher proportion of Mo and BAMs (7% and 5% respectively), although rare Mo were also present in invasive T101 PDOX. In contrast, GL261 tumors contained notably more Mo (**Fig 3D**). Flow cytometry confirmed high proportions of Mg in the PDOX TME compared to Mo and BAMs (**Fig 3E, Fig S4D-E**). While BAM proportions showed a trend towards decreased proportions in the tumor cores, we observed higher levels, although not significant, of Mo in the tumor cores compared to other brain areas in PDOXs and normal brain. It was accompanied by increased levels of neutrophils and lymphocytes in tumor cores of PDOX P13 (**Fig S4F)** suggesting that peripheral immune cell infiltration In PDOXs is very low and is limited to the tumor regions with highly disrupted blood-brain barrier.

Of note, we revealed major differences across GL261 datasets, with Ochocka et al.^7^ and Pires-Afonso et al.^24^ datasets containing 26-32% Mo, whereas Pombo-Antunes et al.^8^ dataset carrying more than 78% Mo, according to the different adopted cell isolation strategies (**Fig S4C**). In fact, within the first two studies, myeloid cells were isolated also at early stages of tumor development from a larger part of the tumor-containing hemisphere, whereas in Pombo-Antunes et al., cells were extracted at the endpoint and specifically from the tumor center. This highlights that Mg/Mo proportions depend on the sampling approach, suggesting diverse spatial locations in the tumor and adjacent brain regions.

### Mg-derived TAMs display heterogeneous transcriptional programs

We further hypothesized that cells of Mg, Mo and BAMs origin adapt their transcriptome toward GBM-specific states in PDOXs. To interrogate the phenotypic heterogeneity of myeloid cells in normal brain and tumors of different histopathological features, we next took advantage of our unique dataset containing normal brain and TME from PDOXs and GL261. To avoid batch effects arising from different scRNA-seq technologies, we performed reference-free analysis of our in-house Drop-seq dataset on the myeloid compartment, containing Mg, Mo and BAMs. The analysis stratified myeloid cells into nine phenotypic states: Mg formed seven phenotypic clusters (CL0-6), whereas CL7 and CL8 displayed transcriptional profiles of Mo and BAMs, respectively (**Fig 4A-C, Table S5**). PDOXs showed pronounced transitions toward heterogeneous Mg states, although with variable proportions (**Fig 4B**). Homeostatic Mg (Ho-Mg), highly enriched in the normal brain, grouped into two clusters (CL0-1), with CL1 showing lower expression levels of homeostatic genes (e.g. *P2ry12, Tmem119, Gpr34,* **Fig 4C**). Importantly, this was not the result of Mg activation via enzymatic digestion, since the markers of enzymatically-activated Mg (e.g., *Erg1*, *Fos*^49^) were expressed by a subset of Ho-Mg in CL0 (**Fig S5A**). Five phenotypic states were observed to be enriched in Mg-derived TAMs (Mg-TAMs, CL2-6) **(Fig 4A-C, Table S5**). These included classical pro-tumorigenic Mg-TAMs, which were present at the highest levels in CL2 and CL3, and were high for e.g. *Spp1, Cst7, Cxcl13 and Apoe* (**Fig 4C)**. Among these two groups, CL3 presented higher cytokine expression levels (*Ccl3, Ccl4*), suggesting stronger secretory properties and education by GBM when compared with CL2. Subset of CL3 cells showed also high transcriptional activity (e.g. *Rpl17*, *Rps12,* **Fig S5A**). As expected, CL3 Mg-TAMs showed higher *Ptprc* (CD45) expression and lower levels of homeostatic Mg genes (**Fig S5A**). This is reminiscent of the decrease of homeostatic genes in reactive Mg, known to occur in the GBM TME, but also under inflammatory and neurodegenerative conditions^50–53^. Mg transition toward CD45^high^ and CCR2^+^ TAM states in the tumor core was further detected by flow cytometry (**Fig 4D**). While Mg in distant brain areas (PDOXs P3, P13) resembled normal brain characteristics, the invasive niche (PDOX P3) showed partial activation of Mg toward Mg-TAMs. Additional subpopulations included Mg-TAMs displaying astrocytic features (CL4, e.g. *Sparcl1, Gfap*), and expression of endothelial cell markers (CL5, e.g., *Pecam1, Cldn5*) at similar levels than the original astrocyte and endothelial cell clusters (**Fig S5A)**. We excluded contamination by other TME subpopulations within these clusters as these cells expressed myeloid cell markers, including *Hexb*, *Csf1r*, *Itgam* (CD11b) and *Ptprc* (**Fig S5A**) and showed low scores for potential doublets (**Fig S5B**). CL4 and CL5 cells also displayed expression of pro-tumorigenic markers, but at more variable levels than CL3 Mg-TAMs. However, we cannot exclude the possibility of detection of the mRNA from astrocytic and endothelial cells due to potential phagocytosis of these cells by Mg-TAMs. We also identified a separate Mg-TAM cluster corresponding to cells with activated cell cycle programs (CL6). Presence of GFAP^+^ astrocytic-like, CD31^+^ endothelial-like and Ki67^+^ cycling Iba1^+^ myeloid cells was confirmed by immunohistochemistry (**Fig S5B**).

**Figure 4.**
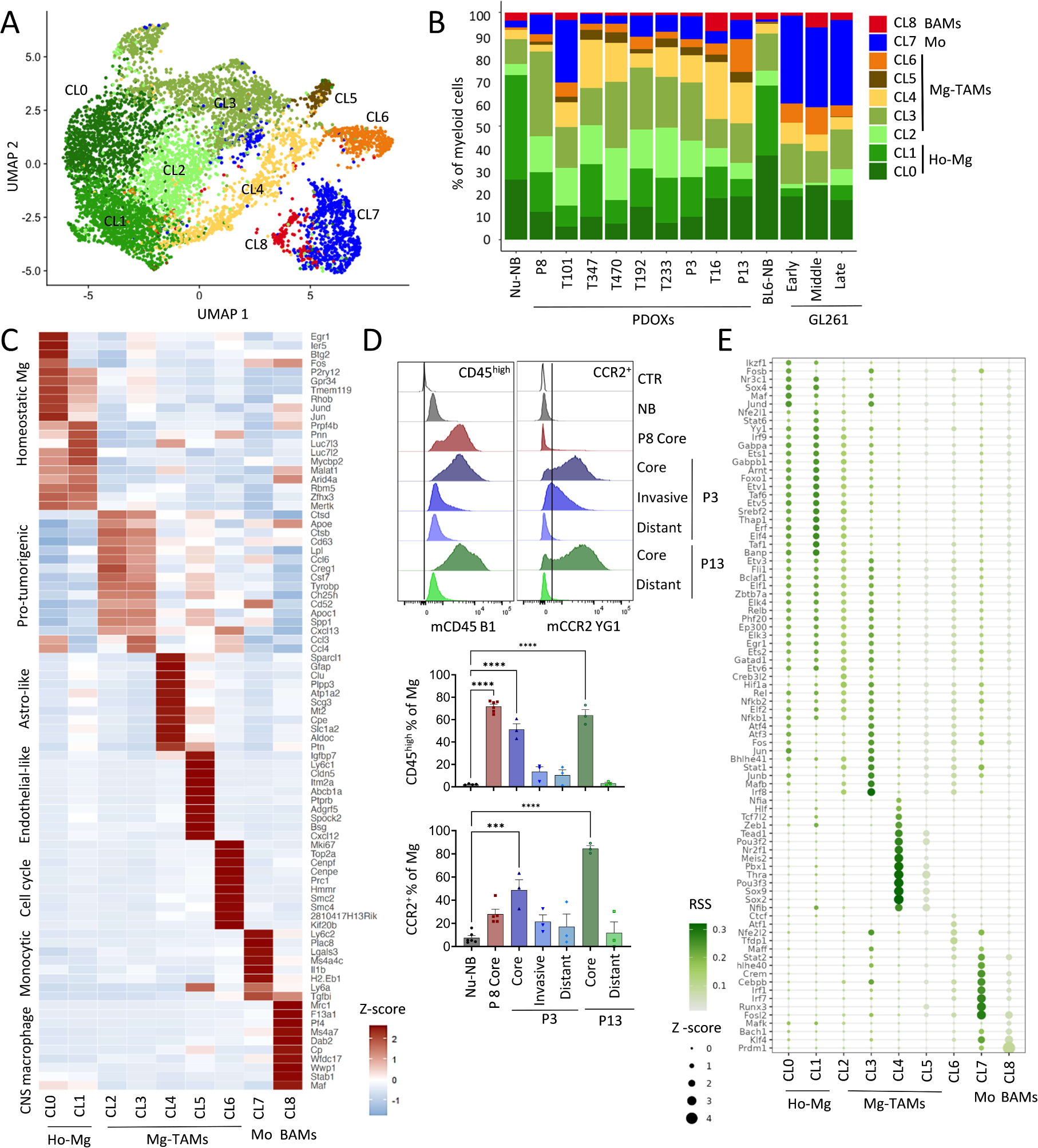
GBM-driven activation of Ho-Mg toward heterogeneous Mg-TAMs. **(A)** UMAP plot showing clusters of myeloid cells in PDOXs, GL261 and normal brain controls revealing nine distinct clusters (CL). CL0-6 represent microglia (Mg), CL7 peripheral monocyte-derived cells (Mo), CL8 border-associated macrophages (BAMs). **(B)** Proportions of cells assigned to nine clusters of myeloid cells in nude mouse normal brain (Nu-NB), PDOXs (9 models), C57BL6/N mouse normal brain (BL6-NB), GL261 tumor (3 collection time points: early, middle late). CL0-1: homeostatic-Mg (Ho-Mg), CL2-6: Mg-derived tumor-associated macrophages (Mg-TAMs), CL7: Mo, CL8: BAMs. **(C)** Discriminative marker genes for each myeloid state (row z-scores of the expression levels). **(D)** Representative flow cytometry graphs and quantification of CD45^high^ and CCR2^+^ cells in CD45^+^CD11b^+^Ly6G^−^Ly6C^−^CD206^−^ Mg in PDOX P3, P8 and P13. For PDOXs P3 and P13 invasive zone and distant normal brain areas were also collected (n≥3 mice/condition, mean ± SEM, one-way ANOVA with Tukey’s HSD correction, ****p<0.0001, ***p<0.001). **(E)** Relative transcription factor (TF) activity of regulons identified by SCENIC in myeloid clusters. Regulons with RSS<0.05 are shown.

To understand the interdependence between identified Mg states, we further performed the trajectory analysis of Mg cells. TSCAN-based trajectory analysis revealed a general transition from Ho-Mg to Mg-TAMs via CL2, with more profound differences found for CL5 endothelial-like and CL6 cycling Mg-TAMs (**Fig S5D-E**). To further examine the transcriptomic differences between identified myeloid states, we next sought to reveal transcriptomic regulators by conducting SCENIC analyses^30^ (**Fig 4E**). We identified a high number of regulons for Ho-Mg states including Maf, Nr3c1 and Sox4. While CL2 Mg-TAMs showed again transitory features between Ho-TAMs and CL3, CL3 Mg-TAMs appeared regulated by transcription factors such as Hif1a, Stat1, Mafb and Irf8 ^54^ suggesting a role of hypoxia in Mg state transitions towards protumorigenic states. Astrocytic-like Mg-TAMs (CL4) showed high activity for Thra, Sox9 and Sox2, which are known to regulate astrocytic states, while cycling Mg-TAMs (CL6) regulons were enriched in classical cell cycle regulators (e.g., Tfdp1, Atf1). Importantly, although certain regulatory networks were shared between Mg-TAM states, Mo and BAMs, we also detected specific regulons unique for Mo (e.g., Fosl2, Irf1, Cebpb) or shared between Mo and BAMs (e.g. Bach1, Prdm1 and Klf4). These data further highlight the factual differences between transcriptomic states of Mg and show the impact of TME niches in shaping Mg heterogeneity in GBM.

### Rare Mo and BAMs undergo phenotypic adaptation toward TAM features in GBM TME in PDOXs

We further aimed to investigate whether Mo and BAMs undergo phenotypic adaptation toward TAMs in GBM tumors developed in PDOXs. Since both cell types were rare in PDOXs, we reverted to the reference-based analysis, which allowed us to discriminate between three Mo states described initially by Ochocka et al.^7^ (**Fig S4A-C**, **Fig S6A**). In 5 out of 9 PDOXs we did not detect Mo. Similarly to GL261 tumors, the majority of Mo detected in the remaining 4 PDOXs (P8, T101, T192, P13) transitioned to Mo-TAMs displaying in intermediate monocyte/macrophage (inMoM) or differentiated macrophage states (**Fig S6B**). Compared to Mo detected in normal brains, Mo detected in PDOXs showed higher expression levels of genes associated with pro-tumorigenic TAMs (e.g., *Cxcl13, Ctsd, Ccl4, Apoe*) as well as Mg mimicry (e.g., *Spp1, Trem2, Cst7, Tyrobp,* **Table S6**), further confirming an active cross-talk with tumor cells. Compared to the normal brain, BAMs detected in PDOXs also showed transcriptional convergence toward Mg by upregulating expression levels of Mg genes (e.g., *Clec7a*) and especially downregulating several BAM markers (e.g. *Cd163*, *F13a1*, *Mrc1*, **Table S6**). Further data will be needed to robustly determine activation of genes in Mo-TAMs and BAM-TAMs, since the amount of cells detected in our cohort was low. At the cell membrane level, while Mo show high CD45 levels already in the normal brain, Mo-TAMs in PDOXs, similarly to Mg-TAMs, increased CCR2 levels in the tumor core (**Fig S6C**). BAMs were high for CD45 and CCR2 both in the normal brain and PDOX tumors.

Despite the differences in proportions, the transcriptomic profiles of Mg, Mo and BAMs were generally similar between PDOXs and GL261 tumors (**Fig S6D)**. We detected differences in Mg at the level of activation of several genes, e.g. higher levels of *Cxcl13, Spp1*, *Apoe,* and *Itgax*, while lower levels of *H2.Eb1, Ccl12,* and *Cxcl4* in PDOXs compared to GL261 tumors (**Fig S6E, Table S7**). Mo in PDOXs showed higher levels of *Hexb, Siglech* and *Cst3*, suggesting stronger priming toward Mg mimicry, whereas Mo in GL261 presented higher levels of *Tgfbi, Thbs1* and *Vegfa*, in line with the higher angiogenic features of the TME in GL261 tumors. Interestingly, both Mo and BAMs displayed higher levels of *Arg1, Tgfbi* and *Il1b* in GL261 than PDOXs (**Table S7**), suggesting a prominent pro-tumorigenic TAM profile in the angiogenic niche.

In summary, although the mobilization of Mo and BAMs in PDOXs is very limited, we detected that an active crosstalk with GBM cells is possible for the rare cells that reach the tumor site. The extent of the activation status towards pro-tumorigenic Mg-TAMs and Mo-TAMs may depend on the model applied and their underlying histopathological features.

### Mg-TAMs display immune-reactive states with increased capacity for chemotactic, phagocytic and dendritic cell-like properties

We next investigated the functional properties of myeloid subpopulations. At the global level, gene ontology analysis of genes activated in myeloid cells in PDOXs uncovered enrichment of terms associated with cell chemotaxis, adhesion and migration, and tumor-associated extracellular matrix proteins (**Fig 2B**). These terms may also reflect changes in the phagocytic and antigen-presenting capabilities of these cells, since cell chemotaxis and migration by degradation of the extracellular matrix are processes required for detecting and tracking tumor cells for engulfing them via phagocytosis, the latter being a critical event for subsequent antigen-presentation. Therefore, we focused further analysis on the gene signatures involved in migration, sensome, phagocytosis and antigen presentation (**Table S7, Fig 5A)**.

**Figure 5.**
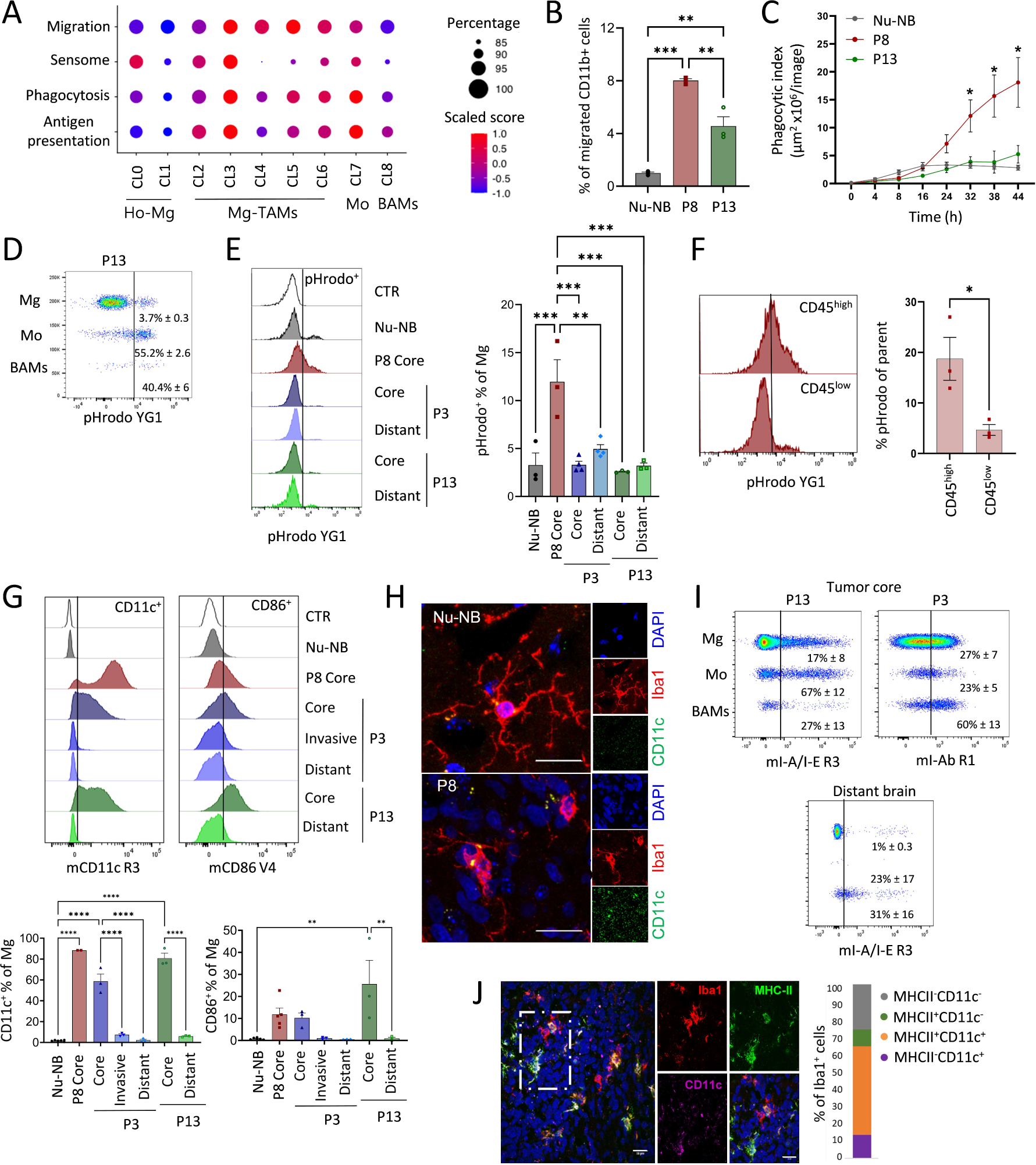
Functional properties of GBM-educated Mg. **(A)** Signature score of genes associated with migration, sensome, phagocytosis and antigen presentation per myeloid cluster: CL0-1 Ho-Mg, CL2-6: Mg-TAMs, CL7: Mo, CL8: BAMs. **(B)** *Ex vivo* assessment of migratory capacity in CD11b^+^ myeloid cells isolated from nude mouse normal brain (Nu-NB) and PDOXs P8, P13 (n=3 mice/condition, mean ± SEM, **p < 0.01, ***p < 0.001, one-way ANOVA with Tukey’s HSD correction). **(C)** *Ex vivo* assessment of phagocytic capacity in CD11b^+^ myeloid cells isolated from Nu-NB and PDOXs P8, P13 (n=3-4 mice/condition, mean ± SEM, *p<0.05, one-way ANOVA with Tukey’s HSD correction). (**D**) Phagocytic uptake measured in CD45^+^CD11b^+^Ly6G^−^Ly6C^−^CD206^−^ Mg, CD45^+^CD11b^+^Ly6G^−^Ly6C^+^CD206^−^ Mo and CD45^+^CD11b^+^Ly6G^−^Ly6C^−^CD206^+^ BAMs. Representative example is shown for PDOX P13 tumor core (mean ± SEM, n=3). (**E**) Phagocytic uptake measured in Mg in Nu-NB and PDOXs P8, P3, P13 (n=3 mice/condition, ***p<0.001, **p<0.01, mean ± SEM, one-way ANOVA with Tukey’s HSD correction). An unstained control (CTR) represents Mg without E. coli particles incubation. For PDOXs P3 and P13 tumor core and distant normal brain area were collected. (**F**) Comparison of phagocytic uptake in CD45^high^ and CD45^low^ Mg in PDOX P8 (n=3 mice, mean ± SEM, *p<0.05, two-tailed Student’s t-test). **(G)** Representative flow cytometry graphs and quantification of Mg positive for CD11c and CD86 in Nu-NB and PDOXs P3, P8, P13 across different brain regions (n≥3 mice/condition each, mean ± SEM, ****p<0.0001, **p<0.01, one-way ANOVA with Tukey’s HSD correction). An unstained control is shown for each population (CTR). **(H)** Representative immunofluorescence images depicting CD11c staining in Iba1^+^ cells in the Nu-NB and PDOX P8 tumor core. Scale bar: 20 µm. **(I)** Representative flow cytometry graphs showing activation of MHC-II (I-A/I-E epitope) in Mg, Mo and BAMs in the tumor core and distant brain. Examples shown for PDOX P13 and P3 (mean ± SEM, n=3 mice). **(J)** Representative immunofluorescence images depicting MHC-II and CD11c co-expression of Iba1^+^ cells in the P8 PDOX tumor core. Quantification represents mean values from 2 technical replicates. Scale bar: 20 µm.

A migration score based on genes associated with monocyte, glial and neutrophil cell migration (e.g. *Fn1, Cxcl13, Ccl3*) inferred increased migratory capacity during transition from Ho-Mg toward different phenotypic states of Mg-TAMs (**Fig 5A, Fig S7A**), while signatures in Mo and BAMs were more moderate. We functionally verified the increased migratory ability of CD11b^+^ myeloid cells freshly isolated from PDOXs (P8, P13) in comparison to naïve brains using Boyden chambers (**Fig 5B**).

We further investigated genes related to the Mg sensome, which reflect main Mg functions in the brain as danger sensing cells. Mg sensome genes (**Table S7**) showed a higher score in CL0, CL2 and CL3 Mg-TAMs compared to other Mg subsets, Mo and BAMs (**Fig 5A**). Importantly, in CL2 and CL3, the sensome signature was driven by genes, such as *Cd74*, *Cd52, Cxcl16* and *Clec7a*, and not homeostatic Mg genes, which were highest in CL0 Ho-Mg (**Fig S7A**), indicating that CL2 and CL3 Mg-TAMs turn on a specific activated sensome program concomitantly decreasing the expression levels of the classical homeostatic gene markers. Interestingly, astrocytic-like (CL4), endothelial-like (CL5) and cycling (CL6) Mg-TAMs showed low scores for the sensome signature, suggesting that while they downregulate the classical homeostatic features, they do not acquire the sensome functions detected at higher levels in CL2 and CL3 (**Fig 5A**).

Mg-TAMs also showed increased expression levels of genes related to phagocytosis (e.g. *Trem2, Tyrobp, Axl)* and antigen presentation (*Itgax* (CD11c), *Igf1, Cd86, H2.Eb1, H2.Ab1 (*I-A/I-E)), with the highest signature levels present in CL3 Mg-TAMs, similar to Mo but not BAMs **(Fig. 5A, Fig S7A**). These signatures showed a strong correlation with each other (**Fig S7B**). We further functionally evaluated the phagocytic capacity of freshly isolated CD11b^+^ myeloid cells in PDOXs with different proportions of CL3 Mg-TAMs (37% PDOX P8, 26% PDOX P3 and 17% PDOX P13) and in normal brain (11% CL3 of myeloid cells) using E. coli particles labeled with fluorescent pH-sensitive dye^55^. Indeed, CD11b^+^ cells isolated from PDOX P8 showed increased phagocytic abilities compared to CD11b^+^ cells isolated from normal brains and PDOX P13 (**Fig 5C**). Within CD11b^+^ cells, the high proportion of rare Mo (55.2% ± 2.6) and BAMs (40.4% ± 6) presented phagocytic capacity (**Fig 5D**). We also observed active phagocytosis in Mg, which was highest in PDOX P8 (12% ± 1.3) with the highest proportion of CL3 Mg-TAMs (**Fig 5D-E**). The majority of phagocytic cells were detected within CD45^high^ Mg-TAMs **(Fig 5F**), further linking this function to CL3. We also confirmed prominent CD11c and CD86 activation at Mg cell membrane, mainly in the tumor core of three PDOXs confirming antigen presentation cell (APC)-like features (**Fig 5G**). Increased expression levels of CD11c were also detected by immunohistochemistry in amoeboid Iba1^+^ cells in the cellular tumor, but not in ramified Mg in the normal brain (**Fig. 5H**). CD11c and CD86 were also expressed by Mo and BAMs, consistent with the gene expression profiles (**Fig S7C)**. We further confirmed activation of MHC-II expression (*H2.Eb1, H2.Ab1* and corresponding epitope I-A/I-E, **Fig 5I-J, Fig S7A**) in subsets of Mg-TAMs and Mo compared to normal brains. Subsets of Mg-TAMs and Mo also expressed checkpoint inhibitors, such as *Cd274* (PD-L1), *Havcr2* (TIM-3) and *Pdcd1* (PD-1, **Fig S7A**), which are known to inhibit the phagocytic capacity of macrophages. Mg-TAMs also showed increased levels of *Sirpa* and *CD47*, maintaining the ‘’do-not-eat-me’’ signal. We did not detect increased expression levels of genes typically associated with macrophage immune suppression (e.g. *Arg1*, *Mrc1, Il10*) in Mg-TAMs (**Fig S5A**). Interestingly, astrocytic-like (CL4), endothelial-like (CL5) and cycling Mg-TAMs (CL6) showed less prominent phagocytic and antigen presentation scores than other Mg-TAMs (CL2, CL3). Further investigations will be needed to understand the balance between activating and inhibitory signatures of these subpopulations at the functional level and their localization within the various tumor niches. Overall, these results suggest that specific subpopulations of Mg-TAMs display phagocytic and dendritic cell-like programs under tumorigenic conditions.

### Heterogeneous Mg states represent central components of GBM patient tumors

To investigate the relevance of our findings in PDOXs, we next probed the composition of myeloid cells in publicly available scRNA-seq datasets of *IDH* wild-type GBM patient tumors by extracting the myeloid compartment from the GBmap, a curated scRNA-seq database of 240 patient tumors^38^, and by our own analysis based on published datasets^8,35–37^. Due to the lack of normal human brain Mg and human blood Mo references, we applied robust gene signatures to assign myeloid cell ontogeny (**Fig 6A, Fig S8A-B, Table S7)**. Mg constituted the main myeloid cell subset across the majority of GBMs. As reported previously^8^, recurrent GBMs generally showed increased proportions of Mo, albeit with high interpatient differences. BAM signatures were rather weak and were not clearly segregated within a specific cluster (**Fig S8A)**. The analysis of gene signatures identified in preclinical models in human myeloid cells confirmed the presence of heterogeneous Mg and Mo subsets in human GBMs (**Fig 6B**). We detected a subset of Mg with high levels of signatures corresponding to Ho-Mg (CL0, CL1) as well as a subset of Mo with high scores for signatures corresponding to naive Mo. This confirms our analysis in PDOX models, suggesting activation of protumorigenic features in the close proximity of the tumor cells. CL2 and CL3 Mg-TAM signatures were robustly expressed by a subset of human Mg (**Fig 6B, Fig S8C)**. Although CL4, CL5 and CL6 signals appeared in general to be weaker than classical pro-tumorigenic signatures, subsets of Mg showed high scores for astrocytic-like (CL4), endothelial-like (CL5) and cell cycle (CL6) Mg-TAM features (**Fig S8C**). We further confirmed the adaptation of Mo to Mo-TAMs, which showed decreased levels of Mo markers concomitant with increased levels of Mg genes. Mo-TAMs appeared also to share Mg-TAM signatures and contained a subset of proliferating cells. Such bilateral convergence of Mg and Mo was further confirmed in bulk RNA-seq profiles of Mg and Mo cells isolated from GBM patient tumors based on the CD49d expression levels^42^ (**Fig S8D**). Similar to mouse myeloid cells, both Mg-TAMs and Mo-TAMs showed activation of migratory, phagocytic and antigen presentation features (**Fig 6B)**. We further confirmed the relevance of our findings in PDOXs by mapping mouse myeloid cells to the GBmap^38^ datasets corroborating the presence of Mg and Mo and their convergence toward tumor-specific programs (**Fig. S8E)**. Altogether, these data show the relevance of distinct Mg states and the robustness of their signatures identified in PDOXs in GBM patient tumors.

**Figure 6.**
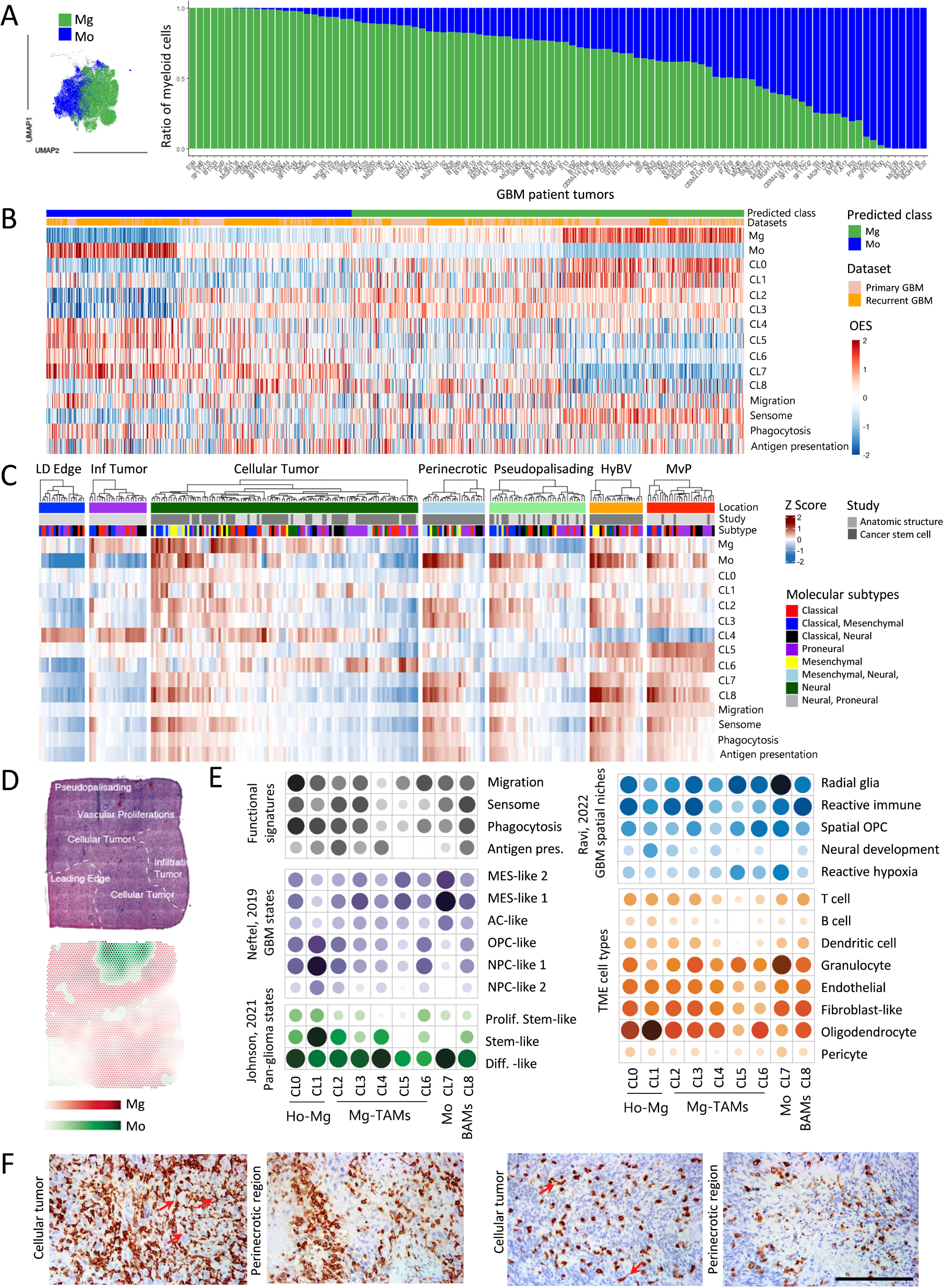
Transcriptomic states of myeloid cells in human GBM. **(A)** UMAP projection of myeloid subsets from GBM patient tumors (103 patient tumors, GBmap datasets^38^). Cells are color coded for Mg and Mo ontogeny based on established gene signatures. Proportions of Mg and Mo are shown in individual GBM patient tumors. (**B**) Convergence of Mg and Mo to TAMs in GBM patient tumors. Heatmap showing overexpression scores (OES) of signatures in scRNA-seq primary (n=7) and recurrent (n=4) GBMs^8^. Each column represents a single cell. **(C)** Heatmap showing OES of signatures in different tumor locations from the IVY GAP bulk RNA-seq data: LD edge: leading edge; Inf tumor: infiltrative tumor; HyBV: hyperplastic blood vessels in cellular tumor; MvP: microvascular proliferation. Each column represents a microdissected tissue fragment (n=279 from 44 patient tumors). (**D**) Representative surface plots of the spatial localization of Mg and Mo signatures in GBM patient tumors. (**E**) Spatially weighted correlation analysis of the enrichment scores in myeloid transcriptomic states (CL0-8) linked to functional myeloid signatures, GBM tumor states, spatial TME niches and TME cell components (n= 16 tumors from 16 individual patients). (**F**) Representative Iba1 stainings in GBM patient tumors. Cellular tumor and perinecrotic regions are shown for 2 patients. Red arrows depict Iba1^+^ myeloid cells with visible ramifications, Scale bar: 100µm

### Heterogeneous Mg and Mo in GBM patients are spatially distributed across different tumor niches

To investigate the spatial distribution of GBM-associated myeloid states, we assessed myeloid signatures in the bulk RNA-seq of GBM tumor niches (Ivy Glioblastoma Atlas Project dataset^39^, **Fig 6C, Fig S9A**). In general, Mg signatures were evident in infiltrative and cellular tumors, hyperplastic blood vessels (HyBV) and microvascular proliferation (MvP) zones, although CL2 and CL3 Mg-TAMs also scored high in perinecrotic and pseudopalisading areas. Astrocytic-like CL4 and endothelial-like CL5 Mg-TAMs showed high signals in leading edge/infiltrating/cellular tumor and HyBV/MvP niches, respectively. We hypothesize that this could be related to a closer interaction with reactive astrocytes (leading edge) or endothelial cells (HyBV/MvP), which are abundant at these locations. Cycling CL6 Mg-TAM signatures showed high levels in cellular tumor, HyBV and MyP niches. Importantly, these signatures should be interpreted with caution as they are likely biased by the bulk RNA-seq signal, where discrimination of signals from different TME cell types is not possible. Mo (CL7) and BAM (CL8) signatures were particularly evident in niches displaying blood-brain barrier leakage, including HyBV, MvP, perinecrotic and pseudopalisading areas, consistent with our findings in PDOXs and GL261. Migration, sensome, phagocytic and antigen presentation signatures were expressed in the cellular tumor, perinecrotic zone, hyperplastic blood vessels and microvascular proliferation zones, and were particularly high in tumors with a high abundance of Mg and Mo. These functional signatures were less pronounced in the leading edge and infiltrative zone, confirming the education of Mg in close proximity to GBM cells. Interestingly, these signatures were relatively lower in pseudopalisading tumor zones regardless of the high abundance of Mo-TAMs, suggesting a potential role of severe hypoxia in the inhibition of myeloid cell functions in this niche. The high score for Mg/Mo signatures was particularly evident for tumors with mesenchymal components, confirming previous studies^12^.

We further confirmed the differential distribution of myeloid states in spatially resolved transcriptomic profiles of GBM patient tumors^45^. Again, Mg were highly abundant in the infiltrating and cellular tumor, whereas Mo were enriched in the pseudopalisading and vascular proliferation areas (**Fig 6D, Fig S9B-C**). Spatially weighted correlation analysis and spatial gene set enrichment analysis confirmed the co-localization of CL0-1 Ho-Mg (CL0-1), CL2-3 Mg-TAM, Mo and BAM signatures with signatures of sensome, phagocytosis and antigen presentation (**Fig 6E**). CL3 Mg-TAMs and Mo co-localized with MES-like states, corresponding to the “Differentiated-like pan-glioma state”. While CL0 Ho-Mg were detected in close proximity to different GBM states, CL1 Ho-Mg were most abundant in close proximity to OPC-like and NPC-like GBM states, equivalent to the “Stem-like pan-glioma states”. This was corroborated by the distribution across spatial GBM niches. All myeloid cells were present in different GBM niches with high tumor content (Radial glia, Reactive immune, Spatial OPC niches). Ho-Mg (CL1) were particularly abundant in the ‘Neural development’ niche at the tumor edge, whereas Mo were associated with ‘Reactive hypoxia’ niches, further confirming Mo localization in the area of high necrosis, BBB leakage and hypoxia. This was different for CL3 Mg-TAMs, which appeared present at higher levels in the spatially distinct ‘Reactive immune’ niche, corresponding to regions with high glial signatures enriched for inflammation-associated genes and non-hypoxic MES-like GBM states. Indeed, we detected distinct morphological features of Iba1^+^ myeloid cells across different niches in GBM patient tumors. Specifically, we identified cells with visible ramifications, reminiscent of naïve microglia, together with amoeboid cells in cellular tumor niches, while the perinecrotic niche predominantly contained amoeboid cells (**Fig 6F**). Interestingly, analysis of bulk RNA-seq CGGA patient tumor profiles^43^ revealed shorter survival of patients with high scores for Mg-TAM CL2 and CL3, but not other clusters in classical GBMs (**Fig S9D**). No statistical difference in survival was observed for proneural and mesenchymal tumors. Taken together, these data confirm the transcriptomic heterogeneity of Mg and Mo in GBM patient tumors associated with different functional features across specific tumor niches.

### TMZ treatment leads to transcriptomic adaptation of GBM cells and adjacent TME

The molecular adaptations of individual cells within GBM tumors upon treatment are incompletely understood^56^. We therefore aimed to assess the adaptation of tumor cells and the TME upon treatment, hypothesizing that chemotherapy can lead to transcriptomic adaptation of the entire GBM ecosystem. For this, we administered TMZ to P3 PDOXs representing *MGMT* promoter-methylated GBM (**Fig S10A**). Tumor growth was validated by MRI, and mice were treated 5 times a week at the clinically relevant TMZ dose for 10 days (total 8 doses received). Experiment was halted for all animals when the control animals showed neurological symptoms and fully developed tumors. Tumors were resected shortly after the last TMZ dose. MRI-based quantification of tumor volumes confirmed that prolonged TMZ treatment led to decreased tumor growth (**Fig 7A-B**).

**Figure 7.**
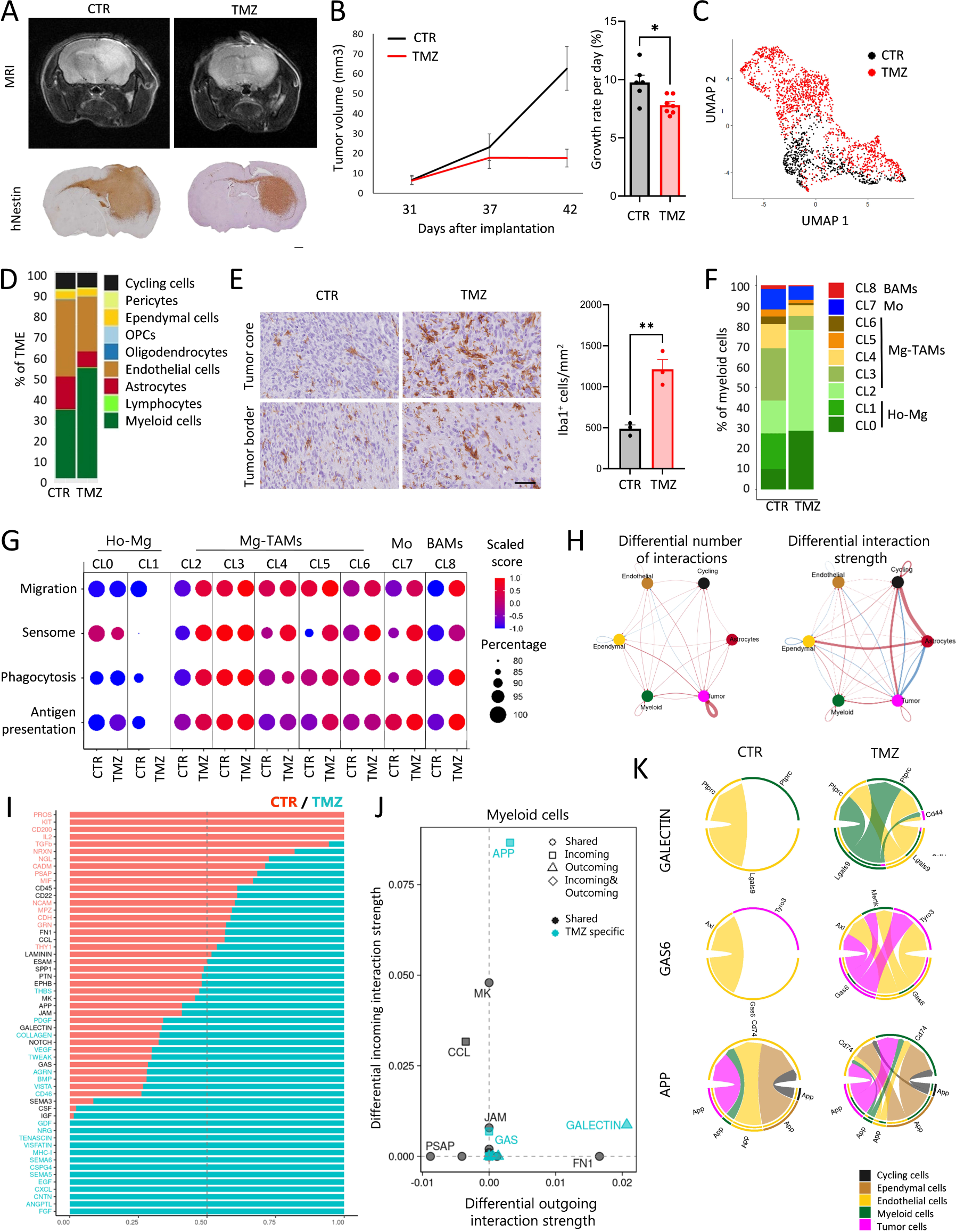
Transcriptomic adaptation of GBM tumor cells and TME upon TMZ treatment. **(A)** Representative MRI images and human-specific Nestin staining showing tumor growth in control PDOXs P3 (CTR) and TMZ-treated mice (TMZ). Scale bar = 1 mm. **(B)** MRI-based assessment of tumor progression over time. The tumor growth rate was calculated during the entire study (day 42 vs day 31, n = 6-7 mice/group, *p<0.05, two-tailed Student’s t-test). **(C)** UMAP projection showing the overall gene expression relationship between TMZ-treated and CTR GBM tumor cells (n=1 mouse/condition). **(D)** Proportions of cell types in TME in CTR and TMZ-treated PDOX inferred from scRNA-seq (n=1 mouse/condition). **(E)** Representative images of Iba1^+^ cells in in CTR and TMZ-treated PDOXs. Tumor cores and borders are highlighted. Scale bar: 50 µm. Quantification of Iba1^+^ cells in tumor core is depicted (n= 3 mice/condition; **p<0.01, two-tailed Student’s t-test) **(F)** Proportions of cells assigned to nine clusters of myeloid cells in CTR and TMZ-treated PDOXs inferred from scRNA-seq (n=1 mouse/condition); CL0-1: Ho-Mg, CL2-6: Mg-TAMs, CL7: Mo, CL8: BAMs. (**G**) Functional signature score per myeloid cluster in CTR and TMZ conditions. (**H**) CellChat-based differential number of interactions and interaction strength of the inferred cell–cell communication networks between cell types in CTR and TMZ PDOXs (threshold > 10 cells/sample). Red or blue colored edges represent increased or decreased signaling in TMZ-treated tumors. (**I**) Relative information flow from cell–cell interaction analysis. Receptor-ligand pathways with blue text are significantly enriched in TMZ-treated cells, and pathways with red text are significantly enriched in CTR cells. (**J**) Signaling changes in myeloid cells in TMZ vs CTR conditions. (**K**) Interactions related to the Galectin, Gas6 and App pathways under CTR and TMZ conditions.

To assess transcriptomic changes in tumor and TME populations, we extracted tumor tissue from control and TMZ treated mice and purified single human tumor cells and single mouse cells. ScRNA-seq analysis of isolated tumor cells revealed transcriptomic changes linked to survival mechanisms such as regulation of p53-associated signal transduction, apoptosis, cell death and cellular component organization (**Fig 7C, Fig S10B-C**), suggesting activation of resistance mechanisms in surviving tumor cells. Assessment of GBM cellular subtypes revealed an increased proportion of MES-like states in line with observations made in GBM patients^11,12^. Corresponding scRNA-seq analysis of the TME revealed changes in the proportions of cell populations (**Fig 7D**). We observed an increased ratio of myeloid cells upon TMZ treatment and a relative decrease in ECs and astrocytes. Indeed, TMZ-treated tumors contained more Iba1^+^ myeloid cells in the tumor core (**Fig 7E**). The analysis of DEGs between TMZ-treated and control tumors revealed transcriptomic changes in myeloid cells and ECs, but not in astrocytes (**Table S8, Fig S10D**). Upon treatment, myeloid cells enhanced the expression levels of genes associated with inflammatory responses, such as migration, chemotaxis, and gliogenesis (e.g., *Cxcl13, Cx3Cr1, Csf1r*). Adaptation was visible at the level of regulation of translation (e.g., *Rps15*, *Rpl32*, *Rpl23*), endocytosis (e.g., *Apoe, Lrp1*), cholesterol homeostasis (e.g., *Abca1, Abcg1*) and actin cytoskeleton (e.g*., Fscn1, Coro1a*). This correlated with decreased levels of TAM markers promoting tumor growth, such as *Igfbp7* and *Gng5*. Although the heterogeneity of the myeloid compartment shifted to a higher ratio of CL2 Mg-TAMs (**Fig 7F**), the activation of functional signatures was observed in the majority of Mg-TAM clusters, Mo and BAMs (**Fig 7G**). In parallel, ECs deregulated genes associated with cell development and death, extracellular space, regulation of chemokine and cytokine production and acetylcholine receptor activity (e.g., up: *Ly6a/c1/e, H2.D1, Timp3, Cxcl12; down: Ctsd, Ctss*) as well as regulation of actin cytoskeleton and protein localization (**Fig S10D**).

Since transcriptomic adaptation was detected in tumor and TME cells we hypothesized an overall adaptation of the molecular crosstalk within the GBM ecosystem. Indeed CellChat analyses revealed significant changes in ligand-receptor signaling pathways between cell populations in CTR and TMZ-treated tumors, with increased number of inferred interactions (236 CTR vs 425 TMZ) and interaction strength (5.717 CTR vs 6.415 TMZ) in TMZ treated tumors (**Fig 7H-I**). Among those, putative ligand-receptor interactions affected in myeloid cells upon TMZ treatment showed increased outgoing interactions associated with and increased incoming interactions associated with Gas6 (Growth arrest-specific 6) and App (Amyloid-beta precursor protein) pathways (**Fig 7J, Fig S10E-F**). Upon TMZ treatment, Galectin signaling was linked to Lgals9 produced by myeloid cells predicted to interact with Cd44 in tumor cells and Ptprc (Cd45) in ependymal and myeloid cells. Gas6 produced by tumor and ependymal cells was predicted to interact with the receptor Mertk in myeloid cells, Tyro3 in tumor cells and Axl in ependymal cells, while additional App-Cd74 signaling axes were predicted between tumor, myeloid, endothelial and ependymal cells (**Fig 7K)**. Taken together, we showed that induction of the cell death resistance mechanism in tumor cells upon TMZ treatment is associated with transcriptomic changes in TME components, including inflammatory responses of Mg-TAMs and activation of the ‘’eat-me’’ Gas6 pathway.

## DISCUSSION

A lack of thorough understanding of the TME and its recapitulation in preclinical models is regularly listed as one of the main challenges for the discovery of effective treatments against brain tumors^15^. Therefore, the cellular and molecular understanding of preclinical models is of utmost importance to avoid failure of clinical trials and has been repeatedly pointed out as a key need in the community^15^. Using unbiased scRNA-seq analysis, we surveyed the TME of nine GBM PDOXs and compared it with the TME of the GL261 mouse glioma model and human GBM tumors. To our knowledge, this is the first in-depth analysis of the TME in PDOX models. We show that GBM PDOXs present diverse cell types in the TME similar to those reported in human GBM^12,34,35^. We provide evidence that human tumor cells instruct the TME subpopulations in PDOX models toward GBM-associated phenotypic states, thus unlocking their relevance as preclinical models to investigate the modulation of the GBM ecosystem upon treatment and testing novel therapies against tumor and TME cells. We further uncovered diverse cellular and molecular specificities of the GBM-associated myeloid compartment. Specifically, we found that resident Mg represent the main myeloid cell population in PDOXs and human GBMs, while peripheral-derived Mo mostly infiltrate the brain at sites of the blood-brain barrier disruption. Mg exhibit molecular plasticity toward diverse GBM-associated states reflecting intratumoral heterogeneity. Notably, we detected reactive dendritic cell-like gene expression programs in a large subset of GBM-educated Mg-TAMs. These cells show high activation of the Mg sensome followed by increased phagocytic and antigen presentation capacity. We show that Mg states are differentially distributed across spatial GBM niches, where they co-localize with varying TME components and GBM phenotypic states. Lastly, we highlight the adaptation of diverse TME components, including myeloid cells upon TMZ treatment, which leads to the differential crosstalk with GBM tumor cells.

Studies of the differences between TAMs of different origins have been confounded by a lack of specific markers to separately purify these cell types within GBMs. By applying robust transcriptomic gene signatures, we found that Mg subsets represent an important fraction of myeloid cells in PDOXs, syngeneic models and GBM patients, confirming reports in mouse chimeras^57^ and human GBMs^3,8,58^. Discrepancies in the field may arise from varying sampling strategies and marker selection. As Mg-TAMs downregulate classical homeostatic genes while activating macrophage markers, Mg/Mo discrimination based solely on the expression of selected homeostatic and/or macrophage markers may lead to biased misidentification of myeloid entities. We show that in human GBMs and preclinical models, Mg are abundant in the cellular tumor and invasive niche, whereas Mo are confined mostly to perinecrotic, and hypoxic areas with the leaky blood-brain barrier. Interestingly, PDOXs with more bulky growth show higher accumulation of Mg at the tumor border and more spatially distributed Mg heterogeneity than invasively grown tumors. This is in accordance with reports showing the diffuse presence of Mg in gliomas in contrast to brain metastases, where Mg appears to be often confined to the tumor border areas^3^. Mo are particularly abundant in the GL261 syngeneic model, which does not recapitulate the cellular tumor and infiltrative tumor zone well. Interestingly, GL261 show more invasive tumor growth in immunodeficient SCID mice concomitant with both less aberrant vasculature and more uniform distribution of TAMs^59^. It remains to be seen whether Mo are active players or a bypass product of the blood-brain barrier leakage and to what extend Mg infiltration influences the invasiveness of tumor cells to the brain. Although we confirmed a higher proportion of Mo in a subgroup of recurrent human GBMs^8,11^, we highlight high inter-patient differences and potential sampling bias linked to GBM niches and tissue isolation strategies, as observed in preclinical models. Analysis of the myeloid compartment in longitudinal PDOXs derived from matched primary and recurrent patient tumors did not reveal inherent changes in the TME composition without genetic and transcriptomic evolution of tumor cells. We rather detected patient-specific profiles that were retained in recurrent models. This is consistent with studies conducted in genetically engineered models of gliomas, where Mg/Mo ratios are model and not treatment dependent^60,61^. A larger cohort will be needed to further interrogate the correlation between TME composition and GBM molecular features. Interestingly, we observed decreased survival of patients with high CL3 Mg-TAMs for classical GBMs, but not mesenchymal and proneural GBMs. We previously showed that the TAM signature associated with Mo correlates with decreased survival in gliomas in general, though the difference is not retained in GBM patients only^24^. This is consistent with the co-localization to hypoxic and perinecrotic niches, two factors associated with poorer survival in gliomas.

Importantly, the observed low levels of Mo in PDOXs might not solely originate from the GBM tissue structure; there might be other contributing factors linked to modeling using immunodeficient mice. For instance, it has been documented that Mo in SCID and nude mice exhibit compromised maturation processes, resulting in diminished responsiveness to external stimuli^62^, which could explain restricted mobilization to the brain during GBM tumor growth in PDOXs. It is also essential to account for the potential influence of impaired communication between human GBM cells and mouse Mo within the PDOX system. Mo functionality depends on the specific cytokine milieu, which frequently exhibits species-specific differences^63^. Thus, the observed disparities in Mo levels could, in part, stem from the species-specific differences in cytokine signaling. Despite their distinct developmental origin and intrinsic transcriptional networks, myeloid cells are known to share signatures of tumor education, although specific functions of Mo- and Mg-TAMs have also been suggested^64,65^. The high abundance of Mg in our PDOXs allowed us to further discriminate distinct phenotypic states of Mg-TAMs, which is more challenging in the syngeneic models^7,24^. We show that Mg activation occurs in PDOXs generated in nude mice, which show strongly reduced and hypo-responsive T cells^66^, suggesting that myeloid-T cell crosstalk is not a prerequisite for TAM activation. Since nude mice still possess B and NK cells as their main lymphocytic subpopulations, it remains to be seen whether the loss of T cells is compensated by other available lymphocytes. Of note, in GBM patients naïve T cells are sequestered in the bone marrow, which contributes to a very low abundance of T cells in the tumor^67^. As a dominant myeloid population, Mg underwent pronounced activation in PDOXs, whereas Mo dominated GL261 tumors, where they acquired highly immunosuppressive states. These observations are in line with results obtained during GL261 progression, where early stages of tumor development are dominated by CD11c^+^ Mg invading the tumor, followed by recruitment of CD11c^+^ Mo-derived DCs^6^.

We show that Mg subpopulations range from homeostatic to GBM-educated states. We identified two states corresponding to Ho-Mg (CL0-1) and five Mg-TAM states (CL2-6). Importantly, these states were detected across PDOXs and patient GBMs with different genetic backgrounds, highlighting pan-GBM significance and intra-tumoral heterogeneity. Ho-Mg states are present within several GBM tumor niches representing cellular tumors and invasive edges, suggesting ongoing education in close proximity to tumor cells. While CL0 Ho-Mg was detected in several GBM spatial niches, CL1 Ho-Mg were particularly abundant in close proximity to OPC/NPC-like GBM states in ‘Neural development’ niches. While ramified Mg are still detectable in close proximity to the tumor cells, we showed accumulation of amoeboid Mg in the dense tumor areas upon tumor growth in PDOXs. Mg-TAM states share classical GBM education, including decreased levels of homeostatic genes, co-expression of pro- and anti-inflammatory molecules and increased levels of markers classically associated with pro-tumoral macrophages. Importantly, Mg states also display discrete transcriptomic features and gene regulatory networks. CL3 show particularly high features of the Mg sensome, but also phagocytosis and antigen presentation at similar levels to Mo, while CL2 appears as a transitory state from Ho-Mg to Mg-TAMs. Importantly, increased sensome activity is driven by a limited set of specific genes (*Cd74*, *Clec7a, Cxcl16*), while key homeostatic genes responsible for general sensing changes in the brain are downregulated, confirming the reduced capacity to sense changes in the TME caused by GBM. While all Mg states are detected across several spatial GBM niches, CL3 Mg-TAMs are particularly abundant in ‘Reactive immune’ niche co-localizing with MES-like GBM cells, whereas ‘Reactive hypoxia’ niche is predominantly enriched with CL7 Mo cells. While we confirmed transition of Mo and BAMs towards TAMs in PDOXs and patient GBMs, we did not further focus on the discrimination of Mo-TAM and BAM-TAM states due to scarcity of these cells in PDOXs. We functionally confirmed that phagocytosis is enhanced in the cellular tumor compared to the adjacent normal brain and can be supported by all myeloid cell entities. Our data are in line with previous reports showing the phagocytic activity of GBM-associated Mg^68,69^. Although the functional implications of phagocytic Mg in GBM are still elusive, emerging data suggest both pro- and anti-tumoral effects. For instance, phagocytic Mg were shown to populate necrotic tumor zones and aid in the clearance of debris to enhance GBM cell invasion^69^. Increased Mg phagocytosis was reported to enhance antigen cross-presentation toward more efficient T cell priming upon combined TMZ and anti-CD47 treatment^70^ as well as upon CTLA-4 blockage ^71^.

Dendritic-like Mg have also been reported in disease-associated microglia (DAM) in Alzheimer’s disease^72–74^, amyotrophic lateral sclerosis and multiple sclerosis^75^. Mg-TAMs and DAM appear to activate similar programs including decreased homeostatic genes, classical activation markers (e.g. *Spp1*, *Il1b*), phagocytic (e.g. *Apoe, Trem2*, *Tyrobp*) and APC (e.g. *Itgax* and *Igf1*) signatures. This suggests that phagocytic and APC-specific transcriptional programs are associated with Mg detecting damage within the CNS^53^ as well as recognizing and clearing pathogenic factors, such as neoplastic cells in GBM and β-amyloid aggregates in Alzheimer’s disease, but not along acute inflammatory processes, where Mg rapidly restore and maintain the homeostatic neuronal network^50^. Mg and brain resident macrophages are also known to act as competent APCs during CNS infections, and are potentially involved in the activation of infiltrating T cells^76^.

Intriguingly, we identified additional clusters within Mg-TAMs expressing astrocytic (CL4) and endothelial (CL5) markers as well as cycling Mg-TAMs (CL6). These phenotypic states, despite high scores for migration, showed lower levels of the sensome, phagocytosis and antigen presentation, suggesting other functions. As bulk RNA-seq and spatial transcriptomics do not present sufficient resolution to deconvolute these signals from bona fide reactive astrocytes and endothelium, their localization in the tumor needs further investigation in the future. The expression of astrocytic genes such as *Gfap* and *Serpina3n* has been previously reported in Mg in mouse models of injury and autoimmune encephalomyelitis where it is speculated that they might suppress pro-inflammatory pathways^77^. GFAP has also been detected in a subset of circulating Mo in brain tumor patients^78^.

Lastly, we demonstrate the utility of PDOX models in understanding adaptation of the GBM ecosystem upon treatment. Although several studies have suggested an increased abundance of TAMs at recurrence^8,11^, a direct analysis upon treatment is not possible in human tumors. By applying TMZ to a *MGMT*-methylated GBM PDOX model, we identified both tumor cell and TME remodeling. Increased apoptotic signaling and a pronounced MES-like GBM state were observed together with increased Mg migration toward the tumor. Mg-TAMs in treated tumors displayed re-polarization toward more pro-inflammatory states with increased phagocytosis and antigen presentation scores. We observed altered molecular crosstalk with tumor cells, e.g. through activation of Gas6 signaling across several cell types. Although further investigation of the key axes in such complex crosstalk is needed, we speculate that Gas6 expressed by apoptotic cells activates efferocytosis, a well-known function of Mg in the brain, aimed at the removal of damaged apoptotic cells, thus contributing to the resolution of inflammation^79^. We cannot exclude a direct impact of TMZ on Mg as cGAS-STING was recently reported to be activated upon damage of the Mg cells themselves, leading to inflammation^80^. Further studies with a larger cohort of treated PDOXs will be needed to fully understand the adaptation of GBM cells with different molecular backgrounds and their adjacent TME. As direct effects of the treatment on Mg have also been reported^81^, further studies are needed to dissect molecular events upon treatment leading to changes in the cellular network.

Research efforts aimed at deciphering the functional heterogeneity of TAMs may contribute to the development of novel immune therapeutic approaches in GBM patients. It remains to be investigated how TAMs can be reprogrammed against GBM cells^82^. TAMs showed enhanced phagocytic ability following CD47 blockade^68,83,84^, an effect that was more pronounced in combination with TMZ^70^. These results suggest that the phagocytic capacity of Mg-TAMs appears to be pervasive and may require fine-tuning in the context of therapeutic reprogramming. Further functional characterization is needed to harness their anti-tumor potential, for instance by addressing their capacity to recruit lymphocytes and their phagocytic ability against tumor cells. Taken together, we show key adaptation of brain resident cells to TME niches in GBM. Our work provides a novel understanding of the TME in GBM *in vivo* models. Such investigations are still scarce in the community, which has led to incorrect interpretation of the preclinical outcomes and failure of numerous clinical trials. Furthermore, we unravel heterogeneous Mg states, which reside in different spatial niches. In-depth characterization of specific signatures of the TME and their adaptation upon standard-of-care treatment will pave the way toward rational design of targeted treatment strategies. The use of PDOX avatars holds promise for the functional assessment of the plastic GBM ecosystem upon treatment and for testing novel therapeutics, including modalities targeting GBM-educated myeloid cells.

## CONCLUSIONS

Overall, we show that PDOX models faithfully recapitulate the major components of the GBM-educated TME and allow assessment of phenotypic changes in the GBM ecosystem upon treatment. Focusing on the myeloid compartment, we show the education of microglia and monocyte-derived cells toward GBM-specific heterogeneous states. We identify reactive dendritic cell-like gene expression programs associated with enhanced phagocytic and antigen-presentation features in GBM-educated microglia subsets that might be harnessed for novel immunotherapeutic approaches. Our data provide an important characterization of the TME in patient-derived models, which is key for tailoring future investigations of treatment responses and resistance mechanisms prior to clinical trials.

## ETHICS DECLARATIONS

### Ethics approval and consent to participate

All patients gave informed consent and tumor collection was approved by the local research ethics committees (National Committee for Ethics in Research (CNER) Luxembourg: 201201/06; Haukeland University Hospital: 2009/117). Animal experiments were performed in accordance with the regulations of the European Directive on animal experimentation (2010/63/EU) and were approved by the Animal Welfare Structure of the Luxembourg Institute of Health and by the Luxembourg Ministries of Agriculture and of Health (LRNO-2014-01, LUPA2019/93 and LRNO-2016-01).

### Consent for publication

Not applicable

### Availability of data and materials

PDOX models are available via PDCM finder (www.cancermodels.org) and are part of the EuroPDX consortium collection (www.europdx.eu). The scRNA-seq data from the PDOX models have been deposited in the Gene Expression Omnibus repository (www.ncbi.nlm.nih.gov/geo/) with accession numbers GSE226468 (mouse TME) and GSE128195 (human tumor cells). Codes used to analyze data in this paper are available at https://github.com/yahayayabo/GBM_TME

### Authors’ contributions

Conceptualization: YAY, SPN, AlM, AG; Methodology: YAY, SKP, AP, AS, SPN, DHH, AlM, AG; Investigation: YAY, PMS, YPA, KG, AL, SKP, AP, AS, AO, BK, MM, DHH, AlM, AG; Formal analysis: YAY, PMS, YPA, TK, DK, KG, BN, LE, SKP, AP, AS, AM, RT, DHH, PVN, AlM, AG; Resources: GB, CB, FH, MM, DHH, SPN, AlM, AG; Supervision: RT, FH, AP, AS, PVN, SPN, AlM, AG; Writing - Original Draft: YAY, AlM, AG; Writing - Review & Editing: all authors

## Acknowledgements

We are grateful to the Neurosurgery Department of the Centre Hospitalier de Luxembourg and Clinical and Epidemiological Investigation Center of the LIH for support in tumor collection. We thank Virginie Baus, Lorraine Richart and Jean-Jacques Gérardy for technical assistance. We acknowledge the excellent support from LIH’s core facilities (Animal facility, In vivo imaging facility, National cytometry platform, LUXGEN Genome Center) and sequencing platform at the Luxembourg Centre for Systems Biomedicine (LCSB) at the University of Luxembourg.

