## Supplementary Material and Figures for "Glioblastoma-instructed microglia transition to heterogeneous phenotypic states with phagocytic and dendritic cell-like features in patient tumors and patient-derived orthotopic xenografts"

[1]NORLUX Neuro-Oncology Laboratory, Department of Cancer Research, Luxembourg Institute of Health (LIH), L-1526 Luxembourg, Luxembourg; [2]Department of Life Sciences and Medicine, Faculty of Science, Technology and Medicine (FSTM), University of Luxembourg, L-4367 Belvaux, Luxembourg; [3]Neuro-Immunology Group, Department of Cancer Research, Luxembourg Institute of Health, L-1526 Luxembourg, Luxembourg; [4]Multiomics Data Science, Department of Cancer Research, Luxembourg Institute of Health, L-1445 Strassen, Luxembourg; [5]Luxembourg Centre for Systems Biomedicine (LCSB), University of Luxembourg, L-4362 Esch-sur-Alzette, Luxembourg; [6]Single Cell Analytics & Microfluidics Core, Vlaams Instituut voor Biotechnologie-KU Leuven, 3000 Leuven, Belgium; [7]National Center of Genetics, Laboratoire National de Santé, L-3555 Dudelange, Luxembourg; [8]Department of Cancer Research, Luxembourg Institute of Health, L-1526 Luxembourg, Luxembourg; [9]German Cancer Consortium (DKTK), 01307 Dresden, Germany; Core Unit for Molecular Tumor Diagnostics (CMTD), National Center for Tumor Diseases (NCT), 01307 Dresden, Germany; German Cancer Research Center (DKFZ), 69120 Heidelberg, Germany; [10]Institute for Clinical Genetics, Faculty of Medicine Carl Gustav Carus, Technische Universität Dresden, 01307 Dresden, Germany; [11]Centre Hospitalier Luxembourg, 1210 Luxembourg, Luxembourg; [12]Luxembourg Center of Neuropathology (LCNP), Luxembourg; [13]National Center of Pathology (NCP), Laboratoire National de Santé, L-3555 Dudelange, Luxembourg; [14]Microenvironment and Immunology Research Laboratory, Medical Center - University of Freiburg, Freiburg, Germany; [15]Department of Neurosurgery, Medical Center - University of Freiburg, Freiburg, Germany; [16]Department of Neurological Surgery, Lou and Jean Malnati Brain Tumor Institute, Robert H. Lurie Comprehensive Cancer Center, Feinberg School of Medicine, Northwestern University, Chicago, Illinois. [17]Department of Physics and Material Science, University Luxembourg, L-4367 Belvaux, Luxembourg; [18]Department of Neuroscience, University of California San Diego, La Jolla, CA 92093, USA

### SUPPLEMENTARY MATERIAL AND METHODS

#### Molecular profiling of PDOX models

Our current cohort comprises 46 glioma PDOX models. 39 PDOX models were characterized at the genetic and epigenetic levels. Targeted sequencing, copy number aberrations and DNA methylation profiling of PDOX models were performed as previously described[1] unless specified otherwise below.

#### Targeted DNA sequencing

500ng of extracted gDNA were diluted in 130µl low TE buffer (Qiagen) and sheared via sonication on a Bioruptor® UCD-200 (Diagenode) to an average fragment size of 150-300 bp. DNA fragment size was determined using the DNA 1000 Kit on the Bioanalyzer 2100 (Agilent Technologies). A custom-made Agilent SureSelect$^{XT}$ Target Enrichment Library (Cat No. G9612B) was used for Illumina Paired-End Multiplexed Sequencing on a MiSeq® instrument (Illumina). The panel design called panel 1 for the Target Enrichment Library was fully adapted from [2]. After design changes that were made using SureDesign - Agilent eArray (Agilent Technologies) we refer to it as panel 2 containing additional regions. Samples have been sequenced with version 1 of the panel or with version 2 of the panel (additional 53 genes). Library preparation was performed according to manufacturers' instructions. The Illumina MiSeq® Reagent Kit v3 (Cat No. MS-102-3003) was selected by applying the Illumina reagent selection algorithm.

Bioinformatics analysis of sequencing reads was performed using a Snakemake-based workflow[3], in accordance with GATK best practices. Briefly paired-end reads from each tumor were quality trimmed with fastp (v. 0.23.2)[4] and aligned to an *In silico* Combine human-mouse Reference Genome (ICRG) for mouse read cleaning using BWA mem (v. 0.7.17) [5] with default parameters. ICRG contained both the mouse (GRCm39) and human (Grch38/hg38) reference genomes. Reads mapping to human chromosomes were extracted from the resulting bam file using SAMtools (v. 1.17) and realigned only to the human reference genome with BWA mem. Duplicates were annotated with MarkDuplicates in GATK4 (v. 4.2.6.1). Base quality score recalibration was carried out in GATK4 using BaseRecalibrator and ApplyBQSR. Mosdepth was used to evaluate bam statistics[6]. MuTect2 (GATK4) was used in a tumor-only mode to identify somatic single nucleotide variants (SNVs) and short insertions and/or deletions (indels). To distinguish between germline and somatic variants, a hg38 GATK Panel of Normals was used. LearnReadOrientationModel (GATK4) was used to evaluate read orientation artefacts, while GetPileupSummaries (GATK4) and CalculateContamination (GATK4) were adopted to determine the proportion of contaminated reads across samples. The following criteria were used to identify variants: a minimum base quality score of 25, a minimum allele frequency of 5%, and a minimum 10X coverage. Variants caused by cross-sample contamination and read orientation artefacts were eliminated. The Ensembl Variant Effect Predictor (VEP)[7] was used to annotate somatic variants. Only variants identified by VEP as having a high or moderate impact were kept. Variants occurring in non-glioma specific genes were filtered out. Glioma specific genes: *IDH1, ATRX, TP53, CDKN2A, CDKN2B, EGFR, CDK4, CDK6, MDM2, MDM4, MET, MYCN, PDGFRA, NF1, PIK3CA, PIK3R1, PTCH1, PTEN, RB1, RET, STAG2*. Multiple genomic alterations were visualised with ComplexHeatmap[8] within R version 4.2.0.

#### DNA methylation

Methylation arrays with Infinium® MethylationEPIC were analysed as previously described[1] to establish the *MGMT* methylation and glioma classification by referencing data to the neuropathological tumors dataset at https://www.molecularneuropathol ogy.org/mnp.

#### 10XGenomics single-cell RNA-sequencing and analysis

Human tumor cells were processed via Drop-seq (P3, P8, P13, T101, T16) or 10XGenomics (T192, T233, T347, T470). Drop-seq and data preprocessing were performed as previously described[1,9]. For the 10XGenomics, purified single tumor cells were diluted to $10^6$ cells/ ml with RNAse free DPBS + 0.05 % FBS. Diluted cells were filtered using a 50µm filter. The viability of cells was verified using Beckman coulter automatic cell counter and C-Chip – Disposable Haemocytometer (NanoEntek) viability analyzer. Cell concentration was adjusted to target the encapsulation of 3000 cells according to 10XGenomics cell preparation guidelines. Chromium Next GEM Chip with single-cell suspension was loaded to the Chromium Controller (10XGenomics) for generation of encapsulated cells and GEMs. scRNA-seq libraries were prepared using the Chromium Next GEM Single Cell 3' GEM, Library & Gel Bead Kit v3.1 and a Chromium i7 Multiplex kit according to the manufacturer's protocol. Libraries were purified using SPRIselect magnetic beads and analyzed using Agilent 2100 Bioanalyzer. Sequencing was performed using NextSeq 500/550 High Output Kit v2.5 (75 Cycles). scRNA-seq analysis was performed in R (v4.1.1) with the Seurat package (v4.0.5)[10]. Barcode processing and UMI filtering were performed using the Cell Ranger 4.0.0. Average saturation of the sequencing was 30-40% as calculated by Cell Ranger. GBM molecular subtypes (Classical, Proneural and Mesenchymal) and cellular states (MES, AC, NPC and OPC-like) were determined using signatures listed in Wang et al., 2017[11] and Neftel et al., 2019[12] respectively. Single-cell signature scores were calculated using scrabble(v1.0.0) used in Neftel et al., 2019[12].

**Gene expression analysis by qPCR**

Myelin was removed prior FACS-sorting with the Myelin removal beads II (Miltenyi) and CD11b$^+$ cells were FACS-sorted directly to Trizol® LS (Life Technologies) according to the manufacturer's protocol (n=2-3; from at least 3 animals each). RNA extraction and RT-PCR were conducted as previously described[13]. The sequences of the primers designed with the Primer-BLAST tool are as follows: *Spp1 F: CTGGCTGAATTCTGAGGGACT, R: TTCTGTGGCGCAAGGAGATT; P2ry12 F: GTGCAAGGGGTGGCATCTA, R: TGGAACTTGCAGACTGGCAT; Tmem119 F: TGCAATGTCGCTGTCACTCT, R: AGTTTGTGTTTCCACGGGGT Grp34 F: GGAAAGCTTCAACTCAGTTCCTG, R: TCCATGAGAGGAGCAAAGCC, Rpl27 F: ACATTGACGATGGCACCTC, R: GCTTGGCGATCTTCTTCTTG*

### SUPPLEMENTARY FIGURES

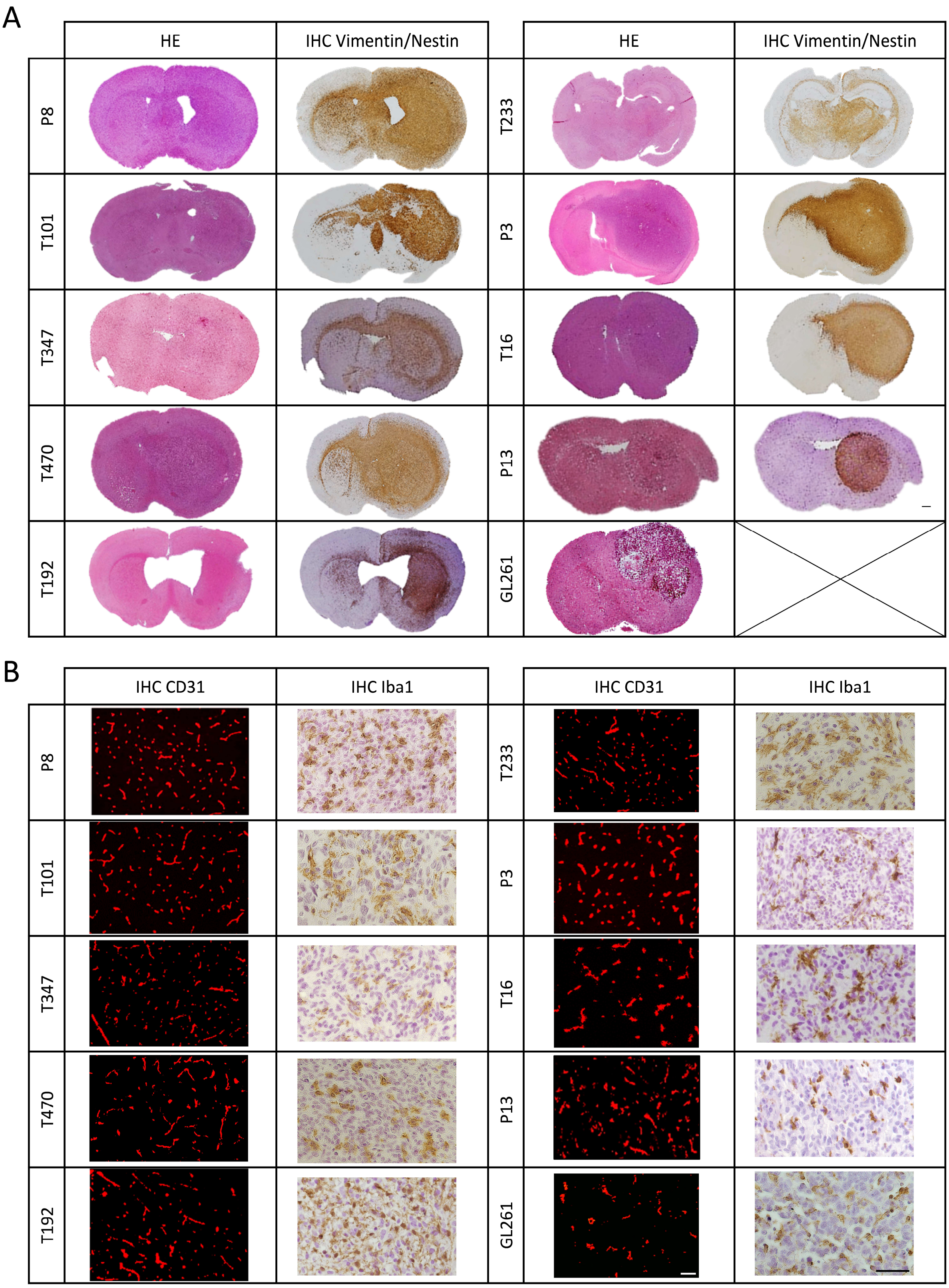


**Figure S1. Characteristics of GBM PDOX models and the GL261 mouse model. (A)** Hematoxylin and Eosin (H&E) and human-specific Vimentin/Nestin staining of representative PDOXs and GL261 at the endpoint of tumor growth. Human-specific Vimentin/Nestin staining highlights varying levels of invasion in PDOX models. Images represent coronal sections of the entire mouse brains. Mice were sacrificed at the endpoint of tumor growth. Scale bars represent 1mm. **(B)** Mouse-specific CD31 and Iba1 stainings show aberrant blood vessels and myeloid cells respectively. Brains of representative PDOXs and GL261 at the endpoint of tumor growth were used for the stainings. Images represent magnified areas in the tumor core. Hematoxylin staining was applied to assess tumor cell density. Scale bars represent 100µm (white) and 50 µm (black).

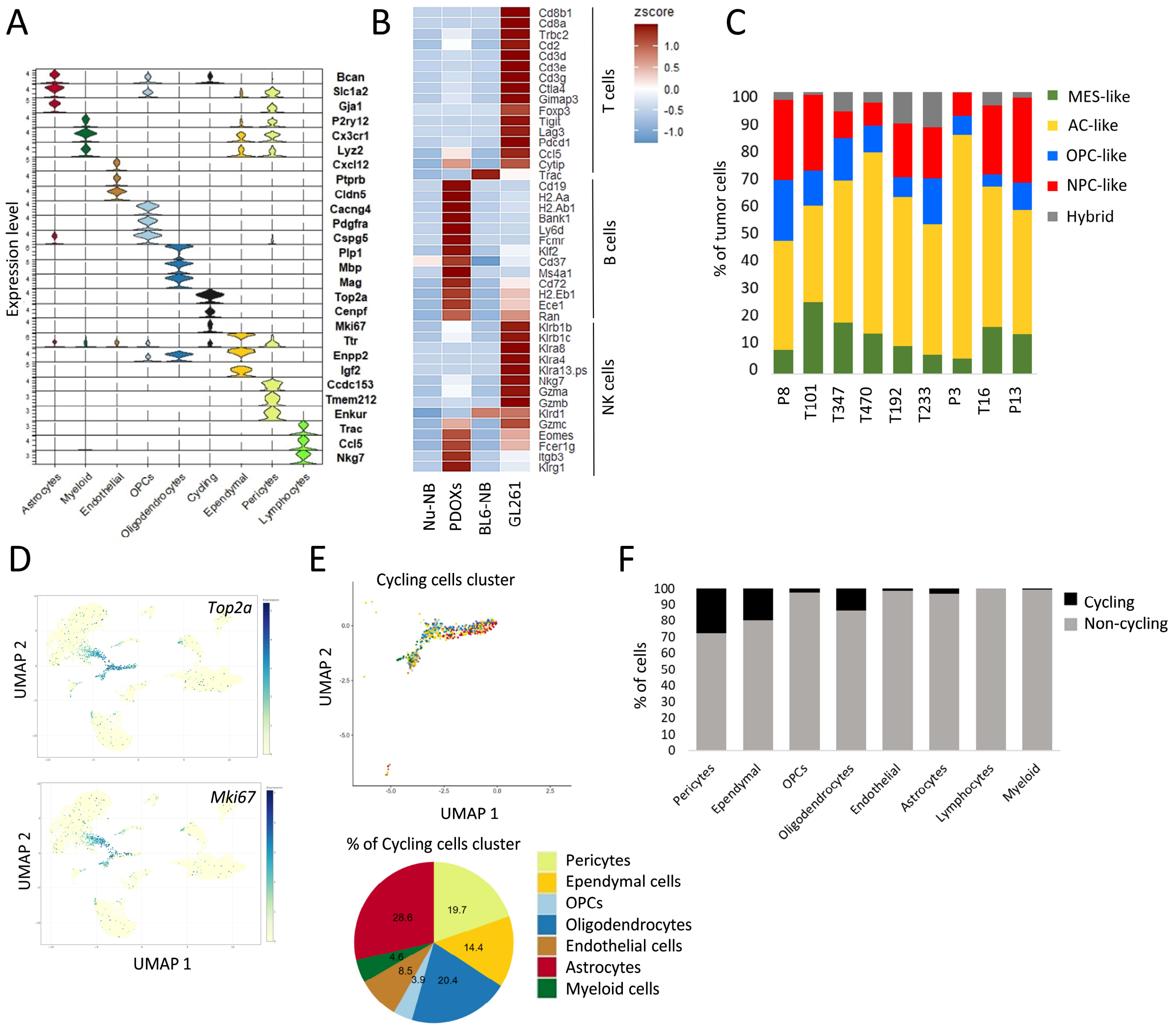


**Figure S2. Composition of mouse-derived TME in PDOXs and GL261. (A)** Violin plots showing expression of marker genes specific to distinct cell types identified in the scRNA-seq dataset Three top markers of each population are displayed (Wilcoxon Rank Sum Test). For entire gene list see **Table S3**. **(B)** Analysis of the lymphocyte cell cluster in nude mouse normal brain (Nu-NB), PDOXs (9 models), C57BL6/N mouse normal brain (BL6-NB), and GL261 tumor (3 collection time points: early, middle, and late). Heatmap shows the z-score of the average expression of marker genes for T, B and NK cells in lymphocytes detected in each sample group. **(C)** Human tumor cells of PDOX models assigned to four GBM transcriptomic states[12]: mesenchymal-like (MES-like), astrocytic-like (AC-like), oligodendrocyte progenitor cell-like (OPC-like) and neural progenitor cell-like (NPC-like). Cells with shared states were grouped as “Hybrid”. **(D)** UMAPs showing expression of cell cycle marker genes *Top2a*, *Mki67* in TME. The color gradient represents expression levels. **(E)** Top: UMAP of the ‘cycling cells’ cluster and the proportion of cell types identified in the cluster. Cell types are color-coded. Bottom: proportions of identified cell types within the ‘cycling cells’ cluster. **(F)** Proportions of cycling cells in each cell type. 100% represents the sum of cell numbers identified in each original cell type cluster and numbers of the same cell types identified in the ‘cycling cells’ cluster.

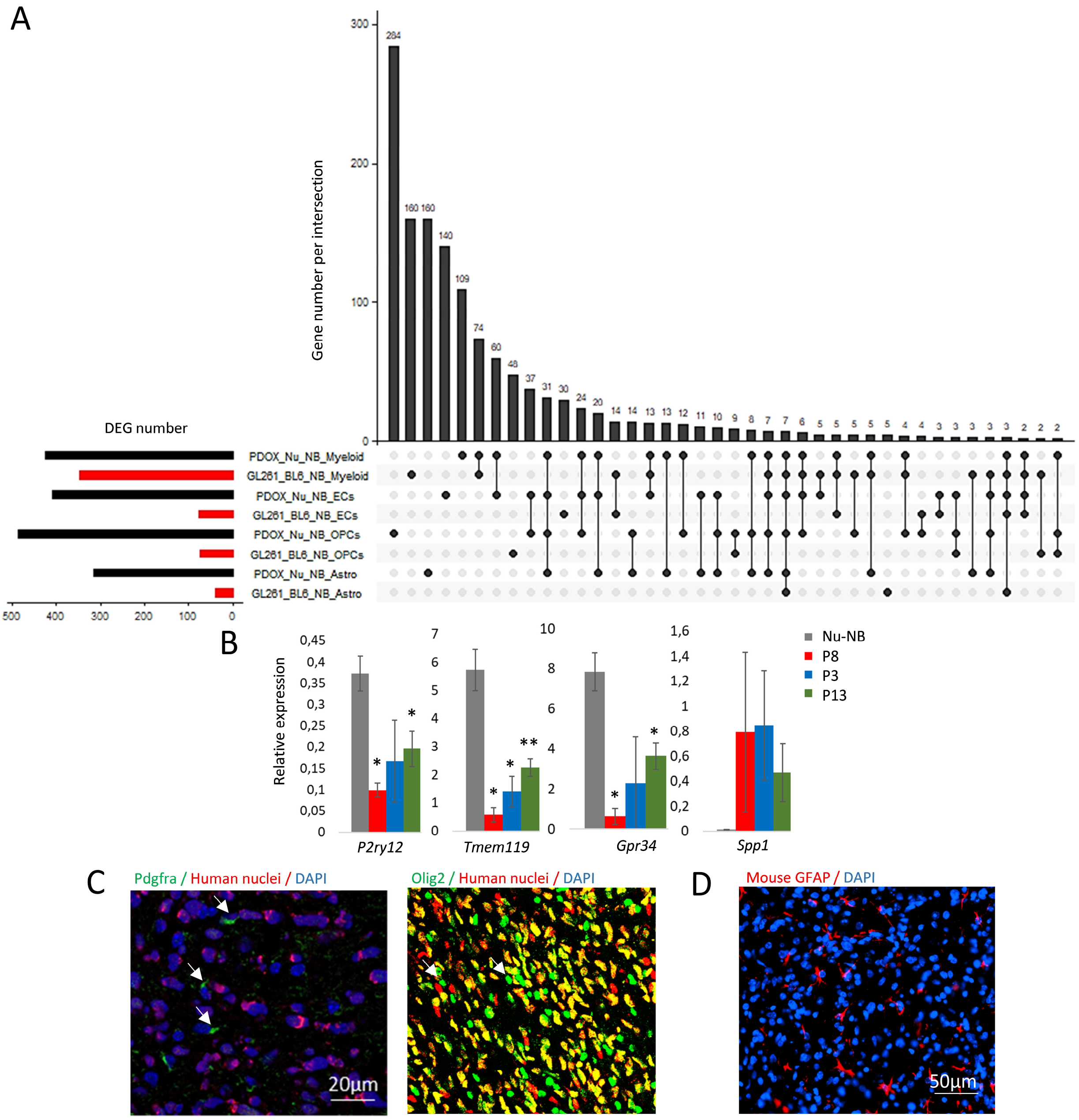


**Figure S3. Tumor-specific activation of TME cell types. (A)** Upset plot showing differentially expressed genes (DEGs) between TME of PDOX and GL261 versus corresponding normal brains in identified cell types (FDR ≤0.01, $|\log_2 FC| \geq 1$, Wilcoxon rank sum test with Benjamini-Hochberg correction). Bar chart depicts the number of differentially expressed genes shared between the different groups. **(B)** Decreased expression of homeostatic Mg markers (*P2ry12*, *Tmem119, Gpr34*) and increased levels of a TAM marker (*Spp1*) were validated by qPCR in $CD11b^+$ myeloid cells isolated as bulk from TME of representative PDOX models (P3, P8, P13) and nude normal brain (Nu-NB); graphs show bulk mean relative expression levels ± SEM (n=2-5, **p<0.01 *p<0.05, two-tailed Student's t-test). *Rpl27* was used as a housekeeping gene. **(C)** Immunofluorescence staining showing Pdgfrα$^+$ and Olig2$^+$ cells in the tumor core of PDOX P8. Arrows point Pdgfrα$^+$ and Olig2$^+$ mouse cells, negative for human nuclei staining. **(D)** Immunofluorescence staining showing mouse GFAP$^+$ astrocytic cells in the tumor core of PDOX T192.

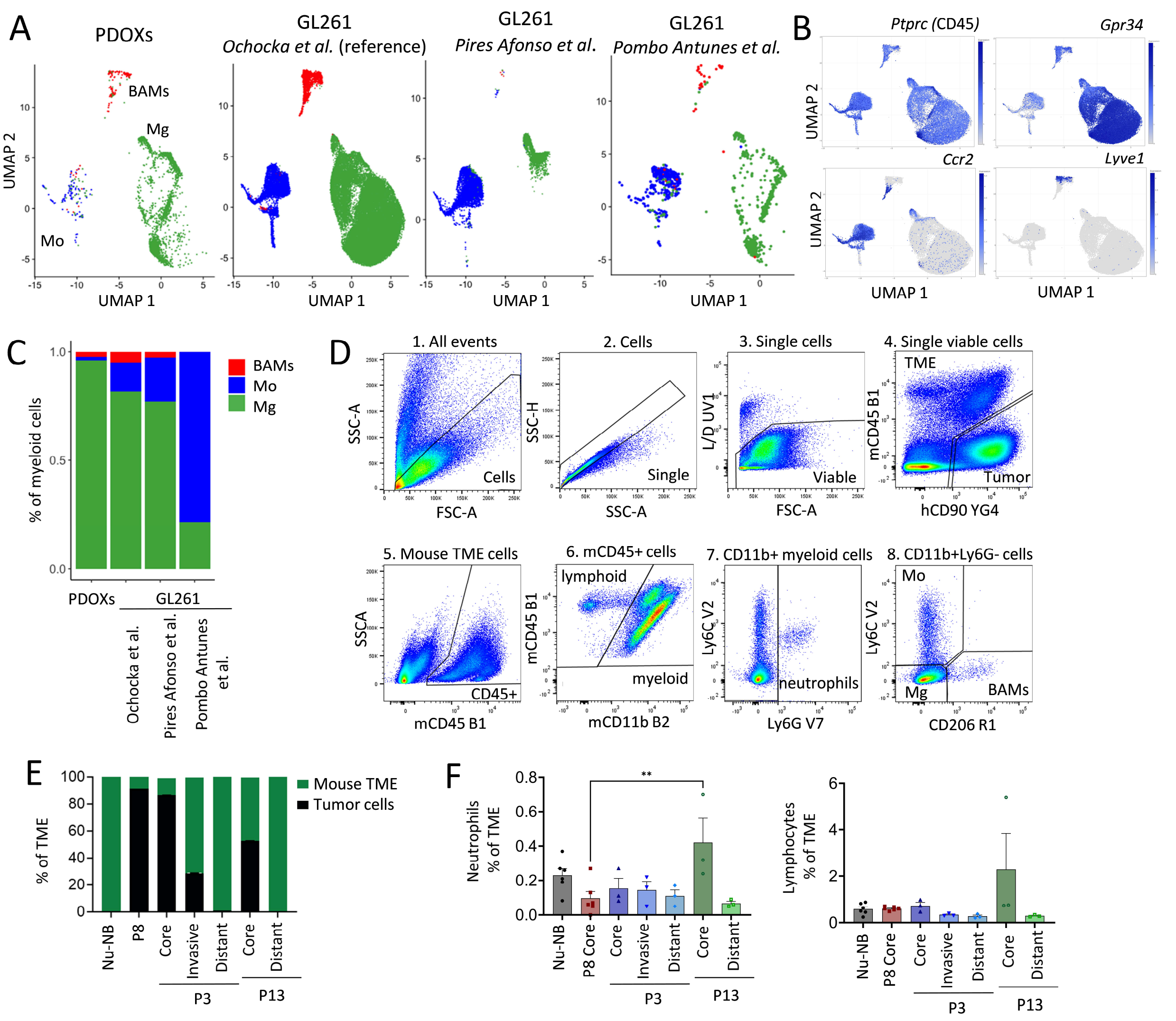


**Figure S4. Myeloid cell heterogeneity and ontogeny in mouse-derived GBM TME. (A)** Split UMAP projection of myeloid cells from different datasets. Myeloid cell types are color-coded (Mg = green, Mo = blue, BAMs = red). **(B)** UMAPs showing expression of marker genes: pan-immune *Ptprc* (CD45), Mg *Gpr34*, monocytes *Ccr2*, BAMs *Lyve*. The color gradient represents expression levels. **(C)** Proportions of myeloid cell types in PDOXs and GL261 for three independent datasets[9,14,15]. Percentage of Mo depends on the tumor stage and dissection strategy. **(D)** Gating strategy for flow cytometry. Example is shown for PDOX P13 tumor: (1) Cells were distinguished from debris based on the Forward Scatter (FSC) and Side Scatter (SSC). (2) Cell aggregates were gated out based on their properties displayed on the SSC area (SSC-A) versus height (SSC-H) dot plot. (3) Dead cells were recognized by their strong positivity for the dead cell marker (L/D) (4) Human tumor cells were recognized as CD90 positive with human-specific antibody. TME was of mouse origin. (5) Immune cells were recognized as CD45 positive within mouse-derived TME. (6) Lymphocytes were recognized as CD11b negative population. Negative gating includes also $CD11b^{low}$ NK cells. CD11b positive cells correspond to myeloid cells. (7) $CD11b^{+}Ly6C^{low}Ly6G^{+}$ cells represent neutrophils. (8) Ly6C and CD206 allows distinguishing $Ly6C^{+}CD206$- Mo, $Ly6C^{-}CD206^{-}$ Mg and $Ly6C^{-}CD206^{+}$ BAMs. **(E)** Ratio of human tumor cells over mouse-derived TME across distinct brain areas in PDOXs P8, P3 and P13 (≥3 mice/condition, mean ± SEM). The ratio of tumor cells to TME was higher in the tumor core compared to invasive zone and distal brain regions. **(F)** Flow cytometry quantification of $CD45^{+}CD11b^{+}Ly6G^{+}Ly6C^{low}CD206^{-}$ neutrophils and $CD45^{+}CD11b^{-}$ lymphocytes in Nu-NB and PDOX TME across different brain regions in PDOXs P3, P8 and P13 ($n \geq 3$ mice/condition, mean ± SEM, one-way ANOVA, $^{**}p<0.01$).

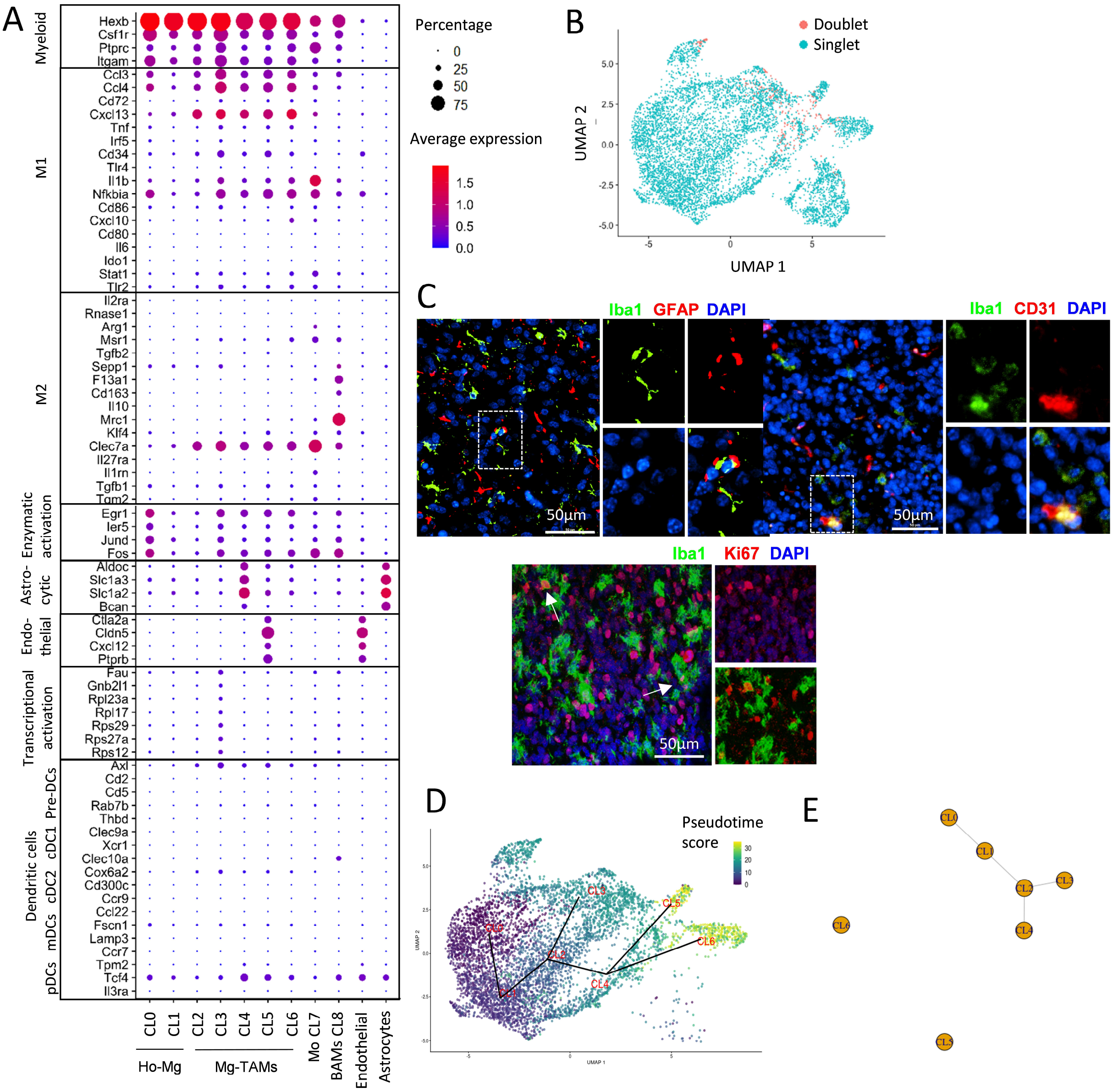


**Figure S5. Analysis of heterogeneous myeloid phenotypic states. (A)** Expression levels of exemplary genes representing myeloid pan-markers, M1/M2 markers, enzymatic activation genes, astrocytic and endothelial Mg-TAM markers, transcriptional activation markers and key markers identifying distinct subsets of dendritic cells (DCs): pre-DCs, cDC1, cDC2, migratory DCs (mDCs) and pDCs per identified myeloid cell cluster: CL0-1 Ho-Mg, CL2-6 Mg-TAMs, CL7 Mo, CL8 BAMs. Astrocytes and endothelial cells were presented to compare marker genes identified in CL4 astrocytic-like and CL5 endothelial-like Mg-TAMs. **(B)** Scoring for putative doublets in the myeloid cells. **(C)** Representative immunofluorescence images showing mouse Iba1$^{+}$ myeloid cells positive for GFAP (astrocytic marker, PDOX T192), CD31 (endothelial marker, PDOX T470) and Ki67 (cell cycle marker, PDOX P8). **(D)** TSCAN-based trajectory inference analysis of Mg clusters (CL0-6) showing a trajectory from Ho-Mg to Mg-TAMs. The Pseudotime ordering of cells is presented by a color code on the original UMAP projection. (**E**) Minimum spanning tree (MST) of the Mg clusters after introducing an outgroup to split unrelated trajectories.

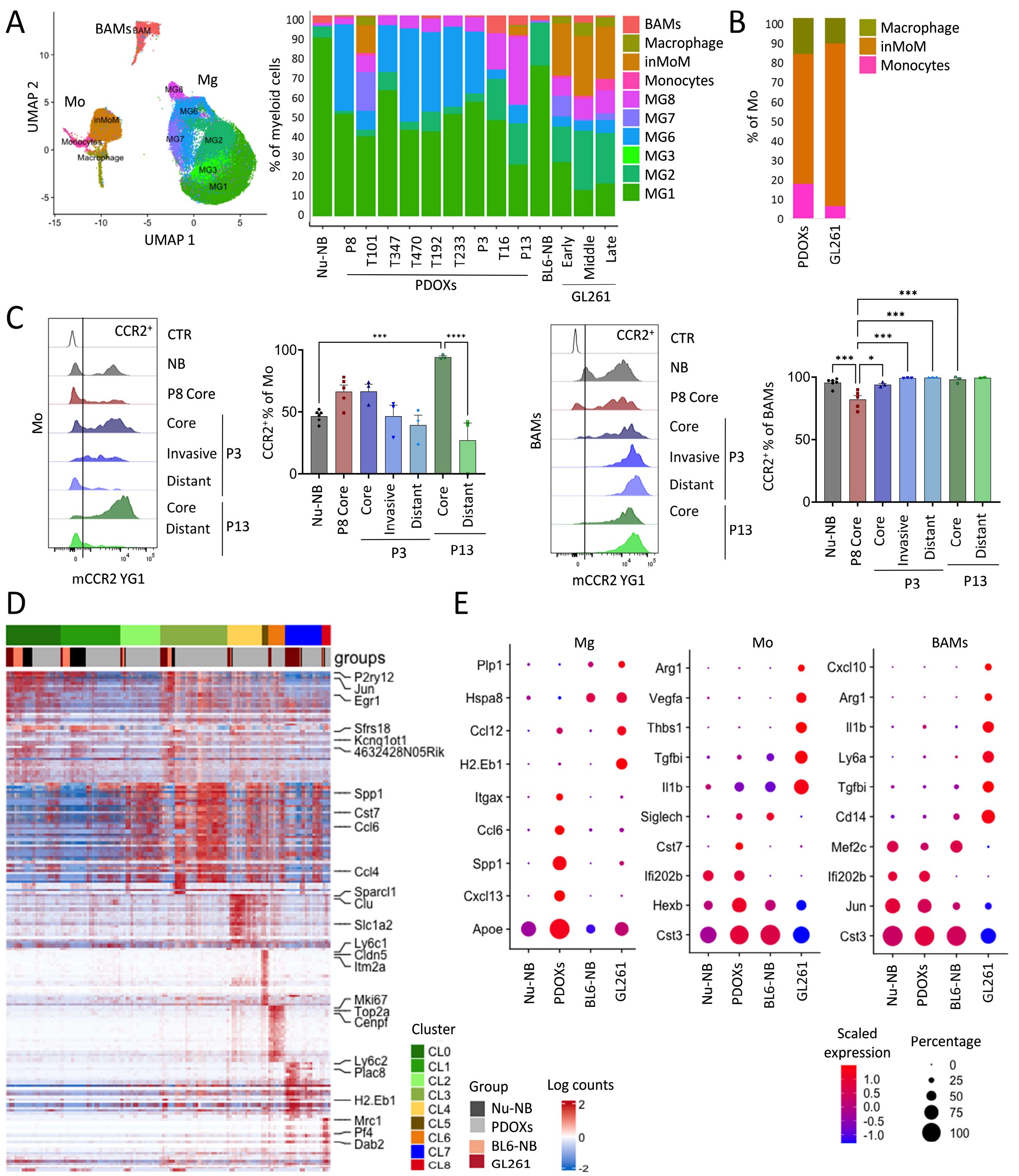


**Figure S6. GBM-specific education of myeloid cells in PDOXs and GL261 TME. (A)** Reference-based analysis of myeloid cell states described by Ochocka et al. in PDOXs and GL261: (i) MG1-MG8 states correspond to Mg, (ii) Monocyte, Intermediate Monocytes/Macrophage (inMoM) and Macrophage states correspond to Mo, no further phenotypic states were identified for BAMs. **(B)** Proportions of phenotypic states identified by reference-based analysis in Mo in PDOXs (4 models with Mo cells detected combined) and GL261 (3 time points combined). **(C)** Representative flow cytometry graphs and quantification of CCR2$^+$ cells in CD45$^+$CD11b$^+$Ly6G$^-$Ly6C$^+$CD206$^-$ Mo and CD45$^+$CD11b$^+$Ly6G$^-$Ly6C$^-$CD206$^+$ BAMs in PDOXs P8, P3, P13. For PDOXs P3 and P13 invasive zone and distant normal brain areas were also collected (n≥3 mice/condition, mean ± SEM, one-way ANOVA, ****p<0.0001, ***p<0.001, *p<0.05). **(D)** Comparison of transcriptomic profiles of myeloid cells in PDOXs and GL261 models. Heatmap represents expression of marker genes of each myeloid transcriptomic state identified (CL0-8). No major differences were observed between myeloid cells of PDOXs and GL261. **(E)** Expression levels of representative genes in Mg, Mo and BAMs for nude mouse normal brain (Nu-NB), PDOXs (9 models), C57BL6/N mouse normal brain (BL6-NB), GL261 tumor (3 collection time points: early, middle, and late).

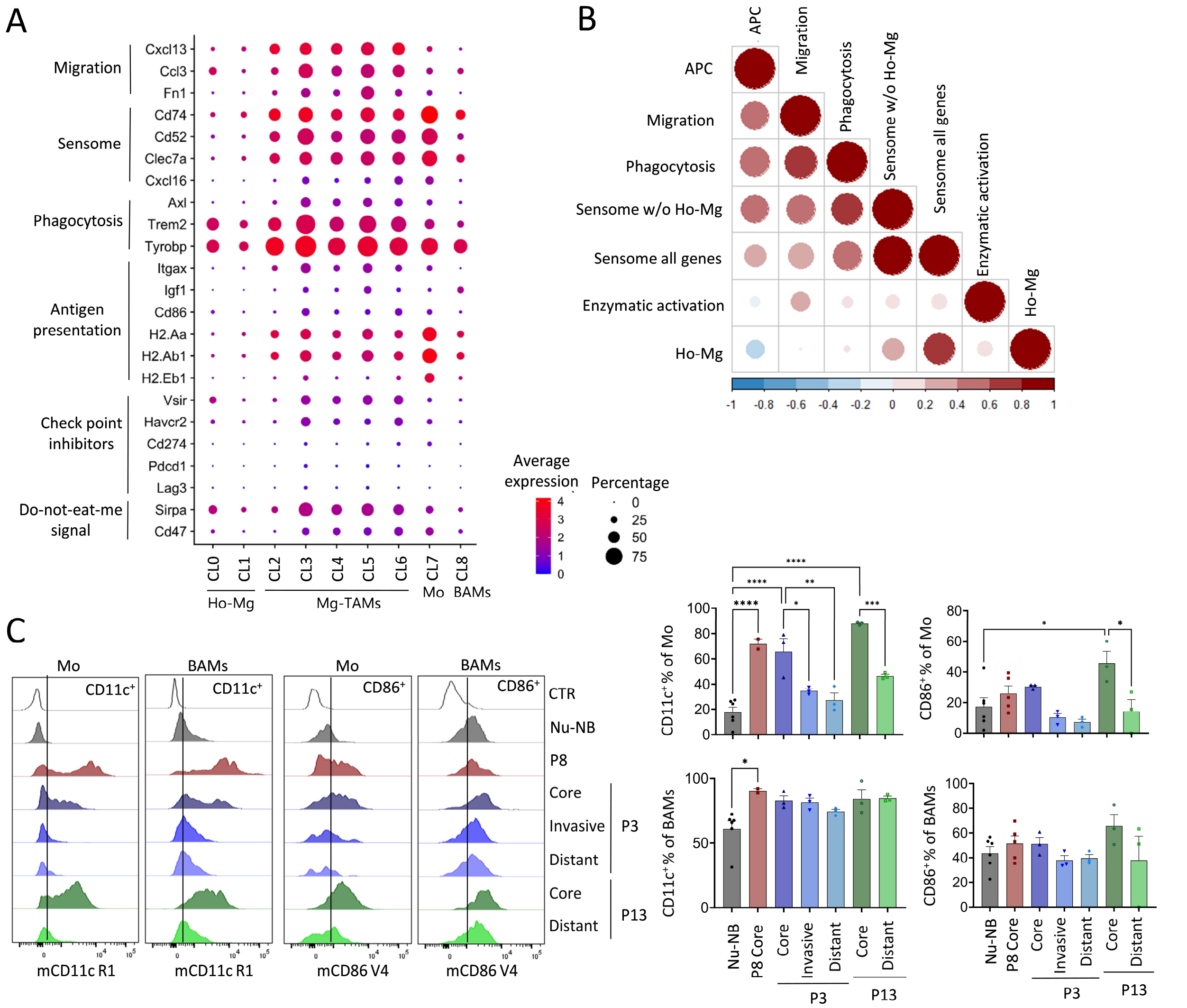


**Figure S7. Analysis of functional markers in TAMs. (A)** Expression levels of representative genes associated with functional signatures for migration, sensome (GBM specific), phagocytosis, antigen presentation, checkpoint inhibition and “do-not-eat-me signal” per identified myeloid cell cluster: CL0-1 Ho-Mg, CL2-6 Mg-TAMs, CL7 Mo, CL8 BAMs. **(B)** Spearman correlation analysis between functional signatures in myeloid cells. **(C)** Representative flow cytometry graphs and quantification of CD11c$^+$ and CD86$^+$ cells in CD45$^+$CD11b$^+$Ly6G$^-$Ly6C$^+$CD206$^-$ Mo and CD45$^+$CD11b$^+$Ly6G$^-$Ly6C$^-$CD206$^+$ BAMs in nude mouse normal brain (Nu-NB) and PDOXs P8, P3, P13. For PDOXs P3 and P13 invasive zone and distant normal brain areas were also collected. Unstained control is shown for each population (CTR) (n≥3 mice/condition, mean ± SEM, one-way ANOVA, **0*p<0.0001, **p<0.01, *p<0.05).

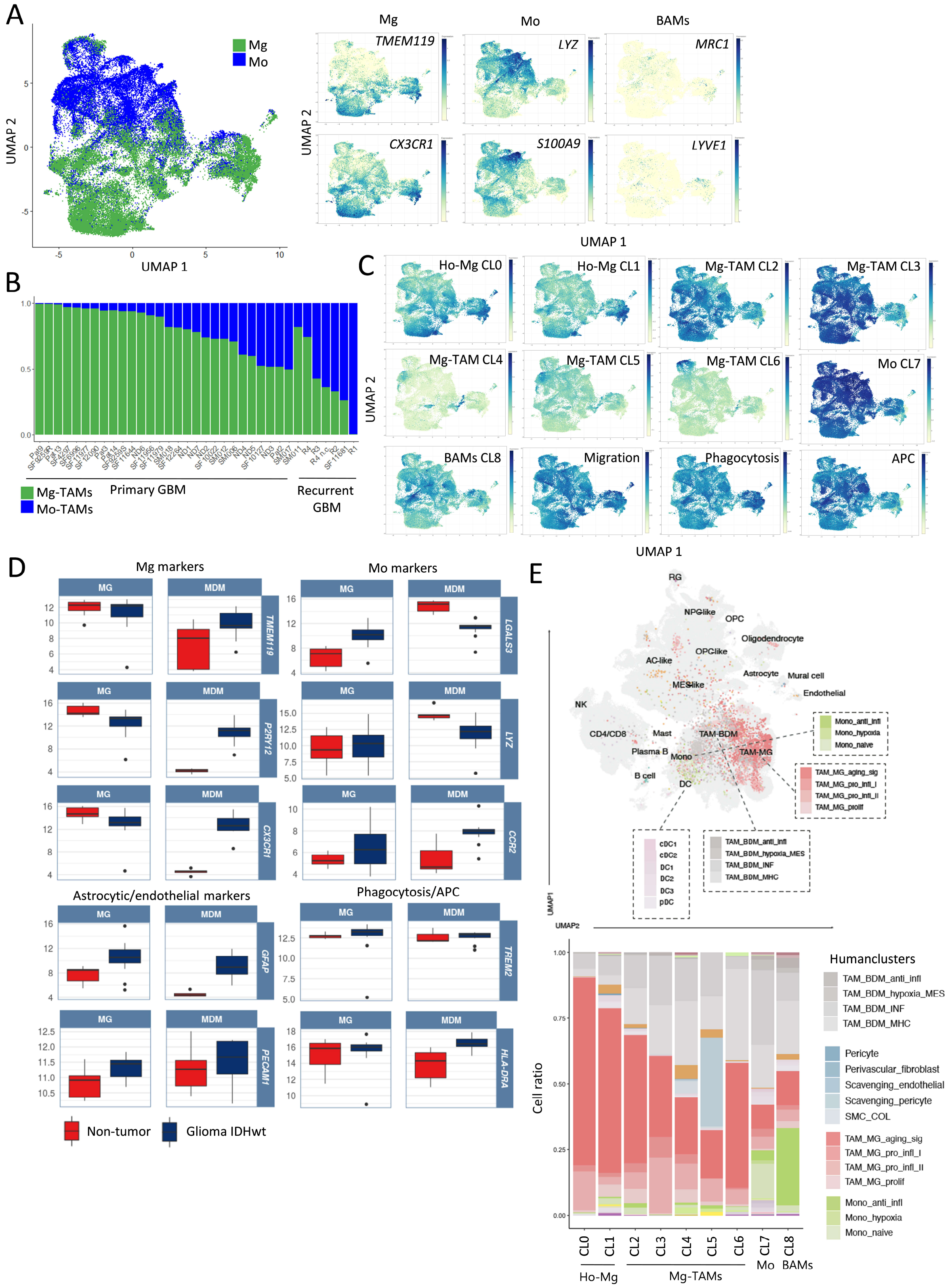


**Figure S8. Analysis of Mg and Mo cells in GBM patient tumors. (A)** UMAP projection of myeloid cells in GBM patient tumors (n=36 patient tumors combined). Cells are color coded for Mg and Mo ontogeny based on established gene signatures. Inserts show expression levels of marker genes for Mg, Mo and BAMs. The color gradient represents expression levels. **(B)** Proportions of Mg and Mo in individual GBM patient tumors (36 patient tumors: 27 newly diagnosed and 9 recurrent tumors). Recurrent GBMs show higher proportions of Mo ($p<0.01$, two-tailed Student's t-test). **(C)** UMAP projection of human GBM myeloid cells displaying expression levels of cluster (CL0-8) and functional signatures identified in PDOX TME. The color gradient represents expression levels. (**D**) Expression of representative Mg, Mo, astrocyte and endothelial cell markers showing convergence of Mg and Mo towards heterogeneous TAMs in GBM tumors. Bulk RNA-seq analysis was performed in purified $CD49d^+$ (MG) and $CD49^-$ (MDM) myeloid cells[16] representing Mg and Mo in normal human brain (n=7) and GBM patient tumors (n= 15). While Mg decreases homeostatic genes, Mo acquire Mg profiles within GBM tumors. Reversed convergence is observed for Mo markers. (**E**) Mapping of mouse myeloid cells to the scRNA-seq profiles of human GBMs (110 patient tumors, GBmap[17]). Mg states (CL0-CL6) predominantly mapped to previously identified human Mg states, although Mg-TAMs (CL3, CL4, CL6) showed increased proportion of TAMs associated with blood-derived macrophages (BDMs), highlighting strong convergence of the Mg and Mo towards comparable TAM signatures. The astrocytic-like CL4 state is clearly associated with human myeloid compartment, whereas the endothelial-like CL5 state only partially overlapped with pericytes and endothelial cells. Mo (CL7) and BAMs (CL8) showed a stronger association with Mo signatures.

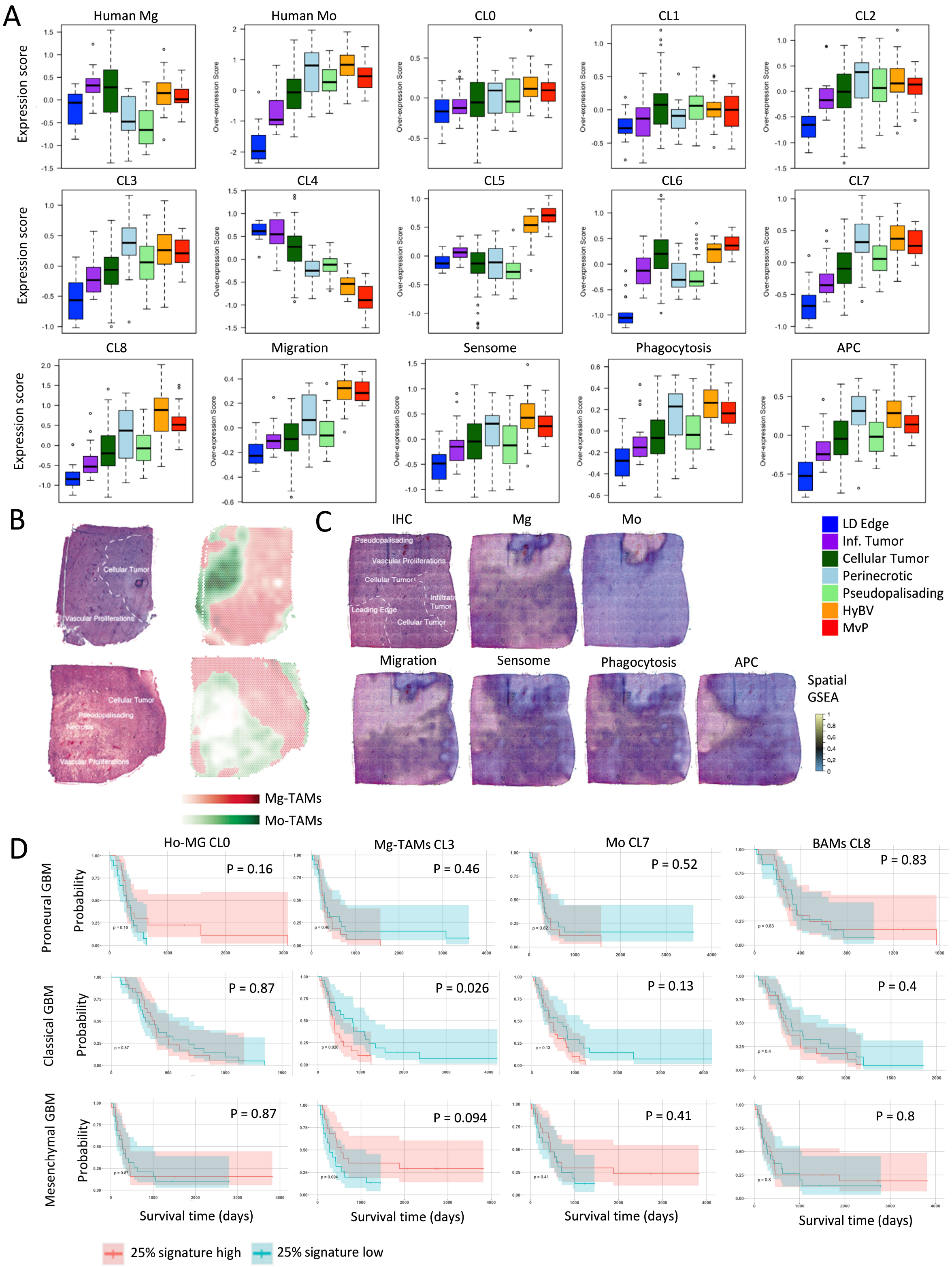


**Figure S9. Assessment of myeloid signatures in bulk RNA-seq profiles of GBM patient tumors. (A)** Bulk RNA-seq profiles of laser-micro dissected regions of GBM patient tumors. Box plots show signatures of Mg/Mo, phenotypic states and functional signatures per tumor niche (LD edge: leading edge; Inf tumor: infiltrative tumor; HyBV: hyperplastic blood vessels in cellular tumor; MvP: microvascular proliferation). Data extracted from the Ivy Glioblastoma Atlas Project obtained from 279 samples from 44 tumors. The box limits indicate the 25th and 75th percentiles, centerlines show the medians; whiskers show a minimum of two measures: extreme values or 1.5 IQR from the closest quartile; points depict outliers. **(B-C)** Representative surface plots of spatial localization of myeloid cell signatures in GBM patient tumors. **(D)** Kaplan–Meier survival analysis in GBM patients (CGGA-GBM IDHwt divided into transcriptomic subtypes: Classical, Mesenchymal and Proneural) with high- (red) and low (blue) -enriched signatures (25 percentile top/bottom).

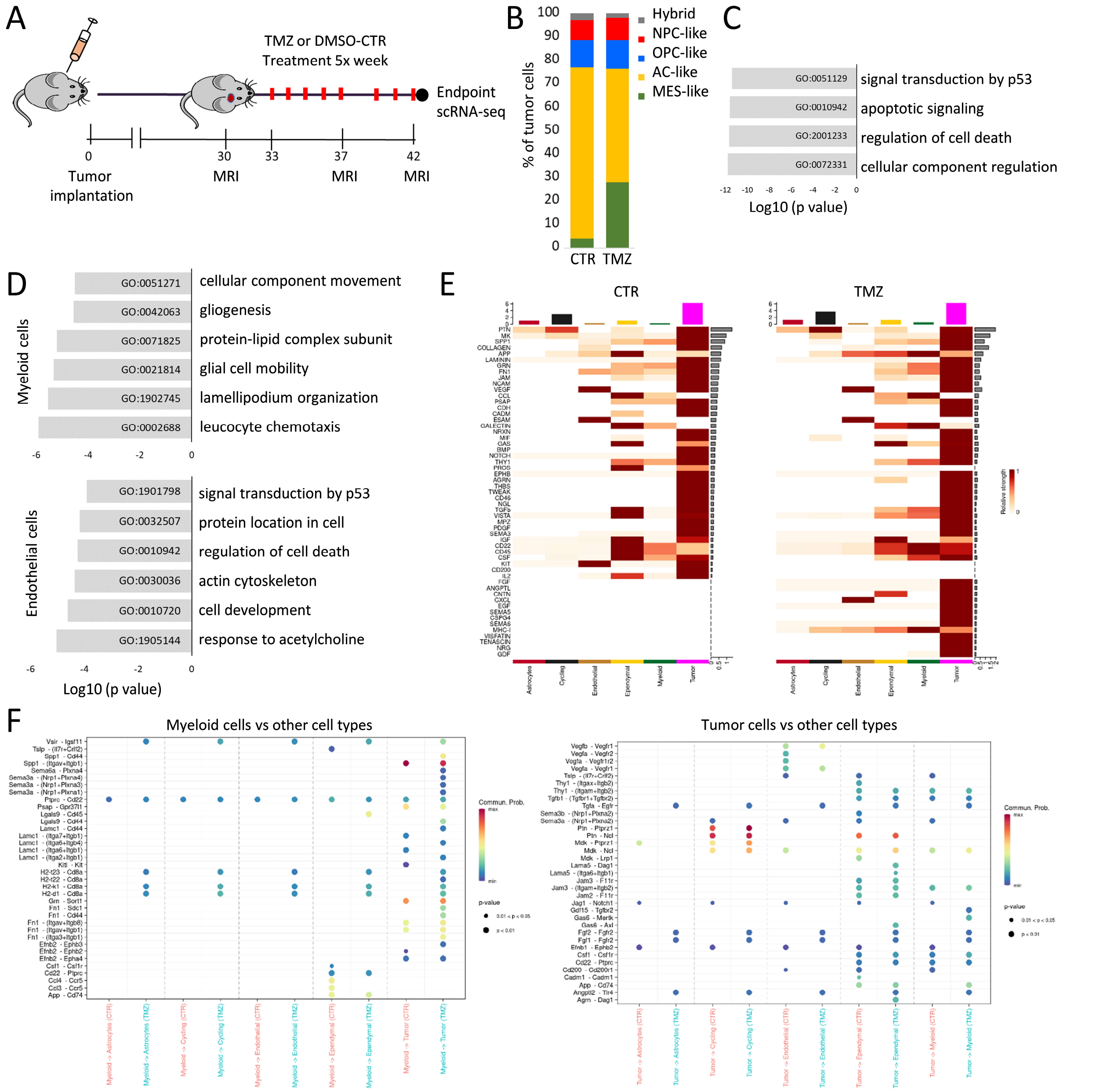


**Figure S10. GBM ecosystem upon TMZ treatment. (A)** Schematic illustration of the TMZ treatment regimen in PDOX P3 *in vivo*. Day 0 = tumor implantation, day 30 = MRI validation of tumor growth, day 33= treatment start, day 37= intermediary MRI, day 42 = endpoint. **(B)** Assessment of GBM cellular states[12] at single cell level in TMZ-treated and CTR GBM cells (n=1 PDOX brain per condition). **(C)** Summary of top four gene ontology terms characterizing DEGs in TMZ-treated versus CTR GBM cells (FDR ≤0.01, |logFC| ≥ 0.5 the Wilcoxon rank sum test with Benjamini-Hochberg correction). **(D)** Summary of top six gene ontology terms characterizing DEGs in TMZ-treated versus CTR myeloid and endothelial cells forming TME in PDOX P3 (FDR ≤0.01, |logFC| ≥ 0.5). **(E)** CellChat-based overall signaling patterns per identified cell type, visualized as heatmaps, in CTR and TMZ-treated PDOXs. **(F)** Predicted ligand-receptor interactions based on communication probability are shown for signals outgoing from myeloid and tumor cells.
