## Supplementary Tables for "Glioblastoma-instructed microglia transition to heterogeneous phenotypic states with phagocytic and dendritic cell-like features in patient tumors and patient-derived orthotopic xenografts"

| Table S1. | Characteristic of PDOX models applied in the study. | | | | | | | | | | | | | | | | |
|---|---|---|---|---|---|---|---|---|---|---|---|---|---|---|---|---|---|
| Model | Patient information | | | | | | | PDOX model characteristics | | | | | | Molecular features in PDOXs | | | |
| PDOX model | Patient code | Age at collection | Sex | Tumor localization | Surgery | Treatment received prior PDOX | Histological diagnosis | Implantation | PDOX generation reached | PDOX survival time (days) | Proliferation index (%Ki67 +/- SD) | Detection by MRI | Ploidy tumor cells | Copy number variations (CNV) | Glioma-specific gene mutations | Brain tumor classifier | MGMT promoter methylation |
| P3 | BER0003 | 64 | Male | NA | NA | NA | GBM Grade IV | organoids | >G10 | 42 +/- 5.2 | 54.7 +/- 3.7 | detectable, quantifiable | 2.1N | +[Chr7, 19p, 20q], -[1p36-1p34.1, 1q21.1-q44, -5p15.33-31, Chr9, Chr10, 11p15-14, 20p] --CDKN2A/B | PTEN (chr10:g.87957923del, ENSP00000361021.3:p.Asp236ThrfsTer20)<br>TP53 (chr17:g.7675085C>T, ENSP00000269305.4:p.Cys176Tyr) | GBM, IDHwt, RTK II | methylated |
| P8 | BER0008 | 64 | Female | NA | NA | NA | GBM Grade IV | organoids | >G10 | 52.5 +/- 3 | 38.3 +/- 10.8 | detectable, quantifiable | 2.2N | ++EGFR, +[5p15.3-p12, 5q31.1-q35.3, Chr7, 8q24], -[ 6q21-q27, Chr10, 13q13.3-q34, 15q21.2-q23, 18q21.2-q22.4], --CDKN2A/B | EGFR (chr7:g.55154128G>A, ENSP00000275493.2:p.Ala289Thr)<br>MET (chr7:g.116771573G>T, ENSP00000317272.6:p.Ala954Ser)<br>PTEN (chr10:g.87961047_87961050del, ENSP00000361021.3:p.Thr319Ter);<br>ATRX (chrX:g.77663459C>G, ENSP00000362441.4:p.Ser1348Thr)<br>ATRX (chrX:g.77683638T>A, ENSP00000362441.4:p.Ser540Cys) | GBM, IDHwt, RTK I/II | methylated |
| P13 | BER0013 | NA | Female | NA | NA | NA | GBM Grade IV | organoids | >G10 | 35 +/- 2.5 | 42.9 +/- 10.8 | detectable, non quantifiable | 2.3N | +(Chr7, Chr19, Chr20), -(1p21.1-p31.2, 6q16.3-q21, Chr10, Chr13, 17q11-12), --CDKN2A/B | NF1 (chr17:g.31221849G>A, - )- likely not pathogenic | GBM, IDHwt, RTK II | methylated |
| T16 | LIH0016 | 52 | Female | Fronto-parietal right | recurrence | TMZ | GBM Grade IV | organoids | >G10 | 58.5 +/- 5 (>G7) | 18.9 +/- 5.8 | detectable, quantifiable | (2.1N) + 3.4N | ++[EGFR, MDM4], +7q, - [6q, Chr10, 13q11-q31.1], --CDKN2A/B | | GBM, IDHwt, RTK II | unmethylated |
| T101 | LIH0101 | 60 | Male | NA | NA | NA | GBM Grade IV | organoids | G7 | 106 +/- 5.5 | 13.4 +/- 4.8 | detectable, non quantifiable | (2N) + 4.6N | ++[EGFR, MDM2] +[Chr1, Chr7, Chr9, Chr13, Chr16, Chr17, Chr19] -[3q, Chr4, Chr6, Chr10, Chr11, 14q11, Chr15], --CDKN2A/B, | No Glioma-specific mutations | GBM, IDHwt, RTK II/I | methylated |
| T158 | LIH0158 | 70 | Female | Frontal right | primary | no prior treatment | GBM Grade IV | organoids | G5 | 68 +/-1.6 | 35.2+/-3 | detectable, quantifiable | 2.3N | [++MDM2], +[Chr 7, Chr19, Chr20], -[Chr10, Chr15q], --CDKN2A/B | CDK6 (chr7:g.92615172T>A, ENSP00000265734.4:p.Ser317Cys)<br>NF1(chr17:g.31163302G>A, ENSP00000491431.1:p.Gly31Glu)- likely not pathogenic | GBM, IDHwt, RTK II | unmethylated |
| T185 | LIH0185 | 76 | Female | Occipital right | NA | no prior treatment | GBM Grade IV | organoids | G6 | 89 +/- 10 | 19.4 +/- 6.9 | detectable, non quantifiable | 2.2N | ++EGFR, +[Chr7, Chr20], -[Chr 10q, Chr22q] --CDKN2A/B | EGFR (chr7:g.55165350G>T, ENSP00000275493.2:p.Gly598Val)<br>MET (chr7:g.116695757A>T, ENSP00000413857.1:p.Met1?)<br>NF1 (chr17:g.31350201A>G, ENSP00000348498.3:p.Glu2426Gly)<br>STAG2 (chrX:g.124090748A>G, ENSP00000218089.9:p.Thr1151Ala) | GBM, IDHwt, RTK I/II | methylated |
| T186 | LIH0186 | 34 | Male | NA | NA | no prior treatment | Anaplastic Oligodendroglioma Grade III | fresh organoids/tissue | G9 | 159 +/- 43 | 33+/-4.8 | detectable, non quantifiable | 2.1-2.4N | complex genome ++[MYCN, CDK4] | IDH1 (chr2:g.208248388C>T, ENSP00000260985.2:p.Arg132His)<br>PDGFRA (chr4:g.54285926A>T, ENSP00000257290.5:p.Asp842Val)<br>MET (chr7:g.116771936C>T, ENSP00000317272.6:p.Thr1010Ile)<br>RET (chr10:g.43100551C>A, ENSP00000344798.4:p.Leu56Met)<br>TP53 (chr17:g.7673803G>A, ENSP00000269305.4:p.Arg273Cys)<br>ATRX (chrX:g.77652140C>A, ENSP00000362441.4:p.Glu1511Ter) | Glioma, IDHm, high grade astrocytoma | methylated |
| T188 | LIH0188 | 68 | Male | Temporal right | primary | no prior treatment | GBM Grade IV | organoids | G6 | 63+/-3,94 | 37.1+/- 2.9 | detectable, quantifiable | 2.3N | [++EGFR], +[Chr7, Chr 19, Chr 20], -[1p36.23, 6p21.32, 6q27, Chr 10, 11q24.2], --CDKN2A/B | MDM4 (chr1:g.204549329A>C, ENSP00000356150.3:p.Lys374Gln)<br>PTCH1 (chr9:g.95459693C>T, ENSP00000332353.6:p.Val932Ile) | GBM, IDHwt, RTK II/I | unmethylated |
| T192 | LIH0192 | 42 | Female | Frontal left | primary | no prior treatment | GBM Grade IV | organoids | G6 | 71+/-17,8 | 26.5 +/-8 | detectable, non quantifiable | 2.4N | ++[EGFR, 2q34], +[1q21.2-24.2, Chr7, Chr19, Chr20], - [9p24.3 - 13.3, Chr10], | EGFR (chr7:g.55154017C>T, ENSP00000275493.2:p.Arg252Cys) | GBM, IDHwt, RTK II | unmethylated |
| T226 | LIH0226 | 61 | Male | Supratentoriell | NA | NA | GBM Grade IV | organoids | G5 | 127+/-4,9 | NA | detectable, non quantifiable | NA | ++[EGFR], +[Chr 5p, Chr7, Chr 20p], -[1p21.1-p31.2, 6p22.3, 6q16.3-q21, Chr 10, 14q, 16p12.2, 19q, 22q], , --CDKN2A/B | PIK3R1 (chr5:g.68296465A>C, ENSP00000513328.1:p.Gln235His)<br>EGFR (chr7:g.55173057G>A, ENSP00000275493.2:p.Gly665Asp)<br>PTEN (chr10:g.87933154G>A, ENSP00000361021.3:p.Gly132Asp) | GBM, IDHwt, RTK I | methylated |
| T233 | LIH0192 | 42 | Female | Frontal left | recurrence 1 | radiotherapy + TMZ | GBM Grade IV | organoids | G5 | 148+/-7,83 | 11,3 | no detectable | 2.3N | ++[EGFR, 2q34], +[1q21.2-24.2, Chr7, Chr19, Chr20], - [9p24.3 - 13.3, Chr10, Chr 14q] | | GBM, IDHwt, RTK II/I | methylated (v. wide range) |
| T238 | LIH0238 | 42 | Male | NA | NA | no prior treatment | GBM Grade IV | organoids | G3 | 100+/-8,77 | 22.1 | detectable, non quantifiable | (2.1N) + 3.7N | complex genome, +[Chr7, Chr19, Chr20] -[4q35.2, 6q, Chr10, 13q, 17q], --CDKN2A/B | MDM4 (chr1:g.204542937C>G, ENSP00000356147.3:p.Thr107Arg)<br>PTCH1 (chr9:g.95447309T>C, ENSP00000332353.6:p.Tyr1316Cys)<br>PTEN (chr10:g.87961095C>T, ENSP00000361021.3:p.Arg335Ter)<br>TP53 (chr17:g.7675157G>A, ENSP00000269305.4:p.Pro152Leu) | GBM, IDHwt, RTK II | methylated |
| T239 | LIH0239 | 80 | Male | Parieto-occipital | NA | no prior treatment | GBM Grade IV | organoids | G3 | 97,5+/-2,34 | 4.5+/-4.9 | NA | 2.3N | +[Chr7, Chr19], -[1p36.32-p34.3, Chr10], --CDKN2A/B | MYCN (chr2:g.15942237C>T, ENSP00000281043.3:p.Thr58Met)<br>PIK3R1 (chr5:g.68280986del, ENSP00000513333.1:p.Asn299MetfsTer19)<br>PTEN (chr10:g.87957955C>T, ENSP00000361021.3:p.Pro246Leu)<br>RB1 (chr13:g.48381415G>A, ENSP00000267163.4:p.Arg556Gln) | GBM, IDHwt, RTK I/II | methylated |
| T251 | LIH0192 | 43 | Female | Frontal left | recurrence 2 | radiotherapy +TMZ | GBM Grade IV | organoids | G7 | 65,2+/-1,5 | 36.1 +/- 7.7 | detectable, non quantifiable | 2.3N | ++[EGFR, 2q34], +[1q21.2-24.2, Chr7, Chr19, Chr20], - [9p24.3 - 13.3, Chr10], | | GBM, IDHwt, RTK II | unmethylated |
| T281 | LIH0281 | 51 | Female | Parietal Left | primary | no prior treatment | GBM Grade IV | organoids | G4 | 126+/-6,06 | 5.3 +/-0.41 | detectable, quantifiable | 2.0N + 3.5N | complex genome, +Chr7, - (partial Chr1, 3p, Chr6, Chr9, Chr10, Chr13, Chr14, Chr16, partial Chr17)--CDKN2A/B | PDGFRA (chr4:g.54274864G>C, ENSP00000257290.5:p.Trp559Cys)<br>PTEN (chr10:g.87933012G>A, -)- likely not pathogenic | GBM, IDHwt, RTK I/II | methylated |
| T304 | LIH0304 | 53 | Male | Frontal left | primary | no prior treatment | GBM Grade IV | organoids | G7 | 89,03+/-4,49 | 6-janv | detectable, quantifiable | 2.2N + 3.2N +4.4N | ++[EGFR, CDK4, MDM2], +[Chr7, Chr20], -(partial Chr1, Chr10, partial Chr12) | PTEN (chr10:g.87933166G>A, ENSP00000361021.3:p.Cys136Tyr) | GBM, IDHwt, RTK I/II | unmethylated |
| T331 | LIH0331 | 84 | Male | Fronto-temporo-parietal left | primary | no prior treatment | GBM Grade IV | organoids | G4 | 116 +/- 8,13 | 33.2 +/- 2.73 | detectable, quantifiable | 2.0N | +[Chr 1, Chr 7, Chr 20], -[9p, Chr 10, Chr 22], --[CDKN2A/B, PTEN] | MDM4 (chr1:g.204549329A>C, ENSP00000356150.3:p.Lys374Gln)<br>EGFR (chr7:g.55143387G>A, ENSP00000275493.2:p.Arg108Lys)<br>EGFR (chr7:g.55181309_55181317dup, ENSP00000275493.2:p.Ala767_Val769dup)<br>RET (chr10:g.43124887C>T, ENSP00000344798.4:p.Arg982Cys) | GBM, IDHwt,RTK I/II | methylated |
| T341 | LIH0337 | 75 | Female | Temporal right | primary | no prior treatment | GBM Grade IV | organoids | G4 | 47,25+/-0,4 | 37.7 +/- 6.1 | NA | 2.1N + 3.65N | ++[PDGFRA, CDK6], - [3q25.3-q26.2, Chr10 ], --CDKN2A/B | TP53 (chr17:g.7673806C>T, ENSP00000269305.4:p.Val272Met) | GBM, IDHwt, RTK I | methylated |
| T347 | LIH0347 | 41 | Male | Frontal left | primary | no prior treatment | GBM Grade IV | organoids | G4 | 125,56+/-5,6 | 42.1+/-1.9 | detectable, non quantifiable | 2.2N | ++EGFR, +[Chr7, 19p, Chr20], - [1p36.3-p34.3, 1q32.3-q44, Chr10, 13q12.11-31.1, 14q11.2-q21.3], --CDKN2A/B | MET (chr7:g.116700208A>G, ENSP00000317272.6:p.Asn375Ser)<br>PTEN (chr10:g.87961003del, ENSP00000361021.3:p.Cys304SerfsTer3) | GBM, IDHwt, RTK I/II | unmethylated |
| T356 | LIH0281 | 52 | Female | Parietal Left, para-sagittal | recurrence 1 | radiotherapy +TMZ | GBM Grade IV | organoids | G5 | 203,6+/-19,74 | 28.17 +/- 4.49 | detectable, quantifiable | 2.1N | complex genome, +Chr7, -(partial Chr1, partial Chr2, partial Chr3, Chr4, Chr6, Chr10, Chr14, Chr17, Chr18) | PIK3CA (chr3:g.179234297A>G, ENSP00000263967.3:p.His1047Arg)<br>PDGFRA (chr4:g.54274864G>C, ENSP00000257290.5:p.Trp559Cys)<br>NF1 (chr17:g.31327521_31327537del, ENSP00000348498.3:p.Val1743AspfsTer12) | GBM, IDHwt, RTK II | unmethylated |
| T361 | LIH0361 | 69 | Female | Temporal left, para-brainstem | NA | no prior treatment | GBM Grade IV | organoids | G5 | 79+/-0 | 46.3+/-1.6 | detectable, quantifiable | 2.3N | ++(EGFR, MDM4) + [Chr7, 19p, 20p13-p11.1, 20q13.1-q13.3], -[3q13.33-q21.1, 6p25.3-p21.32, 10q23.2-q26.3], --CDKN2A/B | EGFR (chr7:g.55143387G>A, ENSP00000275493.2:p.Arg108Lys) | GBM, IDHwt, RTK I/II old II | methylated |
| T363 | LIH0363 | 84 | Female | Occipital lobe right | primary | no prior treatment | GBM Grade IV | organoids | G3 | 140+/-0 | 6.36 | no detectable | 2.1N | complex genome ++EGFR,-[Chr 1, 9p, Chr10, 15q], --CDKN2A/B | MET (chr7:g.116771869C>T, ENSP00000317272.6:p.Arg988Cys)<br>RET (chr10:g.43128249A>G, ENSP00000347942.3:p.Met1109Val) | GBM, IDHwt, RTK I/II | methylated |
| T367 | LIH0367 | 69 | Male | Temporal left | primary | no prior treatment | GBM Grade IV | organoids | G3 | 168+/-0 | 29.7+/-3.1 | detectable, quantifiable | 2.3N | ++[EGFR, MDM4], +[Chr7, Chr19], - Chr10 | PDGFRA (chr4:g.54267424G>A, ENSP00000257290.5:p.Val299Ile)<br>MET (chr7:g.116700208A>G, ENSP00000317272.6:p.Asn375Ser)<br>CDKN2A (chr9:g.21971209C>T, -)<br>PTEN (chr10:g.87957873C>T, ENSP00000361021.3:p.Gln219Ter) | GBM, IDHwt, RTK I/II | methylated |
| T384 | LIH0384 | 50 | Female | Temporo-Occipital left | primary | no prior treatment | GBM Grade IV | organoids | G4 | 202+/-6,88 | NA | no detectable | NA | ++[EGFR, 7p15.2-p11.2, 7q31.31-32.2], -[4p, 9q, 10q, partial Chr 12, partial Chr 15], --[FGFR1\TACC1, MYBL1] | TP53 (chr17:g.7674257A>T, ENSP00000269305.4:p.Tyr236Asn) | GBM, IDHwt, RTK II (no match) | methylated |
| T386 | LIH0386 | 51 | Male | Frontal left | primary | no prior treatment | GBM Grade IV | organoids | G4 | 104,1+/-6,83 | 28.9+/-0.72 | detectable, quantifiable | 2.1N | +[Chr7, 19p], - [1p36.3-p34.3, Chr10, 11q12.3-q13.2, 11q24.1-q13.2, 14q11.2-q24.1, 17q11.2-q21.2, 18p11.32-p11.22, 22p ], --[CDKN2A/B, PTEN] | PIK3R1 (chr5:g.68293801_68293803del, ENSP00000323512.8:p.Asp164_Arg165delinsGlu)<br>NF1(chr17:g.31221932A>G, ENSP00000412921.4:p.Tyr575Cys) likely not pathogenic | GBM, IDHwt, RTK I | unmethylated |
| T394 | LIH0394 | 45 | Female | Fronto-basal right | recurrence 1 | radiotherapy | GBM Grade IV | fresh organoids/tissue | G5 | 68,56+/-0,9 | 67.8+/-8.3 | detectable, quantifiable | 2.1-2.3N + 3.5N | complex genome ++[PDGFRA,FGFR3/TACC3, KIT, KDR, MET, Chr 4, 7q31.31], --CDKN2A/B | IDH1 (chr2:g.208248388C>T, ENSP00000260985.2:p.Arg132His)<br>MET (chr7:g.116700208A>G, ENSP00000317272.6:p.Asn375Ser)<br>TP53 (chr17:g.7670697_7670707del, ENSP00000269305.4:p.Glu336AspfsTer7)<br>ATRX (chrX:g.77681626_77681628del, ENSP00000362441.4:p.Asp1212del)<br>ATRX (chrX:g.77682604_77682607del, ENSP00000362441.4:p.Glu886LeufsTer18) | Glioma, IDHm, high grade astrocytoma | methylated |
| T407 | LIH0394 | 45 | Female | Fronto-basal right | recurrence 1 | radiotherapy | GBM Grade IV | fresh organoids/tissue | G6 | 64+/-4,27 | 49.5+/-6.8 | detectable, quantifiable | 2.1N + 3.9N | complex genome ++[PDGFRA,FGFR3/TACC3, KIT, KDR, MET, Chr 4, 7q31.31], --CDKN2A/B | IDH1 (chr2:g.208248388C>T, ENSP00000260985.2:p.Arg132His)<br>MET (chr7:g.116700208A>G, ENSP00000317272.6:p.Asn375Ser)<br>TP53 (chr17:g.7670697_7670707del, ENSP00000269305.4:p.Glu336AspfsTer7)<br>ATRX (chrX:g.77681626_77681628del, ENSP00000362441.4:p.Asp1212del)<br>ATRX (chrX:g.77682604_77682607del, ENSP00000362441.4:p.Glu886LeufsTer18 | Glioma, IDHm, high grade astrocytoma | methylated |
| T434 | LIH0304 | 54 | Male | Frontal left | recurrence 2 | radiotherapy + TMZ | GBM Grade IV | organoids | G6 | 43,20+/-0 | 40.4 +/- 11.43 | detectable, quantifiable | 2.0N + 2.3N +3.8N | ++[EGFR, CDK4, MDM2], +[7p,7q36.1-q36.2, Chr20], -[1p36.23-p36.21, 6q26-q27, Chr10, 12q15, 14q13-qter] | EGFR (chr7:g.55154023T>A, ENSP00000275493.2:p.Phe254Ile)<br>PTEN (chr10:g.87933166G>A, ENSP00000361021.3:p.Cys136Tyr) | GBM, IDHwt, RTK II | unmethylated |
| T470 | LIH0347 | 42 | Male | Frontal left | recurrence 1 | radiotherapy + TMZ | GBM Grade IV | organoids | G4 | 78,33+/-15,27 | 17.98 +/- 0.88 | detectable, quantifiable | 2.2N | ++EGFR, +[Chr7, 19p, Chr20], - [ 1p36.3-p34.3, 1p31.1-p11.21, 1q32.3-q44, Chr10, 13q12.11-31.1, 14q11.2-q21.3], --CDKN2A/B | MET (chr7:g.116700208A>G, ENSP00000317272.6:p.Asn375Ser)<br>PTEN (chr10:g.87961003del, ENSP00000361021.3:p.Cys304SerfsTer3) | GBM, IDHwt, RTK II | unmethylated |
| T476 | LIH0476 | 75 | Male | Fronto-parietal left | primary | no prior treatment | GBM Grade IV | organoids | G4 | 66+/-0 | 44.3+/-3.8 | detectable, non quantifiable | 2.3N | ++[EGFR, MDM4], +[Chr 7], - [9p24.3-p21.3, Chr 10], --[CDKN2A/B, PTEN] | EGFR (chr7:g.55165350G>T, ENSP00000275493.2:p.Gly598Val)<br>EGFR (chr7:g.55200333G>C, ENSP00000275493.2:p.Asp956His)<br>MET (chr7:g.116771936C>T, ENSP00000317272.6:p.Thr1010Ile) | GBM, IDHwt, RTK I | unmethylated |
| T515 | LIH0515 | 46 | Male | Frontal-right | primary | no prior treatment | Anaplastic Oligodendroglioma Grade III | fresh organoids/tissue | G5 | 393,79+/-55,56 | NA | detectable, quantifiable | NA | NA | NA | NA | NA |
| T591 | LIH0384 | 52 | Female | Temporo-Occipital right | recurrence 1 | radiotherapy +TMZ + Avastin + Irinotecan | GBM Grade IV | organoids | G3 | 110+/-16 | 22.08+/-12.72 | detectable, quantifiable | 2.2N + 3.5N | ++[EGFR, 7p15.2-p11.2, 7q31.31-32.2], -[1p21.1, 3q26.1, 8p, 9p24.3, Chr 10, 11q24.2, 13q21.33-q31.1, 15q] | No Glioma-specific mutations | GBM, IDHwt, RTK II/I | unmethylated |
| T744 | LIH0744 | 69 | Male | Bilateral involvement of the hippocampus | primary | no prior treatment | GBM Grade IV | organoids | G3 | 100+/-0 | 28.9+/-23.5 | detectable, quantifiable | 2.2N | +Chr 7, -[6q, 9p, Chr10, Chr 11, 14q, 22q], --CDKN2A/B | NF1 (chr17:g.31338092C>T, ENSP00000348498.3:p.Arg2237Ter) | GBM, IDHwt, RTK I/II | unmethylated |
| T756 | LIH0556 | 46 | Male | Multifocal left | recurrence 3 | radiotherapy | GBM Grade IV | fresh organoids/tissue | G5 | 77,93+/-1,42 | 12.09 +/-1.62 | detectable, quantifiable | 2.2N + 2.9N + 3.9N | + [4q12, 10p], -[4p15.32-q35.2, 6q27, 10q, partial Chr 16, 22q], --CDKN2A/B | IDH1 (chr2:g.208248388C>T, ENSP00000260985.2:p.Arg132His)<br>PTCH1 (chr9:g.95469004G>A, ENSP00000332353.6:p.Thr666Met)<br>TP53 (chr17:g.7675085C>T, ENSP00000269305.4:p.Cys176Tyr) | Glioma, IDHm, high grade astrocytoma | methylated |
| T772 | LIH0615 | 54 | Female | Fronto-temporal left | recurrence 1 | radiotherapy + TMZ | GBM Grade IV | organoids | G4 | 107+/-0 | 32.1+/-1.2 | detectable, non quantifiable | 2.1N | ++EGFR, +[7p, Chr19], -[1p36.3, 9p, Chr 10, 15q] --CDKN2A/B | EGFR (chr7:g.55165350G>T, ENSP00000275493.2:p.Gly598Val)<br>RET (chr10:g.43124887C>T, ENSP00000344798.4:p.Arg982Cys)<br>PTEN (chr10:g.87961115del, ENSP00000361021.3:p.Phe341LeufsTer3) | GBM, IDHwt, RTK I | methylated |
| T784 | LIH0784 | 56 | Male | Frontal left | primary | no prior treatment | GBM Grade IV | organoids | G4 | 138+/-2 | 43.5+/-3.3 | detectable, non quantifiable | 2.1N | +[Chr 7], -[1p36.3-22.1, 3p12.1, 10q], --CDKN2A/B | MET (chr7:g.116695757A>T, ENSP00000413857.1:p.Met1?)<br>RET (chr10:g.43102379C>A, ENSP00000491505.1:p.Leu107Ile)<br>RB1 (chr13:g.48381295G>A, ENSP00000267163.4:p.Trp516Ter)<br>STAG2 (chrX:g.124094127dup, ENSP00000218089.9:p.Thr1230AsnfsTer21) | GBM, IDHwt, RTK I | methylated |
| T797 | LIH0797 | 55 | Female | Left occipital | primary | no prior treatment | GBM Grade IV | organoids | G3 | 119+/-34,5 | 9.65 +/-0.40 | detectable, quantifiable | 2.2N + 3.9N | complex genome ++[CDK4, MDM2], +[Chr 7, Chr 20], -[Chr 10, 13q, 15q, 22q] | NA | GBM, IDHwt, RTK I | methylated |
| T831 | LIH0831 (n | 51 | Female | Bifrontal | primary (multifo | no prior treatment | GBM Grade IV | organoids | G3 | 70+/-0 | 3.46 +/- 2.33 | NA | NA | ++EGFR, +[Chr 7], -[3p12.1, 6p24.2, Chr 10, 13q, 16p12.2], --CDKN2A/B | PIK3CA (chr3:g.179199048C>G, ENSP00000263967.3:p.Gln75Glu) | GBM, IDHwt, RTK II/I | methylated |
| T832 | LIH0831 (n | 51 | Female | Bifrontal | primary (multifo | no prior treatment | GBM Grade IV | organoids | G3 | 138,5+/-14,50 | 0.25 +/- 0.04 | no detectable | 2.1N | NA | NA | NA | NA |
| T841 | LIH0841 | 38 | Male | Right temporal | primary | no prior treatment | GBM Grade IV | organoids | G3 | 49,2+/-1,09 | NA | NA | NA | complex genome ++[MYCN, CDK4], +Chr 7, -[2p24.2, 6q12, Chr 10, 13q31.2, 14q12, 15q15.3] | PIK3CA (chr3:g.179199156_179199161del, ENSP00000263967.3:p.Lys111_Ile112del)<br>MET (chr7:g.116695757A>T, ENSP00000413857.1:p.Met1?)<br>CDKN2A (chr9:g.21970917C>T, ENSP00000307101.5:p.Ala148Thr) | Glioblastoma, Idh Wildtype, With Primitive Neuronal Component (No match) | unmethylated |
| T861 | LIH0841 | 38 | Male | Right temporal | recurrence 1 | radiotherapy + TMZ | GBM Grade IV | organoids | G2 | 157,2+/-3.88 | NA | detectable, non quantifiable | NA | NA | NA | NA | NA |
| T899 | LIH0609 | 75 | Female | Right Central | recurrence 1 | no prior treatment | GBM Grade IV | organoids | G2 | 70,75 + /- 2,63 | NA | NA | NA | complex genome ++[PDGFRA, CDK4], -[Chr10, 14q] | PIK3R1 (chr5:g.68296465A>C, ENSP00000513328.1:p.Gln235His)<br>PTCH1 (chr9:g.95468828G>A, ENSP00000332353.6:p.Pro725Ser)<br>CDK4 (chr12:g.57749225G>A, ENSP00000257904.5:p.Ser259Leu)<br>TP53 (chr17:g.7674188C>A, ENSP00000269305.4:p.Asp259Tyr) | Diffuse Pediatric Type High Grade Glioma, H3 Wildtype And Idh Wildtype, RTK I (No match) | methylated |
| T905 | LIH0609 | 76 | Female | Right Central | recurrence2 | no prior treatment | GBM Grade IV | organoids | G3 | 91+/-1,73 | NA | detectable, non quantifiable | NA | NA | NA | NA | NA |
| T1053 | LIH1053 | 62 | Male | Right intraaxial | primary | no prior treatment | GBM Grade IV | organoids | G2 | 98.75+/-0.5 | NA | NA | NA | NA | NA | NA | NA |
| T1070 | LIH0784 | 60 | Male | Left frontal | recurrence 1 | radiotherapy + TMZ | GBM Grade IV | organoids | G1 | 254.25 +/- 36.18 | NA | detectable, quantifiable | NA | NA | NA | NA | NA |

**Table S2.** List of antibodies used in the study.
*Flow cytometry test $10^6$ cells/100µl

| Antibody | Supplier | Catalog number | Concentration used |
|---|---|---|---|
| hCD90 PE-Cy7 | BD | 561558 | 5 µl/test* |
| hCD90 BV605 | BD | 562685 | 5 µl/test* |
| CD31 Dy590 (PE-TR) | Immunotools | 21270317.5 | 10µl/test* |
| CD45 PE-Cy7 | BD | 557748 | 5µl/test* |
| CD16/32 | eBioscience | 14-0161-85 | 1 µl/test* |
| mCD45 FITC | eBioscience | 11-0454-82 | 1 µl/test* |
| mCD11b Percp-Cy5.5 | BD | 550993 | 5 µl/test* |
| mLy6C PB | Biolegend | 128014 | 0.5 µl/test* |
| mLY6G BV785 | Biolegend | 127645 | 2.5 µl/test* |
| mCCR2 PE | R&D | FAB5538P | 5 µl/test* |
| iso IgG PE | R&D | IC108P | 5 µl/test* |
| mCD206 APC | Biolegend | 141708 | 2.5 µl/test* |
| mCD11c APC-Cy7 | Biolegend | 117323 | 5 µl/test* |
| mCD86 BV605 | Biolegend | 105037 | 5 µl/test* |
| iso IgG2a BV606 | Biolegend | 400540 | 5 µl/test* |
| m I-A/I-E APC-Cy7 | Biolegend | 107628 | 1 µl/test* |
| mCD45 BV605 | Biolegend | 11045182 | 1 µl/test* |
| LIVE/DEAD™ Fixable Near-IR | Invitrogen | L34975 | 0.1 µl/test* |
| Nestin | Abcam | AB6320 | IHC:1/500 |
| Vimentin | Thermo Fisher Scientific | Mab3400 | IHC:1/200 |
| CD31 | Cell Signaling | 77699 | IHC:1/100 |
| Iba1 | Biocare Medical | CP 290A | IHC:1/1000 |
| Iba1 | Wako | 019-19741 | IHC:1/1000 |
| CD11c | Abcam | ab11029 | IHC:1/100 |
| GFAP | Dako | Z0334 | IHC:1/1000 |
| MHC-II | Abcam | ab25333 | IHC:1/100 |
| Ki-67 | ThermoScientific | 14-5698-82 | IHC: 1/100 |
| Pdgfra | Biolegend | 135905 | IHC:1/100 |
| Anti-Rat IgG Alexa Fluor 555 | Invitrogen | A21434 | IHC:1/1000 |
| Anti-Rabbit IgG Alexa Fluor 555 | Thermo Fisher Scientific | A-11039 | IHC:1/500 |
| Anti-Mouse IgG Alexa Fluor 647 | Thermo Fisher Scientific | A-11037 | IHC:1/500 |
| Anti-Rat IgG Alexa Fluor 488 | Thermo Fisher Scientific | A-21244 | IHC:1/500 |
| Opal 520 Flurophore | Akoya Biosciences | NEL810001KT | IHC:1/100 |
| Opal 570 Flurophore | Akoya Biosciences | NEL810001KT | IHC:1/100 |

**Table S3.** Characteristics of cell clusters identified in DROP-seq data in mouse-derived TME. Marker genes for each cluster were defined as differentially expressed genes between clusters at threshold $log_2FC>=0.5$
Genes are listed from the highest to lowest $log_2FC$

| Cluster number | Astrocytes | Myeloid | Endothelial | OPCs | Oligodendrocytes | Cycling | Ependymal | Pericytes | Lymphocytes |
|---|---|---|---|---|---|---|---|---|---|
| | Slc1a2 | Ctss | Ly6c1 | Cacng4 | Plp1 | Hist1h4c | Ttr | Ccdc153 | Ccl5 |
| | Aqp4 | Hexb | Cxcl12 | Pdgfra | Ermn | Top2a | Enpp2 | Tmem212 | Nkg7 |
| | Gja1 | Lyz2 | Cldn5 | Olig1 | Mbp | Gm14892 | 1500015O10Rik | Rarres2 | Ms4a4b |
| | Aldoc | C1qb | Itm2a | Rtn1 | Mag | Cenpf | Prlr | Rsph1 | Trac |
| | Slc1a3 | Cx3cr1 | Igfbp7 | Cntn1 | Ptgds | Gm12331 | Calml4 | Enkur | Gzmb |
| | Bcan | Cd74 | Rgs5 | Gng3 | Cldn11 | Prc1 | Igf2 | Fam183b | Trbc2 |
| | Ttyh1 | Tyrobp | Bsg | Lhfpl3 | Mobp | Ezh2 | Kl | Ak7 | Cd3g |
| | Slc4a4 | C1qa | Flt1 | 3110035E14Rik | Mal | Smc4 | Kcnj13 | Mns1 | Trbc1 |
| | Clu | P2ry12 | Abcb1a | Cspg5 | Cnp | Gm17909 | Cab39l | Dynlrb2 | Gimap4 |
| | Plpp3 | Spp1 | Slco1a4 | Ccnd1 | Stmn4 | RP24.232I9.2 | Stk39 | Spef2 | Klrd1 |
| | Atp1a2 | C1qc | Spock2 | Pcsk1n | Apod | Gm12152 | Igfbp2 | Cfap44 | Cd3d |
| | Scg3 | Ctsd | Adgrf5 | Nap1l5 | Tspan2 | Smc2 | Trpm3 | Ccdc146 | Gimap3 |
| | Acsl3 | Csf1r | Ly6a | Meg3 | Aplp1 | Gm12504 | Folr1 | Wdr66 | Ptpn22 |
| | Pla2g7 | Laptm5 | Ctla2a | Tnr | Car2 | Rps19.ps4 | Cdkn1c | Rsph4a | P2ry10 |
| | Htra1 | H2.Ab1 | Car4 | Ncald | Trf | Gm5853 | Chchd10 | Calml4 | Ptprcap |
| | Cldn10 | Cxcl13 | Pltp | S100a1 | Qdpr | Cenpe | Sostdc1 | Cfap126 | Cytip |
| | Tspan7 | Fcer1g | Ptprb | Cdo1 | Gm21984 | Ube2c | Clic6 | Cfap45 | Sept1 |
| | Id4 | Mpeg1 | Egfl7 | Gap43 | Sept4 | Gm12165 | Rbp1 | Iqcg | Lck |
| | Acsl6 | Siglech | Palmd | Fabp7 | Pex5l | Gm18014 | Otx2 | Fam216b | H2.Q7 |
| | Ntsr2 | H2.Aa | Ramp2 | Olig2 | Mog | Gm14537 | Pcp4 | 4933434E20Rik | Thy1 |
| | Gpr37l1 | Trem2 | Gng11 | Scg5 | Opalin | Tpx2 | Tmem72 | Spa17 | Mndal |
| | Sparcl1 | Ccl4 | Adgrl4 | Vcan | Tmem88b | Gm13121 | Ppp1r1b | Gm45396 | AW112010 |
| | Dclk1 | Clec7a | Kitl | Sulf2 | Ugt8a | Gm4321 | F5 | Ccdc39 | Nr4a2 |
| | Mt3 | Itgam | Esam | Epn2 | Il33 | Nusap1 | Fam81a | Meig1 | Stk17b |
| | Atp1b2 | Aif1 | Ablim1 | Matn4 | Ttll7 | Gm17511 | Ahcyl2 | Ccdc113 | S100a4 |
| | Gpm6a | Il1b | Slc39a10 | Anks1b | Ptprd | RP23.390K13.4 | Cul4b | Gm973 | Gimap6 |
| | Cpe | Gpr34 | Arhgap29 | Bex2 | Gjc3 | Gm44010 | Vat1l | Lrrig1 | Ptprc |
| | Sox2 | Ctsb | Tm4sf1 | Pllp | Anln | Birc5 | Car12 | Ppp1r36 | Vps37b |
| | Id2 | Hpgds | Col4a2 | Stmn3 | Epb41l3 | Gm11223 | Hemk1 | Wdr78 | Plac8 |
| | Luzp2 | Ccl3 | Pcp4l1 | Gria4 | Ccp110 | 2810417H13Rik | Sulf1 | 1110017D15Rik | Il2rg |
| | S1pr1 | Lgmn | Sdpr | Kcnip3 | Gnai1 | Kif15 | Ociad2 | Mia | Ifi203 |
| | Ptprz1 | Rnase4 | Slco1c1 | Scrg1 | Tubb4a | Gm8790 | Spint2 | Dnah12 | Hcst |
| | Ndrg2 | Lpl | Pecam1 | Il1rap | Aspa | Gm44122 | Msx1 | Spag17 | S100a10 |
| | Mfge8 | Fyb | Uaca | Gjc3 | Spock3 | Hmmr | Rsph1 | 1700016K19Rik | Rps18 |
| | Nrxn1 | Tgfbr1 | Nostrin | Gria3 | Tppp | RP23.144N15.3 | Fhad1 | Foxj1 | Rps15a |
| | Tmem47 | Ctsz | Abcg2 | Bcas1 | Map7 | Ncapg | Slc4a10 | Cd24a | Rpl34 |
| | Gria2 | Grn | Col4a1 | Tuba1a | Slc24a2 | Gm7860 | Kcne2 | Tppp3 | H2.K1 |
| | Mt2 | Selplg | Ifitm3 | Serpine2 | Lpar1 | Bub1 | Htr2c | Sntn | Rps24.ps2 |
| | Mt1 | Ccl6 | 9430020K01Rik | Cspg4 | Cntn2 | Kif23 | Atp2b3 | 1700012B09Rik | Cd52 |
| | Lsamp | Plek | Hspb1 | Phlda1 | Pllp | Nuf2 | Col9a3 | Tmem107 | H2.Q6 |
| | Ntrk2 | Fcgr3 | Tsc22d1 | Dbi | Mapt | Gm5070 | Emb | Riiad1 | Gm11478 |
| | Ednrb | Ly86 | Slc9a3r2 | S100a16 | Bcas1 | Mki67 | Drc7 | Ccdc180 | AU020206 |
| | Prnp | Rgs2 | Pglyrp1 | Ednrb | Sox2ot | Rps12.ps19 | Cox6b2 | Cfap43 | Gm6030 |
| | Ckb | Apbb1ip | Cp | Tmem100 | Edil3 | Tubb3 | Fam161a | Drc1 | Rac2 |
| | Gpm6b | Cd68 | Itih5 | Gria2 | Gjb1 | Gm28344 | Lrrig1 | Iqca | Rps3 |
| | Ptn | Rgs1 | Kdr | Nnat | Desi1 | Gm3788 | Mlf1 | Bphl | Srgn |
| | Pou3f3 | Cd52 | Nes | Ptprz1 | Gpr37 | Gm8513 | Krt18 | Cep126 | Ms4a6b |
| | Prdx6 | Mafb | Sptbn1 | Nav1 | Ndrg1 | Rpl21.ps5 | Rdh5 | Mlf1 | Gm15427 |
| | Cspg5 | Rgs10 | Slc2a1 | Syt11 | Olig1 | Gm2522 | Plppr4 | Fhad1 | Rps2 |
| | Gstm1 | Slc15a3 | Rflnb | Phactr3 | Dusp26 | RP23.408J7.2 | Col8a1 | Cfap54 | Rplp1 |
| | Cxcl14 | Olfml3 | Ccdc141 | Nrxn2 | Map1a | Gm2178 | Tbc1d9 | Odf3b | Tmsb10 |
| | Ldhb | Atf3 | Myo1b | Map2 | Grb14 | Nasp | Car14 | Dnali1 | Rps24 |
| | Fabp7 | Npc2 | Tpm1 | Gpm6b | Enpp2 | Rpl26.ps2 | Atp1b1 | Cfap161 | Gm9794 |
| | Syt11 | Zfp36 | Ece1 | Rpl13 | Scd2 | Rpl34.ps2 | Tmem237 | 1500015O10Rik | Rps15a.ps5 |
| | Timp3 | Man2b1 | Utrn | Myo5a | Sez6l2 | Acta1 | Gpx8 | Elof1 | Wdr89 |
| | Fam213a | Cd14 | Tspan13 | Lbh | Plekhb1 | Gm11810 | Sh3d19 | Dnah6 | Rpl39 |
| | Syne1 | Slc11a1 | Slc16a1 | Ttc3 | Hapln2 | Gm11517 | Ctnnal1 | Ccdc173 | Rps14 |
| | Asrgl1 | Cst7 | Ets1 | Ahi1 | Elavl3 | Rpl10l | Arsg | Lrrc23 | Gm10275 |
| | Glul | Irf8 | Myl12a | Tspan3 | Ank3 | Rps12.ps24 | Acad8 | Ascc1 | Rps13.ps2 |
| | Scd2 | Pld4 | Klf2 | Asrgl1 | Trim59 | Gm7351 | Slc22a17 | Ak9 | B2m |
| | Luc7l3 | Golm1 | Esyt2 | Fip1l1 | Ppp1r14a | Gm2225 | Pcp4l1 | Hydin | Shisa5 |
| | Tspan3 | Creg1 | Igf1r | Hip1 | Pls1 | Tubb5 | Fxyd1 | Cfap100 | Rps26 |
| | Hnrnpa2b1 | Cyth4 | Tmsb10 | Basp1 | Slc44a1 | Gm13215 | Gsta4 | Dnaic1 | Rpl32 |
| | Kif1b | Unc93b1 | Tpm4 | Rev3l | Slc48a1 | Gm7558 | Acaa2 | Dnah5 | Rpsa.ps10 |
| | Apc | Ptprc | BC028528 | Arl2bp | Efnb3 | Tmpo | Sfrp1 | Nme5 | Samd9l |
| | Ccdc88a | Maf | Gm10080 | S100b | Cmtm5 | Gm5370 | Gm17018 | Ttc21a | Ets1 |
| | Glud1 | Ctsa | Ptprg | Ccnd2 | Kcna1 | Gm15896 | Mrps6 | Hdc | Rps24.ps3 |
| | Serpine2 | Ctsc | Epas1 | Scd2 | Fa2h | Gm11472 | Mcee | Ccdc81 | Gm9843 |
| | Zbtb20 | Itgb5 | Akap12 | Ppfibp1 | Mapk8ip1 | Gm15159 | Atp5g1 | Efcab1 | Gm15500 |
| | Spag9 | Cyba | Ppfibp1 | Slc25a4 | Nfasc | Gm9625 | Rabgap1l | Nme9 | Ifngr1 |
| | Id3 | Plxdc2 | Crip2 | Hmgcs1 | Aatk | Rpl17.ps4 | Phactr2 | Elmod1 | Rps15 |
| | Macf1 | Mef2c | Vwa1 | Nrxn1 | Sept8 | Rpl27.ps2 | Tubb4b | Akap14 | Rplp0 |
| | Sept7 | Cd53 | Myl12b | Tmem176b | Lgi3 | Gm14277 | Car2 | Rsph10b | Rpl22 |
| | Dst | Ptgs1 | Arl4a | Tpm1 | Tmeff2 | Rpl13.ps5 | Mdh1 | Rgs22 | Rpl4 |
| | Vim | Cd84 | Golim4 | Psip1 | Cpox | Gm6543 | Bphl | Gm28729 | Rpl35a |
| | Malat1 | Cybb | Ly6e | Ramp1 | Tppp3 | Lmnb1 | Map1lc3a | Efcab2 | Rps20 |
| | mt.Rnr1 | Lair1 | App | Zeb1 | Dnajb2 | Incenp | Ift74 | 3300002A11Rik | Rps5 |
| | Qk | Ccr5 | Itgb1 | Pcdh17 | Dixdc1 | Nucks1 | Slc16a2 | Ttc25 | H2.D1 |
| | Pnisr | Hexa | Sparc | Atxn7l3b | Josd2 | Gm14450 | Prdx5 | 1700007K13Rik | Gm5963 |
| | Fermt2 | Fcgr1 | Id3 | Lsamp | Nkx6.2 | Gm14480 | Gm10039 | Armc3 | Slfn2 |
| | Arglu1 | Cst3 | Calm1 | Pcdh10 | Shtn1 | Rrm1 | Tbcel | Cfap69 | Rps21 |
| | Son | Junb | Sept4 | Pebp1 | Arsg | H2afv | Maob | 1700024G13Rik | Eef1b2 |
| | Arhgap5 | Cd83 | Atox1 | Prdx2 | Sox10 | Rps3a3 | Mtch1 | Lrrc51 | Rplp2 |
| | Rac1 | Kctd12 | Crip1 | Ndufa4 | Sv2a | Gm12857 | Hist1h2bc | 1700088E04Rik | Lcp1 |
| | Rtn3 | B2m | Id1 | Klhl5 | Cryab | Rad21 | 1110004E09Rik | Tctex1d4 | Rps19 |
| | Tsc22d4 | Ncf1 | Cald1 | Kifap3 | Pcdh9 | Gm29228 | Thyn1 | Syt10 | Rps3a1 |
| | Tcf4 | Lpcat2 | Fxyd5 | Luzp2 | Anks1b | Gm11560 | Elovl7 | Efcab10 | Gm5805 |
| | Ywhae | Smap2 | Luzp1 | Atp1b1 | Plcl1 | Fth.ps2 | Tgfb2 | Armc4 | Rpl37rt |
| | Serinc1 | Apoe | Col4a3bp | Alcam | Tmod2 | Gm7618 | Gas6 | Capsl | Rpl3.ps1 |
| | Slc25a4 | Cd37 | Slc6a6 | Hsd17b12 | Slc12a2 | Gm6136 | Ifi27 | Lrrc48 | Rps16.ps2 |
| | Hnrnpdl | Nfkbiz | Cd2ap | Ifi27 | Agpat4 | Bex3 | 1110032A03Rik | Cdhr3 | Gm6472 |
| | Ktn1 | Ms4a6c | Fnbp1l | Cd9 | Gpm6b | Gm12482 | Hopx | Lrrc6 | Tpt1.ps3 |
| | Gm42418 | Lat2 | Myl6 | Ubb | Syt11 | Rps19.ps6 | Ltc4s | Ccdc30 | Gm5905 |
| | Eif5 | Lcp1 | Fermt2 | Eif1b | Gsn | Gm13815 | Cd59a | Dnah3 | Rpl9.ps6 |
| | | Samsn1 | Cd34 | Atpif1 | Fez1 | Gm13340 | Ndufa11 | Tekt1 | S100a11 |
| | | Arsb | H2.D1 | Hsp90aa1 | Dnm3 | Gm7308 | Ppfibp2 | Maats1 | Rps9 |
| | | Gns | Bnip2 | Ptn | Rtn4 | Gm15773 | Thumpd3 | Nek5 | Rpl3 |
| | | Sgk1 | Cyb5r3 | Nme1 | Csrp1 | Hmgb2 | Ndufv3 | Cfap46 | Cnn2 |
| | | Egr1 | Sept11 | Chpt1 | App | Hnrnpa2b1 | Cox20 | Lca5 | Gm14513 |
| | | Pycard | Srgn | Xist | Trim2 | Atad2 | Usmg5 | Crygn | Rps15a.ps7 |
| | | Gusb | Sh3bgrl | Ankrd11 | Ywhaq | Rbmx | Ldhb | Pifo | Gm6204 |
| | | Cotl1 | Rpl38.ps2 | Cadm2 | Elovl7 | Selenoh | Cox7b | Nnat | Gm8995 |
| | | Btg2 | Ctnna1 | Ncam1 | Otud7b | Gm27046 | Ndufa1 | Chchd10 | Rpl14.ps1 |
| | | Abhd12 | Hes1 | Ckb | Pak1 | Gm2000 | Cox6c | Ift88 | Rps11 |
| | | Ifi27l2a | Dynll1 | Lmo4 | Fnbp1 | Ddx39 | Uqcr10 | Fam81a | Ahnak |
| | | Lyn | Gm4617 | Tubb5 | Shisa4 | Vim | Anapc13 | Fhl1 | Rps10.ps1 |
| | | Jun | Shfm1 | Fubp1 | Secisbp2l | Hnrnpd | Glrx3 | Dnal1 | Rpl18.ps1 |
| | | Basp1 | Fn1 | Smarcc1 | Kif1b | Ybx1 | Cetn2 | Nuf2 | Rps10 |
| | | Ctsh | Anp32a | Hnrnpr | Gatm | Rps15.ps2 | Atp11c | Chek2 | Btg1 |
| | | AU020206 | Aplp2 | Hspd1 | Tulp4 | Gm16379 | Idh3a | Ccdc34 | Gm16580 |
| | | Pmepa1 | Selenop | Bzw1 | Chn2 | Gm4204 | Atp6v1a | Slc38a1 | Pabpc1 |
| | | Sirpa | Iqgap1 | Tsc22d4 | Kif21a | RP23.114G13.7 | Etfa | 2610301B20Rik | Gm5835 |
| | | Syngr1 | Nedd4 | Srrm2 | Spg20 | Gm5778 | Gm12669 | Nudt4 | Psmb8 |
| | | Psap | Nrp1 | Ppp2r2a | Cdc37l1 | Psip1 | Cgnl1 | Hspa4l | Rpl8 |
| | | Serinc3 | Srpr | Nedd4 | Sirt2 | Dnmt1 | Pebp1 | Prelp | Rpsa |
| | | Ier5 | Atp5j | Anp32e | Bin1 | Hspd1.ps3 | Vamp8 | Rassf9 | Crip1 |
| | | Rhob | Kif5b | Top2b | Inpp5f | Gm13498 | Dynlt3 | Fam161a | Arhgap30 |
| | | Ctsl | Ubb | Ostf1 | Sccpdh | Pcna.ps2 | Ezr | Gm5617 | Rps26.ps1 |
| | | Arhgdib | Gnai2 | Ccdc88a | Psat1 | Ssrp1 | Atp5h | Wdr60 | Gm4604 |
| | | Bin1 | Chchd2 | Tecr | Gstp.ps | Calm2 | Gm16418 | Ift74 | Lgals1 |
| | | Lgals3bp | Pomp | Agpat5 | Sept7 | Gm10167 | Arl6ip1 | Aebp1 | Rpl26 |
| | | H2.K1 | Slc3a2 | Pnn | Arpc1a | Gm12222 | 2010107E04Rik | Cetn2 | Gm4149 |
| | | Ivns1abp | Gm6472 | Eid1 | Osbpl1a | Anp32e | Uqcr11 | Ift81 | Rack1 |
| | | Efhd2 | Gm9800 | Hsbp1 | Neat1 | Dek | Aldh2 | 1110032A03Rik | Rpl30 |
| | | Bmp2k | Gm14586 | Ybx1 | Rnf13 | Ccdc34 | Hint2 | Dbi | Ifi27l2a |
| | | Arpc1b | Gm11966 | Rab2a | Cers2 | Gm11488 | Hcfc1r1 | Cep290 | Sp100 |
| | | Cd9 | 1810037I17Rik | Ndufa5 | Erbin | Gm4332 | Cox8a | Hsp90aa1 | Gm10288 |
| | | Scamp2 | Laptm4a | Tial1 | Fam171b | Anp32b.ps1 | Ndufa4 | Nphp1 | Rpl41 |
| | | Timp2 | Nfkbia | Dhx36 | Dbi | Rn7s6 | Gm16399 | Rpgrip1l | Hspa8 |
| | | Rin2 | Msn | Dnaja1 | Pip4k2a | Ilf3 | Rnf7 | Strbp | Rps11.ps2 |
| | | Rrbp1 | Ndufa7 | Ywhab | Gm11223 | Gm22567 | Atp5o | Nin | Gm14303 |
| | | Gm14303 | Psma7 | Cox6a1 | Evi2a | Pa2g4 | Atp5d | Tubb4b | S100a6 |
| | | Fam49b | Tpt1.ps3 | Hnrnpk | Cdc42bpa | Anapc11 | Etfb | 6330403K07Rik | Fxyd5 |
| | | Epb41l2 | H3f3b | Thoc7 | Degs1 | Snrpd1 | Ap1s2 | Hsph1 | Gm4332 |
| | | Cfh | Ptma | Nap1l1 | Nap1l5 | Ranbp1 | Gm10250 | Gm14964 | Rps11.ps1 |
| | | Mef2a | Gm10250 | Dynll2 | Map7d1 | Ybx1.ps2 | Ndufb10 | Fam149a | Sh3bgrl3 |
| | | Mertk | Gm4149 | Cox6c | Tuba1a | Gm6159 | Arl3 | Pcp4l1 | Gas5 |
| | | Rnaset2a | Spop | Cacybp | Efcab14 | Gm9294 | Cox4i1 | Rdh10 | Rpl39.ps |
| | | H2.D1 | Sh3glb1 | Wbp5 | Tsc22d4 | Dnajc9 | Atp5g3 | Zcchc18 | Rpl23 |
| | | Erp29 | Cox6a1 | Hnrnpc | Pla2g16 | Gm13461 | Cisd1 | Gm12346 | Eef1a1 |
| | | Gpx1 | Rpl41 | Rpl13a | Gltp | Gm11942 | Pgrmc1 | Hipk1 | Rpl38.ps2 |
| | | Marcks | Anxa5 | Ran | Ppp2r3a | Hdac2 | Acadsb | Nek1 | Pdcd4 |
| | | Tmsb4x | Snx3 | Pcm1 | Gstp1 | Gm8186 | Slc31a1 | 1110004E09Rik | Jak1 |
| | | Actb | Cdc42 | Rrp1 | Taldo1 | Gm13092 | Slc12a2 | Ccdc191 | Rps10.ps2 |
| | | Itm2b | Rpl37rt | Arl6ip1 | Sec11c | Gm6394 | Polr2m | Tsga10 | Gm9385 |
| | | Rplp1 | Cbx1 | Zmynd11 | Rnf7 | Gm11633 | Ndufb2 | Gm12222 | Rpl10a.ps1 |
| | | Pabpc1 | Slk | Hnrnpab | Kif1a | Snrpe | Ndufc1 | Ccdc181 | Psmb9 |
| | | Fth1 | Timp3 | Cox6b1 | Epn2 | Ppm1g | Dstn | Ildr2 | Rpl37 |
| | | Sparc | Cox6c | Rab10 | Mbnl2 | Snrpd2 | Ccdc34 | Tm4sf1 | Cntrl |
| | | Trf | Tpm3.rs7 | Sec11c | Ncam1 | Hist1h1e | Atp5j | Ipo11 | Junb |
| | | Rps24 | Dnajc8 | Acat1 | Thra | Hnrnpr | Gm14539 | Wwc1 | Npm1 |
| | | Skil | Rps3a1 | Ndufc2 | Ank2 | Hnrnpab | Ccng1 | Hspa2 | H2.T23 |
| | | Zeb2 | Rps19 | Hmgn1 | Dpysl2 | Gm12020 | Ndufa13 | Cep83 | Gm8692 |
| | | Psmb8 | Uqcrb | Ranbp1 | Gnao1 | Ywhae | Smim14 | Syne1 | Gm8730 |
| | | Ywhah | Rps27l | Cwc22 | Kcnj10 | Hint1 | Mrpl58 | Gsta4 | Elf1 |
| | | Ifngr1 | Rpl31.ps8 | Ppib | Qk | Gm12346 | Acot13 | Synm | Rpl36a.ps2 |
| | | Srgap2 | Atp5h | Cct6a | Clasp2 | Hnrnpdl | Aldoa | Eea1 | Mbnl1 |
| | | Mtdh | Srp14 | Set | Myo6 | Gm12696 | Slc25a39 | Mlc1 | Rpl14 |
| | | Gm9843 | Tceb2 | 1810037I17Rik | Hsp90aa1 | Myh10 | Ghitm | Tmem47 | Arhgdib |
| | | Mcl1 | Akr1a1 | Atp5o | Cldnd1 | Rack1 | Ndufb6 | Mycbp | Rps8 |
| | | Ly6e | Cox7a2 | Fkbp3 | Hspa8 | Usp1 | Hprt | Fam213a | Gm15421 |
| | | Hsp90b1 | Rock2 | Tceb2 | S100a16 | Cbx5 | Phyh | Odf2l | Gm43712 |
| | | Gm8995 | Ptms | Rsl1d1 | Atp8a1 | Hn1 | Ubl5 | Irak1bp1 | Pnrc1 |
| | | Capza2 | Chmp2a | Dnm1l | Fth1 | Myef2 | Suclg1 | Dync2h1 | Ccnd2 |
| | | Mbnl1 | Minos1 | Snrpd3 | Tmcc3 | Rheb | Tnfaip8 | Enah | Pfn1 |

| | | | | | | | | | |
|---|---|---|---|---|---|---|---|---|---|
| | | Rpl32 | Txn1 | Cox7a2 | Glul | Rps5 | Atp6v1e1 | Gm17018 | Gm15148 |
| | | Snx5 | Dstn | Gsk3b | Atp1b3 | S100a6 | Sh3bgrl | Tex9 | 2410006H16Rik |
| | | Rps9 | Itsn2 | Psma4 | Phyhipl | Hnrnph3 | Ndufs5 | Gas6 | Zfas1 |
| | | Cyfip1 | Ndufc1 | Akap9 | Fam168a | Atp5e | Vcam1 | Nudc | Actb |
| | | Tpm3 | Nap1l4 | Eprs | Adipor2 | Matr3 | Uqcrh.ps2 | Cfap36 | Gm10073 |
| | | Nfe2l2 | Atp5j2 | Cetn3 | Dock10 | Tcf12 | Tceb2 | Chchd6 | Mir703 |
| | | Akap13 | | Ndufb11 | Vmp1 | Gm3531 | Sdhc | Ezr | Ncl |
| | | Calr | | Mif | Eid1 | Supt16 | Ndufa12 | Pnmal2 | Eef2 |
| | | Clta | | Senp6 | Daam1 | Parp1 | Atp5j2 | Tuba1a | Cd47 |
| | | Fos | | Tm9sf3 | Gng11 | Srsf4 | Pla2g16 | BC005624 | Tmsb4x |
| | | Arpc2 | | Cfdp1 | Limch1 | Tubb2a | Ndufa8 | Tctex1d2 | Rps25.ps1 |
| | | Nfkbia | | Sfr1 | Hepacam | Gm11889 | Atp6v1d | Phtf1 | Gltscr2 |
| | | Rpl39 | | | Clic4 | Rbbp7 | Dnase2a | Sox9 | Serbp1 |
| | | Clic1 | | | Cfl2 | Ckap5 | RP23.269H21.1 | Ramp1 | Rpl22l1 |
| | | Klf6 | | | Selenok | Trim24 | Ndufb11 | Dnajb2 | Akap13 |
| | | Srgn | | | Reep5 | Smc1a | Bola3 | Plekhb1 | Gm12254 |
| | | Sat1 | | | Fkbp1a | Nop58 | Park7 | Nudc.ps1 | Rpl36al |
| | | Anxa3 | | | Zeb2 | Cct6a | Atp5b | Trim37 | Myh9 |
| | | Celf2 | | | Ptpn11 | Lgals1 | Romo1 | Cfap20 | Sub1 |
| | | Picalm | | | Zdhhc20 | Eny2 | Cox7a2 | Etfa | Rpl22.ps1 |
| | | Rps26 | | | Slu7 | Psma7 | Ndufab1 | Srek1ip1 | Cdkn1b |
| | | Gatm | | | Mid1ip1 | Ola1 | Uqcrh | Bola1 | Eif3h |
| | | Rpl26 | | | Gm12346 | Gm9769 | Tomm7 | Gstm1 | Rpl37a |
| | | Hspa5 | | | Ckb | Rpl30 | Uqcrb | Lztfl1 | Ablim1 |
| | | Btg1 | | | Hist1h2bc | Bud31 | Sod1 | S100b | Actr3 |
| | | Ier2 | | | Slc38a2 | Anp32b | Timm17a | Mt2 | Trim30a |
| | | Lamp2 | | | Pik3r1 | Hnrnpm | Nedd4 | Kcnq1ot1 | Tpm3.rs7 |
| | | Lamp1 | | | Dst | Ilf2 | Uqcrq | Hirip3 | Rapgef6 |
| | | Ssr4 | | | Nudt4 | Calm3 | Ggh | Pltp | Gm6863 |
| | | Gdi2 | | | Rnf130 | Gm4366 | Cacybp | Cluap1 | Rpl13 |
| | | Jund | | | Hsp90ab1 | Wdr89 | Ndufb7 | Naaa | Cox17 |
| | | Tln1 | | | Rap1a | RP24.389J11.1 | Gm5436 | Spats2l | Psme2b |
| | | Wasf2 | | | Lamp1 | Pdcd5.ps | Emc2 | Mpdz | Gm10076 |
| | | Sh3bgrl3 | | | Ppp2r2a | Vbp1 | Srek1ip1 | Arl13b | Gm14586 |
| | | Actr2 | | | Lap3 | Gadd45gip1 | Mrps21 | Ntrk2 | Iqgap1 |
| | | Ubc | | | Gm6977 | Srsf7 | Wbp5 | Gm5904 | Rpl31.ps8 |
| | | Tpm3.rs7 | | | Serinc1 | Gm6368 | Churc1 | Trp53bp2 | Rpl6 |
| | | Ccnl1 | | | Sgk1 | Rac1 | RP23.123D6.12 | Cald1 | Eif3e |
| | | Rplp2 | | | Wnk1 | Gm8566 | Grsf1 | Meg3 | Rgs1 |
| | | Ifnar2 | | | Dync1li2 | Sox2 | Dbi | Slf1 | Arpc1b |
| | | Rplp0 | | | Cd47 | Hdgf | Atp5f1 | Mt3 | Arpc5 |
| | | Rpl37 | | | Sypl | Gm10320 | Cox5b | Dnajc3 | Naca |
| | | Gm10275 | | | Fis1 | Gm5805 | Cfap36 | Mettl7a1 | Rpl18 |
| | | Gm5905 | | | Atxn7l3b | Polr2f | Atox1 | Ift20 | Psme1 |
| | | Lrp1 | | | Zfp706 | Dpy30 | Smdt1 | Pdia4 | Rps4x |
| | | Cd63.ps | | | Tmem30a | Syncrip | Arl4a | Dmd | Rpl36 |
| | | Gm10288 | | | Psip1 | Kpnb1 | Pdcd6 | Psph | Gm7027 |
| | | Cd63 | | | Pafah1b1 | Map1b | Oxct1 | Arl3 | Arpc2 |
| | | Msn | | | Slc25a4 | Serbp1 | Gm6265 | Cspp1 | Lrrfip1 |
| | | Ppt1 | | | Ifi27 | Rps16.ps2 | Atp6ap2 | Mt1 | Eif3f |
| | | Gm16580 | | | Hsbp1 | Gm8292 | Ndufa3 | Bex3 | Neat1 |
| | | Cstb | | | Pea15a | Gm6542 | Atp1a1 | Anxa5 | Sfpq |
| | | Rps19 | | | Dynll2 | Txn.ps1 | Timp2 | Itm2c | Tpm3 |
| | | Gm4149 | | | Rtn3 | Khdrbs1 | Ndufa6 | Mgst1 | Skil |
| | | Rpl41 | | | Mat2a | Whsc1 | Fam213a | Usp2 | Cd53 |
| | | Gm10073 | | | Tmem258 | Baz1b | Cox6b1 | Kif21a | Gm14681 |
| | | Rps20 | | | Specc1 | Fkbp3 | Hibadh | Cep162 | Gm8276 |
| | | Pfn1 | | | Reep3 | Hnrnpu | Mrpl42 | Cntrl | Hsp90ab1 |
| | | Rps10 | | | 1810037I17Rik | Gm4617 | Gde1 | Ahi1 | Sept11 |
| | | Coro1b | | | Sqstm1 | Calu | Calm3 | Gm2830 | Ddx5 |
| | | Rpl37rt | | | Ktn1 | Psmd14 | Ndufs6 | Anxa2 | Eef1g |
| | | Rpl6 | | | Tecr | Eif3k | Atp5a1 | Tusc3 | Rps6.ps4 |
| | | Rps25.ps1 | | | Tmbim6 | Hpf1 | Atpif1 | Hmgn5 | Fkbp3 |
| | | Arpc5 | | | Ociad1 | Srsf10 | BC002163 | Hist1h2bc | Anp32b |
| | | Ptp4a2 | | | S100a1 | Skp1a | Cyc1 | Hmgn1 | H2afy |
| | | Psme1 | | | Rab2a | 2310036O22Rik | Atp5e | Lmo4 | Eif3a |
| | | Rpl10a.ps1 | | | Nap1l4 | Brd8 | Acadm | Fam204a | Rps7 |
| | | Capzb | | | Hsd17b12 | Sept7 | Oaz1 | Zbtb20 | Eif5a |
| | | Psme2b | | | Ptn | Tuba1a | Syap1 | Tceal3 | Capzb |
| | | Rps13.ps2 | | | | Cops6 | Nudc | Aig1 | Ddx21 |
| | | Rps24.ps3 | | | | Cct5 | Slc25a5 | Cnn3 | Gm12183 |
| | | Rps15a | | | | Suz12 | Pkm | 2810013P06Rik | Rgs2 |
| | | Akr1a1 | | | | Taf15 | Scp2 | Stox2 | Sf3b1 |
| | | Sppl2a | | | | Dhx9 | Ndufv2 | Fundc1 | Ptbp3 |
| | | Rpl35a | | | | Sept11 | Tuba1b | Ptges3 | Kmt2a |
| | | Rps8 | | | | Naa38 | Hsp90aa1 | Timp2 | Prrc2c |
| | | Gm9794 | | | | Cct3 | Cox14 | Cntln | Tln1 |
| | | Cltc | | | | Nap1l1 | Zfp106 | Nap1l5 | Klf6 |
| | | Oxct1 | | | | Nfib | Srp14 | Crip2 | Eef1d |
| | | Rps3a1 | | | | Csnk2b | Cox7c | Gm1673 | Mier1 |
| | | Gm15427 | | | | Snrpd3 | Myeov2 | Prdx6 | Jund |
| | | Atpif1 | | | | Cct2 | Acat1 | Fam184a | Clk1 |
| | | Rps15 | | | | Ncl | Nenf | Fxyd1 | BC005537 |
| | | Grb2 | | | | Mrps24 | Sod2 | Wls | Clic1 |
| | | Rpl13 | | | | Sub1 | Ndufb5 | Gm4617 | Pfdn5 |
| | | Rpl37a | | | | Kif2a | Ndufa5 | Gm9800 | Srrm2 |
| | | Zfand5 | | | | Psmd8 | Eno1 | Cpd | Tpr |
| | | Rps10.ps1 | | | | Zfr | Ndufa2 | Atp1a1 | Myl12b |
| | | Os9 | | | | Gm8276 | Mpc2 | Cacybp | Txn1 |
| | | Arhgap5 | | | | Srsf2 | Cdv3 | Ccng1 | Cotl1 |
| | | Prpf38b | | | | H2afy | Trim37 | Clu | Rsbn1l |
| | | Tmem50a | | | | Rps6.ps4 | Slc25a3 | Atpif1 | Itgb1 |
| | | Mycbp2 | | | | Gm7809 | Gm12350 | Glod4 | Msn |
| | | Rpl36a.ps2 | | | | Gm14681 | Akr1a1 | Prdx4 | Btg2 |
| | | Wnk1 | | | | Tcea1.ps1 | Uqcrc2 | Pink1 | Rock1 |
| | | Rpl31.ps8 | | | | Fabp7 | Mdh2 | Cystm1 | Rsrp1 |
| | | | | | | Smc3 | Gm15920 | Pebp1 | Oaz1 |
| | | | | | | Pdap1 | Lrpap1 | Acadm | Itsn2 |
| | | | | | | Rpl8 | Swi5 | Cpe | Xrn2 |
| | | | | | | Gm5835 | Cox6a1 | Cetn3 | Tra2a |
| | | | | | | Sf3b6 | Ndufa7 | Oxct1 | Dek |
| | | | | | | Gm5963 | Slc25a4 | Spop | Hnrnpd |
| | | | | | | Gm6863 | Cep290 | Creld2 | Atp5d |
| | | | | | | Papola | Acadl | Gm12350 | Ywhaz |
| | | | | | | Rps7 | Hspa4 | Txn1 | Myl12a |
| | | | | | | Rpl14.ps1 | Dynll2 | Paip2 | Mycbp2 |
| | | | | | | Rps4x | Uqcrc1 | Csrp2 | Syf2 |
| | | | | | | Dynll1 | Atp6v1f | Akap9 | Gm12338 |
| | | | | | | Gm9385 | Chchd2 | Cfdp1 | Hnrnpab |
| | | | | | | Nudc | Pdha1 | Smim11 | Psma2 |
| | | | | | | Btf3 | Atp5k | Clcn4 | Thrap3 |
| | | | | | | Rps20 | Atp6v1g1 | Gng5 | Kmt2e |
| | | | | | | Atp5g3 | Atp5c1 | Stip1 | Ptma |
| | | | | | | Eif1ax | Nedd8 | Phpt1 | |
| | | | | | | Ppp1cb | Anp32e | Tomm7 | |
| | | | | | | Ran | Tmem256 | Pla2g16 | |
| | | | | | | Gm13436 | Rpl41 | Chchd2 | |
| | | | | | | Pou3f3 | Gm11966 | Hsp90ab1 | |
| | | | | | | Snx3 | Usp25 | Calm1 | |
| | | | | | | Hp1bp3 | Acbd5 | Bag1 | |
| | | | | | | Rtf1 | Mif | Nol7 | |
| | | | | | | Xrn2 | Lamp2 | Rnh1 | |
| | | | | | | Romo1 | Glrx5 | Laptm4a | |
| | | | | | | Gm13835 | Tmem59 | Cxcl13 | |
| | | | | | | Gm6563 | Paics | Shfm1 | |
| | | | | | | Ddx1 | Gnas | Gm42418 | |
| | | | | | | Ndufb6 | Gm12338 | | |
| | | | | | | Ptprz1 | mt.Nd5 | | |
| | | | | | | Zeb1 | Eif4g2 | | |
| | | | | | | Fez1 | mt.Nd2 | | |
| | | | | | | Bclaf1 | Tax1bp1 | | |
| | | | | | | Gm16580 | | | |
| | | | | | | Cbx1 | | | |
| | | | | | | Tuba1b | | | |
| | | | | | | Cyp51 | | | |
| | | | | | | Pcbp1 | | | |
| | | | | | | Spcs2.ps | | | |
| | | | | | | Txn1 | | | |
| | | | | | | Rps11 | | | |
| | | | | | | Rpsa.ps10 | | | |
| | | | | | | Hnrnph1 | | | |
| | | | | | | Lmo4 | | | |
| | | | | | | Zcrb1 | | | |
| | | | | | | Bzw1 | | | |
| | | | | | | Gm9892 | | | |
| | | | | | | Tmem258 | | | |
| | | | | | | Arl6ip1 | | | |
| | | | | | | Cct4 | | | |
| | | | | | | Cct8 | | | |
| | | | | | | Yy1 | | | |
| | | | | | | Eif2s2 | | | |
| | | | | | | Sox4 | | | |
| | | | | | | Srpk2 | | | |
| | | | | | | Set | | | |
| | | | | | | Eif3i | | | |
| | | | | | | Gspt1 | | | |
| | | | | | | Tceb1 | | | |
| | | | | | | Calm1 | | | |
| | | | | | | Rps3 | | | |
| | | | | | | Zfp207 | | | |
| | | | | | | Pcm1 | | | |
| | | | | | | Hnrnpc | | | |
| | | | | | | Txnl1 | | | |
| | | | | | | Pcbp2 | | | |
| | | | | | | Tmsb4x | | | |
| | | | | | | Psmd7 | | | |
| | | | | | | Suclg1 | | | |
| | | | | | | Nap1l4 | | | |
| | | | | | | Cox5a | | | |
| | | | | | | Gm6204 | | | |
| | | | | | | Srp72 | | | |
| | | | | | | Stag2 | | | |
| | | | | | | Pdia6 | | | |
| | | | | | | Dnajc2 | | | |
| | | | | | | Gm43712 | | | |
| | | | | | | Tmsb10 | | | |
| | | | | | | Rpl14 | | | |
| | | | | | | Nol7 | | | |
| | | | | | | U2surp | | | |
| | | | | | | Psmc1 | | | |
| | | | | | | Map2 | | | |
| | | | | | | Bola2 | | | |
| | | | | | | Banf1 | | | |
| | | | | | | H3f3a | | | |
| | | | | | | Rps27l | | | |
| | | | | | | Hspd1 | | | |
| | | | | | | Minos1 | | | |
| | | | | | | Nme1 | | | |
| | | | | | | Cacybp | | | |
| | | | | | | Ptma | | | |
| | | | | | | Npm1 | | | |

**Table S4.** Lists of differentially expressed genes between PDOXs and nude brain per each major cell type. Differentially expressed genes were defined at threshold: FDR <=0.01 and |log$_2$FC|>=1
Genes are listed from the highest to lowest |log2FC|

| **Cell type** | **Astrocytes** | | **Endothelial cells** | | **Ependymal cells** | | **Myeloid cells** | | **Oligodendrocytes** | | **OPCs** | | **Pericytes** | |
|---|---|---|---|---|---|---|---|---|---|---|---|---|---|---|
| Fold change | upregulated | downregulated | upregulated | downregulated | upregulated | downregulated | upregulated | downregulated | upregulated | downregulated | upregulated | downregulated | upregulated | downregulated |
| **Number of genes** | 186 | 129 | 304 | 105 | 21 | 22 | 287 | 136 | 7 | 124 | 286 | 199 | 2 | 71 |
| | Gm42418 | Sfrs18 | Gm42418 | Plp1 | Tmsb4x | Gm26924 | Spp1 | Sfrs18 | Olig1 | Igfbp5 | Pdgfra | Plp1 | Gm42418 | Wdr96 |
| | Cryab | Ppap2b | Rpl14.ps1 | Gm26924 | Gm42418 | Sez6 | Gm42418 | Snord49b | Pdgfra | Mal | Gm42418 | Apod | Gm12346 | Gm26924 |
| | Vim | Gm26924 | Adgrf5 | Gpr116 | Lyz2 | Vps8 | Cxcl13 | Sepp1 | Gria2 | Plp1 | Lhfpl3 | Npy | | 1700009P17Rik |
| | Tmsb4x | Gm26339 | Selenop | Sfrs18 | Tyrobp | Rpl35 | Clec7a | Gm26924 | Cspg5 | Phldb2 | Pnisr | Sepp1 | | 1700026D08Rik |
| | Pnisr | Snord42a | Gm6977 | Ttyh1 | Hexb | Vimp | Gm6977 | 4632428N05Rik | Cacng4 | Cd200 | Gap43 | Igfbp5 | | Ptplad1 |
| | Plpp3 | Xist | Gm5905 | Sepp1 | C1qa | D3Bwg0562e | Gm5905 | Epb4.1l2 | mt.Cytb | 4922501C03Rik | Cspg5 | Slc4a4 | | Stxbp3a |
| | Rack1 | Gm3764 | Gm9794 | Fabp7 | C1qb | Gm6625 | Gm14303 | Gm26339 | Actb | Slc4a4 | 3110035E14Rik | Clca1 | | A330021E22Rik |
| | Gm6977 | 2810055G20Rik | Rpl37rt | Igfbp5 | B2m | Calml4 | Gm9794 | Entpd1 | | Ptn | Rpl14.ps1 | Gja1 | | Wdr52 |
| | Gm9625 | Slc38a3 | Gm11478 | mt.Tc | Ctss | BC023829 | Rpl14.ps1 | Abca9 | | Sash1 | Gm5905 | Ptn | | Ccdc135 |
| | Rpl13.ps5 | D4Wsu53e | Gm4617 | Fut9 | C1qc | 1110001A16Rik | Ccl4 | D4Wsu53e | | Clca1 | S100a1 | Nr2f2 | | Ccdc19 |
| | Rps3a3 | Sepp1 | Gm15427 | D4Wsu53e | Ctsd | 0610011F06Rik | Rpl37rt | P2ry12 | | Mxi1 | Pllp | Ccdc80 | | 2610015P09Rik |
| | Gm14892 | Snord49b | Gm9385 | Ppap2b | Cd63.ps | Ppia | Lpl | Gpr34 | | Ccdc157 | Slc25a5 | Vtn | | Ccdc104 |
| | Gm7558 | Ptplad1 | Rps16.ps2 | Slc4a4 | Ctsb | Ccdc41 | Ttr | Il7r | | Tlr3 | Gng3 | Gm26924 | | Spag6 |
| | Fth.ps2 | Vegfa | Gm14303 | Eltd1 | Rpl14.ps1 | 1110059M19Rik | H2.Ab1 | Ddx26b | | Sepp1 | Scg5 | Frzb | | Rps17 |
| | Rpl14.ps1 | Sc4mol | Gm5805 | Dcn | Gm9794 | Rpl29 | Gm15427 | P2ry13 | | Ttyh1 | Ramp1 | Ttyh1 | | Sfrs18 |
| | Rps5 | Slco1c1 | Rpl32 | Gm26339 | Wdr89 | 4922501C03Rik | Fth1 | Csmd3 | | Lifr | Rps5 | Apoe | | Ifltd1 |
| | Gm12346 | Adrbk2 | Gm4149 | Rps6 | Rack1 | Gm9493 | Gm11478 | Gm17087 | | Frzb | Gm9385 | Hmgcs2 | | 2510003E04Rik |
| | Gm12857 | 2610017I09Rik | Gm10080 | Snord49b | Lamp1 | Cox7b | Cd63.ps | Ccdc55 | | Wdr60 | Olig2 | Timp3 | | Gm872 |
| | Gm13215 | Slc1a2 | Gm5963 | Hnrnpa3 | Sparc | Stk39 | H2.Aa | Maf | | Timp3 | Rpl3.ps1 | Ncam1 | | Ppap2b |
| | Gm6136 | Vimp | Pnisr | Ptn | Cst3 | Selt | Rack1 | Gpr56 | | Trp53bp2 | Gm12346 | Inhba | | 2010015L04Rik |
| | Rps16.ps2 | Zfml | Rpl3.ps1 | Slc7a11 | Actb | Enpp2 | Cd74 | Hnrnpa3 | | Mybpc1 | Rpl37rt | Sash1 | | A030009H04Rik |
| | Gm13498 | Tmem66 | Adgrl4 | Vimp | | 4930546H06Rik | Ccl3 | Tmem66 | | Ralgps2 | Gm9794 | Iqub | | Pax3 |
| | Gm22567 | Syne1 | Rpl38.ps2 | Pcdhga9 | | | Gm16580 | Ppap2b | | Sfrs18 | Gm11478 | Col5a2 | | Vps28 |
| | Gm12183 | St6galnac5 | Tpt1.ps3 | Atp1a2 | | | Gm11942 | Zfhx3 | | Aspa | Tnr | Tspan15 | | D2Wsu81e |
| | Gm9385 | CT030684.1 | Gm9843 | Abca9 | | | Gm5963 | Jund | | Slc7a11 | Gm11560 | Stmn1 | | Gm6625 |
| | Gm5805 | Nim1 | Rack1 | Hes5 | | | Gm4149 | CT030684.1 | | Gja1 | Olig1 | Lpar1 | | Ccdc41 |
| | Rn7s6 | Fut9 | Rps24.ps3 | Slc1a3 | | | Rps24.ps3 | Zfml | | Plekha4 | Clu | Igf1 | | Selk |
| | Id1 | Cwc22 | Gm14586 | Cyp2j9 | | | Selenop | Nrip1 | | Rgag4 | Rack1 | Wls | | Lsmd1 |
| | Fth1 | Gnao1 | Gm6204 | Timp3 | | | Cst7 | Slco2b1 | | Gas7 | Gm15427 | Sparc | | 6720401G13Rik |
| | Gm12222 | Pcdhga9 | Gm11966 | Fam213a | | | Rps16.ps2 | Tmem119 | | Ahctf1 | Rsrp1 | Col23a1 | | B630005N14Rik |
| | Gm5370 | 2610203C20Rik | Rflnb | mt.Tp | | | Gm9385 | Vimp | | Inhba | Bcas1 | Slc7a11 | | AK129341 |
| | Gm11560 | Trim9 | Gm11560 | Nell2 | | | Gm5805 | Hpgds | | Fmo1 | Rps16.ps2 | Fam213a | | Gm26339 |
| | Gm12165 | S1pr1 | Rps13.ps2 | Scd2 | | | Rpl3.ps1 | Lrrc58 | | Gtpbp2 | Gm4617 | Sfrs18 | | Sept15 |
| | Gm13340 | Macf1 | Rpl36a.ps2 | Sash1 | | | Rps13.ps2 | Cx3cr1 | | Fbln2 | Tmem100 | Gm26339 | | Ankrd32 |
| | Rpl10l | 5830428H23Rik | Gm11942 | Ptgds | | | Pnisr | Il6ra | | Gm872 | Stmn3 | Hey2 | | Dpcd |
| | Gm11942 | Ttyh1 | Fn1 | Vegfa | | | Apoe | Cep110 | | Cldn5 | Gm12338 | Phldb2 | | CT030684.1 |
| | Rpl17.ps4 | Ank2 | Gm16580 | Gm17087 | | | Fn1 | Gm13476 | | Mdk | Calm1 | Cd59a | | Gm12942 |
| | Gm4332 | 4930546H06Rik | Wdr89 | Mllt4 | | | Ccl6 | Mertk | | Cyp4v3 | Nap1l5 | Gdpd4 | | Gm5918 |
| | Gm11810 | Ctage5 | Gm10073 | 2810055G20Rik | | | Gm6204 | Wapal | | Pla2g16 | Bex3 | Fmo1 | | Pet112 |
| | Rps15.ps2 | Dclk1 | Gm4332 | Slc1a2 | | | Gm8995 | Naalad2 | | Scd1 | Id2 | Pla2g16 | | Sc4mol |
| | Gm14450 | Malat1 | Rsrp1 | Colec12 | | | Rpl36a.ps2 | RP24.312B12.1 | | Wls | Pcdh15 | Mfap3l | | 2410018M08Rik |
| | Gm12696 | Clmn | Rps26 | Wapal | | | Cd52 | Rps23 | | Pbxip1 | Rpsa.ps10 | Atp1a2 | | I7Rn6 |
| | Gm6368 | Acss1 | Rplp0 | Mdk | | | H2.K1 | Snord42a | | Fam213a | Ranbp1 | Mybpc1 | | BC023829 |
| | Wdr89 | Itih3 | Slc38a5 | Gm14964 | | | Kctd12 | Rpl5 | | Igf1 | C1ql1 | Mal | | Ccdc37 |
| | Gm2225 | Vwa1 | Gm9800 | Mboat2 | | | Epb41l2 | Ppp1r9a | | Dab2 | Cd81 | Col27a1 | | Epb4.1l4a |
| | Gm5963 | Pitpnc1 | Rplp1 | Rpl15 | | | Rpl10a.ps1 | Smap2 | | Col5a2 | Ostf1 | Dnajc1 | | 9330101J02Rik |
| | Ttr | Fam21 | Gm6863 | Suv420h1 | | | Rps5 | Jhdm1d | | Slc43a2 | Gria3 | Pex11a | | Epb4.1l2 |
| | Gm11472 | Cep110 | Rpl31.ps8 | Fbln2 | | | Ch25h | Rps29 | | Lysmd3 | Igfbp2 | Abca8a | | 4922501C03Rik |
| | Rps12.ps19 | 4930402H24Rik | Gm12338 | Rpl13a | | | Gm4617 | Rpl21 | | Fstl1 | Cox8a | Dock7 | | 2700089E24Rik |
| | Gm17909 | Appl2 | Rpsa.ps10 | Atp13a5 | | | Gm14586 | 2810055G20Rik | | Frg1 | Dynll1 | Fbln2 | | 1700094D03Rik |
| | Rac1 | Tril | Rps10.ps1 | Rpl5 | | | Gm4332 | Tanc2 | | Hey2 | Nxph1 | Rnd3 | | Bai3 |
| | Gm12152 | Epb4.1l2 | Gm10288 | Sept15 | | | Rsrp1 | Son | | Ncam1 | Selenok | Filip1l | | Sepn1 |
| | Cd63.ps | Mycbp2 | Gm6265 | 4933426M11Rik | | | Rpsa.ps10 | Slc16a6 | | Atp8a1 | Alcam | Golgb1 | | Lphn3 |
| | Gm5835 | Atp1a2 | Scgb3a1 | Gja1 | | | Ctsd | D15Ertd621e | | Acss1 | Gm4149 | Thrsp | | 2310044G17Rik |
| | Gm9843 | Zfp949 | Fth1 | Gpm6b | | | Lilrb4a | RP24.288C12.6 | | Cd82 | Actb | 2610203C20Rik | | Rpl24 |
| | Cald1 | Acsl6 | Ctla2a | Kcnj16 | | | Ifi27l2a | Arid4a | | Glul | Rps26 | Gas7 | | Nim1 |
| | Id3 | Gja1 | Rps5 | Kmt2a | | | Rps26 | A430104N18Rik | | Mfsd6 | Gm14586 | Glul | | 2410066E13Rik |
| | Hspd1.ps3 | Timp3 | Gm10275 | Tmem66 | | | Rps25.ps1 | Chd9 | | Selk | Gm6863 | Sdc4 | | Gm10311 |
| | Gm2178 | Son | Apold1 | Spag9 | | | Tpt1.ps3 | Numb | | Lgi1 | Tpt1.ps3 | Cry2 | | Gm13225 |
| | Gm9794 | Ccdc104 | Gm12183 | Ptms | | | Mt1 | Srgap2 | | Gramd1b | Emid1 | Scd1 | | 5430417L22Rik |
| | Map1b | Btbd17 | Gm7809 | Gprasp1 | | | Gm6030 | Elmo1 | | Npy | Ndufa4 | Hsd17b11 | | Gm17388 |
| | Gm44010 | Ddhd1 | Afdn | Morf4l1 | | | Calr.ps | Pwwp2a | | Sco1 | Igfbp3 | Vimp | | Gpr125 |
| | Rps19.ps6 | Suv420h1 | Gm15500 | Rpl10 | | | Lyz2 | Ctage5 | | Slc25a13 | Car8 | Mllt4 | | Grlf1 |
| | Gm5778 | Ppia | Gm5835 | Zbtb20 | | | Gm10288 | Col27a1 | | Adam23 | B2m | Gucy1b3 | | Leprel2 |
| | Gm4204 | Slc1a3 | Rps25.ps1 | Hspa8 | | | Wdr89 | Ogfrl1 | | Tgfbr1 | Rps13.ps2 | Fam3c | | Trp53rk |
| | Selenop | Ttc14 | Gm9892 | Htra1 | | | Gm10073 | Mycbp2 | | Fgd4 | Gm14303 | Gpr56 | | Zbed6 |
| | Gm15159 | Slc7a10 | Calr.ps | S1pr3 | | | H2.D1 | Scamp2 | | Tmem66 | Gm6977 | Abca5 | | Fam101a |
| | RP23.390K13.4 | Nrcam | Rpl18.ps1 | Rpl23a | | | Gm15500 | Lpar6 | | Nr2f2 | Pcsk1n | Edil3 | | Fam115a |
| | Gm11478 | Pdcd4 | Rpl10a.ps1 | Kcnj10 | | | Rpl31.ps8 | Marcks | | BC016423 | 2900011O08Rik | Huwe1 | | 9830147E19Rik |
| | Cxxc5 | Pla2g7 | Gm8894 | Ccdc80 | | | Rpl18.ps1 | Erbb2ip | | Zmat1 | Gm5805 | Ift57 | | Rom1 |
| | Bex3 | Fam115a | Gm8292 | Rps23 | | | Apoc1 | Rbm5 | | Gm26339 | Ndufb6 | Aldh2 | | Zfp191 |
| | Gm16580 | Gyk | Rps6.ps4 | Ncam2 | | | Gm5835 | Tmem100 | | Aim1 | Cd63.ps | Ahcyl2 | | |
| | Gm15896 | Csgalnact1 | Selenom | Gnb2l1 | | | AU020206 | Rps6 | | Dock7 | Ndufa8 | Efnb2 | | |
| | Gm27046 | Nrxn1 | Ly6c1 | Inhba | | | Gm9843 | Macf1 | | Tiam1 | Rpl22l1 | Grik3 | | |
| | Gm29228 | Hlf | Eif1.ps1 | Cwc22 | | | Gm8276 | Ythdc1 | | 4931406C07Rik | Banf1 | Mgst1 | | |
| | Cnot2 | Rorb | Gm12254 | Prrc2c | | | Rplp1 | Ssh2 | | Vtn | Ddah1 | Hmgn2 | | |
| | Gm9294 | D14Abb1e | Cd34 | Ncam1 | | | Tyrobp | Qk | | Osbpl11 | Ttr | Gm12688 | | |
| | Gm11517 | Flrt1 | Rps15a.ps5 | Zfml | | | Il1b | Hmgb1 | | Phf17 | Gpm6a | Sparcl1 | | |
| | Rpl34.ps2 | Frmd4a | Rps11 | Gpx4 | | | Ccrl2 | Med12l | | Hspa8 | Scrg1 | Fut9 | | |
| | Gm9521 | Fjx1 | Ttr | S100b | | | Kcnq1ot1 | Arl4c | | Cpt1a | Wdr89 | Mest | | |
| | Gm8513 | Acin1 | Tmsb4x | Rps17 | | | AC149090.1 | Cwc22 | | 5830418K08Rik | Gm12222 | Map4k4 | | |
| | Gm4321 | Arhgap5 | Gm3531 | Xist | | | Rpl32 | Dusp6 | | Tpr | S100a16 | Ptgds | | |
| | Gm15427 | Hnrnpa3 | B2m | 4932438A13Rik | | | Rps10.ps1 | Sall1 | | Apoe | Gm12696 | Plcb4 | | |
| | Gm18014 | Mfge8 | Ly6e | Ahcyl2 | | | Axl | Dip2b | | Tmem245 | Tuba1a | Slc27a1 | | |
| | Gm12482 | C030037D09Rik | Rps9 | RP24.288C12.6 | | | Id2 | Rps27a | | Hadha | Cdo1 | Mex3a | | |
| | RP23.408J7.2 | Tnik | Rps26.ps1 | Ddit3 | | | Vim | Frmd4a | | Atp1a2 | Tceal3 | Grin3a | | |
| | Gm17511 | Selk | Selenok | Ppia | | | Ctsb | Rps28 | | Nnat | Gm9892 | Cyp2d22 | | |
| | Gm6543 | Ubn2 | Gm12346 | Rps27 | | | Itgax | P4ha1 | | Snx24 | Gm6204 | Rgag4 | | |
| | Matr3.ps2 | Slc4a4 | Cdkn1a | Mga | | | Vsir | Morf4l1 | | Dnajc1 | Gm5778 | Fabp7 | | |
| | Gm7308 | Tmed5 | Rpl8 | Ccdc104 | | | Rps20 | Olfml3 | | Plk1s1 | Nrxn1 | Cd200 | | |
| | Gm5514 | Cadm2 | Selenof | Rps12 | | | Cd300c2 | Hspa8 | | Clca2 | Plpp3 | Wapal | | |
| | Gm4366 | Nwd1 | Gm12350 | Kdm5b | | | Gm6863 | Prrc2c | | Ccdc39 | Tmem176b | Syne2 | | |
| | Rsrp1 | Dio2 | Gm15772 | Igfbp2 | | | Ccl12 | Fau | | Stmn1 | Gm5835 | Ccnd2 | | |
| | Id2 | P4ha1 | Igfbp7 | Diap2 | | | Gm11560 | Map4k4 | | Cd55 | Calr.ps | D3Bwg0562e | | |

| | | | | | | | | | | | | |
|---|---|---|---|---|---|---|---|---|---|---|---|---|
| | Rps3 | Acsl3 | Pomp | Rpl21 | | | Gm14681 | Safb | | 2700089E24Rik | Cox6b1 | Nid1 |
| | S100a6 | Tex9 | Gm6472 | Apod | | | Ints6l | Arhgap5 | | Mfap3l | Mmp15 | Suv420h1 |
| | Gm3531 | Myo6 | Sparc | Nupr1 | | | mt.Cytb | Acin1 | | P2ry12 | Dynlt1.ps1 | Hadha |
| | Gm7618 | Sparcl1 | Slc25a5 | Wdr60 | | | mt.Nd1 | Ifngr1 | | Fabp7 | Cfap20 | Tlr3 |
| | Gm5905 | Mfn1 | Gm7027 | Tmx4 | | | Cd83 | Bin2 | | Fut9 | Ccnd1 | Clca2 |
| | Gm12912 | Gm973 | Gm6368 | Son | | | Gm10275 | Srrm2 | | Fam3c | Gm5963 | Dtx4 |
| | Gm13815 | Slc6a11 | Gm14681 | Ednrb | | | Gm9892 | Tsc22d4 | | Trim41 | Cfap36 | Abcb1a |
| | Rps11 | Kcnj16 | Ppfibp1 | Stmn1 | | | Rps15a.ps5 | Selplg | | Mecp2 | Tagln2 | Gm6483 |
| | Gm11488 | Ythdc1 | Rab1a | Casc4 | | | Gnas | Zfp292 | | Rbms3 | Gm6265 | Gpd1 |
| | Pabpc1 | Mdn1 | Rpl22.ps1 | Asph | | | Rpl38.ps2 | Mtdh | | Scd2 | H2.D1 | Slc1a3 |
| | Dclk2 | Ntrk2 | Gm4366 | Nr2f2 | | | Flt1 | Ccr5 | | P2rx4 | Rpl36a.ps2 | Slc13a3 |
| | Gm12020 | Sycp2 | Rpl9.ps7 | | | | Gm7027 | Adap2 | | E130311K13Rik | Hspd1.ps3 | Gpd2 |
| | Gm15773 | Trpm3 | Gm8995 | | | | Selenos | Abcb4 | | Lrig1 | Grm5 | Zbtb20 |
| | Calm2 | Wapal | Gm15421 | | | | Ank | Vwa1 | | Rnd3 | Pcp4 | Itsn2 |
| | App | AI464131 | Hsp25.ps1 | | | | Gm12338 | Ivns1abp | | Paqr6 | Cacng4 | Chd9 |
| | Gm6204 | Mmp14 | Gm14539 | | | | Rps6.ps4 | Plcl2 | | Ltn1 | Meg3 | Sat1 |
| | Calr.ps | Pfkp | Plpp3 | | | | B2m | Pde3b | | Lama4 | Dnm3 | 4922501C03Rik |
| | Vcan | Neat1 | Gm6563 | | | | Srgn | Sox4 | | Nek9 | Rab31 | Hnrnpa3 |
| | Spp1 | Nebl | Anp32.ps | | | | Gm12346 | Fchsd2 | | Son | Slc38a3 | Ptms |
| | Marcks | Chrdl1 | Tpm3.rs7 | | | | Atf3 | Rps27 | | Slc16a1 | Pdcd5.ps | Tspan18 |
| | Fos | Dtna | Gm12696 | | | | Cd63 | Malat1 | | B4galt6 | Ubb | Arhgap5 |
| | Hacd3 | Aifm3 | Saraf | | | | Atox1 | Rock2 | | Zbtb20 | Gm10288 | Magt1 |
| | Gm6863 | Utp14b | Ctsd | | | | Ifi202b | Ttc14 | | Ccdc171 | Ppib | Son |
| | Gm21399 | Peg3 | Gm5436 | | | | Cstb | Tgfbr1 | | Nexn | Dpp6 | Gbp7 |
| | Atp5e | 9330159F19Rik | Gm5865 | | | | Nfkbia | Ubn2 | | Colgalt2 | Oxct1 | Nupr1 |
| | Ctsd | Sfxn5 | S100a11 | | | | Rps26.ps1 | March1 | | Sparcl1 | Serpine2 | Nfat5 |
| | Gm12892 | Slc7a11 | Cyb5a | | | | Ctsz | Pnn | | Calr | Arl2bp | 9330159F19Rik |
| | Golim4 | Prpf4b | RP23.123D6.12 | | | | Ms4a6c | Rbm25 | | 4930480K23Rik | Gm3531 | Cyp39a1 |
| | B2m | Slitrk2 | Cd63.ps | | | | Gm12254 | Rpl35 | | Kdm2a | Gm11966 | Igfbp4 |
| | Jun | Gprasp1 | Rpl39 | | | | Rps10.ps2 | Ppia | | | Rtn1 | Ccdc104 |
| | Selenos | Ankrd11 | Ybx1.ps2 | | | | Npc2 | Pmepa1 | | | Gng11 | Gm6625 |
| | Gfap | Cxcl14 | Shfm1 | | | | Atp5e | Serinc3 | | | Rpl10a.ps1 | Snx29 |
| | Gm11966 | Gapdh | AU021092 | | | | Ifi44 | Rpl10 | | | Arpc2 | Tec |
| | Sept7 | Kcnk1 | Gm8276 | | | | Csf1 | Nav3 | | | Rps26.ps1 | Cmtm5 |
| | F3 | Ccdc50 | Gm13835 | | | | Gm15772 | Gnb2l1 | | | Agpat5 | Mboat2 |
| | Hacd2 | | Gm5644 | | | | Tnf | Rpl18a | | | Ndufs4 | Kdm5b |
| | Gnas | | Gm8730 | | | | Mia2 | Rpl17 | | | Gm5865 | Abca1 |
| | Cct2 | | Smim10l1 | | | | Gm15421 | Ptgs1 | | | Phlda1 | RP24.288C12.6 |
| | Rps20 | | Gm5778 | | | | Rpl9.ps6 | Ywhah | | | Rps24.ps3 | M6pr |
| | Rpl32 | | Gimap4 | | | | Plpp3 | Hmha1 | | | Cox4i1 | Nfe2l2 |
| | Gatm | | Gm6733 | | | | Rgs1 | Basp1 | | | Tmem179 | Zfhx3 |
| | Rpl8 | | Vim | | | | Calm1 | Srrm1 | | | Ahi1 | Sort1 |
| | Sbds | | Pdcd5.ps | | | | Rpl39 | | | | Fis1 | Wnt6 |
| | Skp1a | | Cfap36 | | | | Rpl22.ps1 | | | | Selenof | Tmem132b |
| | Msmo1 | | Rps3 | | | | Gm11966 | | | | Eif3i | Frmd4a |
| | Snx3 | | Tspan13 | | | | Gm7266 | | | | Ly6h | Phf17 |
| | Ybx1 | | Rpl18.ps2 | | | | Gm12183 | | | | Rplp1 | Entpd1 |
| | H3f3b | | Nsrp1 | | | | Cox4i1 | | | | Ndufa7 | Agt |
| | Eif3k | | Rps19.ps6 | | | | Ctsl | | | | Gm16580 | Cldn5 |
| | Sox4 | | Wfdc1 | | | | Gm9800 | | | | Gm15500 | Crip2 |
| | Hint1 | | Ifitm3 | | | | Cox6a1 | | | | Gm4204 | Plcd4 |
| | Calm1 | | Selenow | | | | Rps15 | | | | Omg | Lix1 |
| | Psma7 | | mt.Cytb | | | | Dbi | | | | Ednrb | Ctbp2 |
| | Fez1 | | Tcea1.ps1 | | | | Rps11.ps1 | | | | Gria2 | Cyp2j6 |
| | Rheb | | Higd2a | | | | Cxcl2 | | | | Rpl36al | Baz2b |
| | Nucks1 | | Ier2 | | | | Saraf | | | | Tspan13 | Socs2 |
| | Eef2 | | H2.K1 | | | | Rps19.ps6 | | | | Gm6368 | Atp8a1 |
| | Cbr1 | | Gm7266 | | | | Cox6b1 | | | | Cox7b | Cyp4v3 |
| | H2.D1 | | Selenos | | | | Hacd2 | | | | Il1rap | Snord49b |
| | Tcf12 | | Gm12481 | | | | Wfdc17 | | | | Commd6 | Stx5a |
| | Eef1a1 | | Gm15920 | | | | mt.Nd4 | | | | Gm14681 | Cep85l |
| | Matr3 | | Cox7a2 | | | | Capg | | | | Mpc2 | Nell2 |
| | Rps9 | | Gpx1 | | | | Prdx1 | | | | Atp5g3 | Fxyd1 |
| | Anapc11 | | Rps10.ps2 | | | | Pld3 | | | | Ndufa1 | Parp4 |
| | Rps14 | | mt.Nd1 | | | | Lgals3bp | | | | Cct7 | Zfp191 |
| | Gtf2i | | Cox4i1 | | | | Postn | | | | Nsg1 | 2310022B05Rik |
| | Bcan | | Actb | | | | Gm6136 | | | | Psmb1 | Bmp7 |
| | Rps4x | | Igf1r | | | | Gm4366 | | | | Cfl1 | Hspa8 |
| | Ubb | | Ostf1 | | | | Gm8730 | | | | Prdx2 | Foxo1 |
| | Hnrnpd | | Mia2 | | | | Grn | | | | Dbi | Eps15 |
| | Dync1li2 | | Fkbp1a | | | | Slc25a5 | | | | Sulf2 | Cdh8 |
| | Rps21 | | Eif3j2 | | | | Rplp0 | | | | Dtd1 | Paqr6 |
| | Rps15 | | Rps15.ps2 | | | | Gm4604 | | | | Cspg4 | Slc43a3 |
| | Tpm3.rs7 | | Ube2d3 | | | | Fcer1g | | | | Pcyt1b | Chd2 |
| | Rpl4 | | Arpc2 | | | | Cd9 | | | | Dscam | Fnbp1l |
| | Rplp1 | | Gm4604 | | | | Zfp638 | | | | Mlf2 | Notch2 |
| | Dstn | | Tonsl | | | | Cox8a | | | | Gm1673 | Sipa1l1 |
| | Sec11c | | Ramp2 | | | | Rps3 | | | | Nop10 | Tmtc2 |
| | Cnn3 | | Pabpc1 | | | | Pabpc1 | | | | Uqcrq | Lipg |
| | Nasp | | Gm14253 | | | | Eif1.ps1 | | | | Atxn10 | Prrg3 |
| | Ppp1cb | | Hint1 | | | | Gusb | | | | Ckb | Scd2 |
| | Vapa | | Id3 | | | | Cntrl | | | | A730017C20Rik | Megf10 |
| | Ilf3 | | Gm9840 | | | | Gm6265 | | | | Serpina3n | Hivep3 |
| | Pcbp2 | | Slc16a4 | | | | Ier3 | | | | Aplp1 | Slc25a13 |
| | Cct6a | | Rps15 | | | | Lgals1 | | | | Higd2a | Ncoa7 |
| | Ywhab | | Rpl9.ps6 | | | | Tmsb10 | | | | Cxcl14 | Synm |
| | Pcbp1 | | Pglyrp1 | | | | Gpx1 | | | | Rps9 | Leprel2 |
| | Ctss | | Gstp.ps | | | | Csf2ra | | | | Dner | RP23.32A8.1 |
| | Zhx1 | | BC002163 | | | | Gm43712 | | | | Pcbp2 | Prss56 |
| | Ywhae | | Tpm4 | | | | Hpf1 | | | | Olfm1 | Fam107a |
| | Arpc2 | | Epb41l2 | | | | Gm6368 | | | | Cryab | Tiam1 |
| | Rpl23 | | Naa38 | | | | Rps15.ps2 | | | | Kcnd2 | Gm4076 |
| | | | Gm12020 | | | | Rps9 | | | | Znhit6 | CT030684.1 |
| | | | Ndufa11 | | | | Anxa5 | | | | Gtf2h5 | Tmem47 |
| | | | Calm1 | | | | E230029C05Rik | | | | Ndufc2 | Cpne8 |
| | | | S100a16 | | | | Gm5644 | | | | Opcml | Ccdc173 |
| | | | Gm6542 | | | | Uqcrq | | | | 1810037I17Rik | Erbb3 |
| | | | AC149090.1 | | | | Rpl18.ps2 | | | | Uchl3 | Vwa1 |
| | | | Ssr2 | | | | Gm6472 | | | | Cdh13 | Cgnl1 |
| | | | Rpl37 | | | | Wapl | | | | Sfr1 | Tnfrsf19 |

| | | | | | | | | | | | | |
|---|---|---|---|---|---|---|---|---|---|---|---|---|
| | | | Gm13436 | | | | Rps19 | | | | Gm2a | Itga6 |
| | | | Gm10039 | | | | Nsrp1 | | | | Cp | Zfp609 |
| | | | Grrp1 | | | | Mir703 | | | | Tubb2a | mt.Rnr1 |
| | | | S100a13 | | | | Sh3bgrl3 | | | | 3632451O06Rik | Alx4 |
| | | | Tpm3 | | | | Rpl9.ps7 | | | | Gjc3 | Prrc2c |
| | | | Gm7407 | | | | Gm10080 | | | | Atp6v1g1 | |
| | | | Fam204a | | | | Rps11 | | | | Lnx1 | |
| | | | Myl12a | | | | Cotl1 | | | | Adam9 | |
| | | | Gm10031 | | | | Gm8606 | | | | Pea15a | |
| | | | Txn.ps1 | | | | Lyzl4 | | | | Sfxn1 | |
| | | | Atp5j | | | | H2.DMa | | | | Pfn1 | |
| | | | Psma5 | | | | Plau | | | | Uqcrc1 | |
| | | | Ccdc186 | | | | Akr1a1 | | | | Srp14 | |
| | | | Gm6136 | | | | Gm12020 | | | | Cdk4 | |
| | | | Ifi27l2a | | | | Cybb | | | | Itgb8 | |
| | | | Rplp2 | | | | Psme2b | | | | Deb1 | |
| | | | Dynlt1.ps1 | | | | Ssr4 | | | | Megf11 | |
| | | | Sf3b6 | | | | Cd72 | | | | Cox6c | |
| | | | Zfp638 | | | | Selenof | | | | Myo5a | |
| | | | Rps11.ps1 | | | | Gm15148 | | | | Pou3f1 | |
| | | | Wapl | | | | Zfas1 | | | | Tacc2 | |
| | | | Icam2 | | | | Ndufa2 | | | | Atp5j | |
| | | | Rac1 | | | | Gm9294 | | | | Gnb4 | |
| | | | Gm14513 | | | | Shfm1 | | | | Cadm2 | |
| | | | Bag1 | | | | Il4i1 | | | | Atp6v1a | |
| | | | Gm43712 | | | | Gm13436 | | | | Gria4 | |
| | | | Gm21399 | | | | Gm12481 | | | | Sept7 | |
| | | | Tmem252 | | | | Erbin | | | | Brinp1 | |
| | | | Rps15a | | | | Ppp1r15a | | | | Eif5a | |
| | | | P4hb | | | | Selenok | | | | S100a13 | |
| | | | Lgals1 | | | | Rpl22l1 | | | | Ndufb8 | |
| | | | Tmsb10 | | | | Eif3f | | | | Pgp | |
| | | | Pcbp1 | | | | Ifitm3 | | | | Psmb3 | |
| | | | Uqcrh.ps2 | | | | Rps21 | | | | Bcan | |
| | | | Gm11964 | | | | Zeb2os | | | | Gm10250 | |
| | | | H2.D1 | | | | Gm1966 | | | | Ptprn | |
| | | | RP23.269H21.1 | | | | Gm14513 | | | | Krtcap2 | |
| | | | Ilk | | | | Ctsh | | | | Atp5b | |
| | | | Cox6b1 | | | | Mt2 | | | | Sri | |
| | | | Lmo2 | | | | Glmp | | | | Cask | |
| | | | Rasip1 | | | | Gm10076 | | | | Tceb2 | |
| | | | Psme2b | | | | Pdcd5.ps | | | | Mt3 | |
| | | | Erbin | | | | Pfdn5 | | | | Ppp1r14c | |
| | | | Zmat2 | | | | Cxcl16 | | | | Park7 | |
| | | | Gm9294 | | | | Crip1 | | | | C4b | |
| | | | Gata2 | | | | Gm6394 | | | | Sox6 | |
| | | | Oaz1.ps | | | | Cspg4 | | | | Pcdh9 | |
| | | | Anp32b.ps1 | | | | Ier2 | | | | Uqcrb | |
| | | | Atox1 | | | | Myeov2 | | | | Rgs20 | |
| | | | Rras | | | | Cd68 | | | | Hint1 | |
| | | | Rhoj | | | | Gm8692 | | | | Ugp2 | |
| | | | Ly6a | | | | RP24.389J11.1 | | | | Mdh2 | |
| | | | Mydgf | | | | Gm4204 | | | | 6330403K07Rik | |
| | | | Rps19 | | | | Gm3531 | | | | Ndufab1 | |
| | | | Uqcr11 | | | | Gm14539 | | | | Mt1 | |
| | | | Esam | | | | Lgals3 | | | | Atp6v0e | |
| | | | Rps20 | | | | Nfkbiz | | | | Rpl32 | |
| | | | Gm8129 | | | | Gm8292 | | | | Pomp | |
| | | | Gm8566 | | | | Gm11361 | | | | Rps3 | |
| | | | Psmb2 | | | | Snhg8 | | | | Rpl39 | |
| | | | Cldn5 | | | | Ifi207 | | | | Psma7 | |
| | | | Bvht | | | | Rps14 | | | | Isoc1 | |
| | | | Snhg8 | | | | Gm12696 | | | | Zcchc24 | |
| | | | Gm12231 | | | | Ybx1.ps2 | | | | Bzw1 | |
| | | | Junb | | | | 2010107E04Rik | | | | Cdc42 | |
| | | | Hpf1 | | | | Diaph2 | | | | Cox6a1 | |
| | | | Chchd2 | | | | Mydgf | | | | Lrrtm3 | |
| | | | Ndufa8 | | | | Bcl2a1b | | | | Pafah1b1 | |
| | | | Pea15a | | | | Mpeg1 | | | | Phyhipl | |
| | | | Alas1 | | | | Cox7a2 | | | | Ptpre | |
| | | | Anxa2 | | | | Rpl8 | | | | Ran | |
| | | | Hspd1.ps3 | | | | Spcs2.ps | | | | Gm10260 | |
| | | | Csnk2b | | | | Cox6c | | | | Cox7a2 | |
| | | | Suclg1 | | | | Gadd45b | | | | Wbp5 | |
| | | | Gm16399 | | | | Hspd1.ps3 | | | | Acat1 | |
| | | | Id1 | | | | Atp6v1f | | | | Cd9 | |
| | | | Ost4 | | | | Rab1a | | | | Rps14 | |
| | | | Eea1 | | | | Ccl9 | | | | Atpif1 | |
| | | | Gabarapl1 | | | | RP23.123D6.12 | | | | Atp5a1 | |
| | | | Dpm3 | | | | Arpc1b | | | | Tmem176a | |
| | | | Spcs2.ps | | | | Tspo | | | | mt.Cytb | |
| | | | Gng5 | | | | Txn1 | | | | Phactr3 | |
| | | | Txndc17 | | | | Rpl39.ps | | | | Atp5e | |
| | | | Sec61b | | | | Ndufa1 | | | | Sdhb | |
| | | | Gm6394 | | | | Psmb8 | | | | Ndufa2 | |
| | | | Atp5o | | | | Eef2 | | | | Rps20 | |
| | | | Rbx1 | | | | Gm15920 | | | | Ywhab | |
| | | | Snx3 | | | | Ifi211 | | | | mt.Nd1 | |
| | | | Tcaf1 | | | | Cd14 | | | | Gnas | |
| | | | Cstb | | | | Hcar2 | | | | Tpm1 | |
| | | | BC028528 | | | | H2.T23 | | | | Atp5c1 | |
| | | | Gm2000 | | | | Bcl2a1a | | | | Fermt2 | |
| | | | Gm9392 | | | | Mif | | | | | |
| | | | Mrps5 | | | | | | | | | |
| | | | Uri1 | | | | | | | | | |
| | | | Gm8692 | | | | | | | | | |
| | | | Rpl7.ps7 | | | | | | | | | |
| | | | Dynll1 | | | | | | | | | |
| | | | Snrpb | | | | | | | | | |
| | | | Gm8606 | | | | | | | | | |
| | | | Srgn | | | | | | | | | |
| | | | Lrp8 | | | | | | | | | |
| | | | Gm5881 | | | | | | | | | |
| | | | Col4a2 | | | | | | | | | |
| | | | Anapc11 | | | | | | | | | |
| | | | Gm11361 | | | | | | | | | |
| | | | RP24.389J11.1 | | | | | | | | | |
| | | | Cox8a | | | | | | | | | |
| | | | Myh10 | | | | | | | | | |
| | | | Gm12966 | | | | | | | | | |

**Table S5.** Characteristics of myeloid clusters identified in DROP-seq data. Top 4 gene ontology (GO) terms were listed. Marker genes for each cluster were defined as differentially expressed genes between clusters at threshold: FDR <=0.01 and $lo_{g2}FC$>=0.5

Genes are listed from the highest to lowest $log_2FC$

| Cluster number | CL0 | CL1 | CL2 | CL3 | CL4 | CL5 | CL6 | CL7 | CL8 |
|---|---|---|---|---|---|---|---|---|---|
| Ontogeny | Ho-Mg | Ho-Mg | Mg-TAMs | Mg-TAMs | Mg-TAMs astrocytic-like | Mg-TAMs EC-like | Mg-TAMs cycling | Mo | BAMs |
| Top 4 GO terms | GO:1902075 cellular response to salt | GO:0008380 RNA splicing | GO:0002282 microglial cell activation involved in immune response | GO:0030595 leukocyte chemotaxis | GO:0042063 gliogenesis | GO:0001944 vasculature development | GO:0000278 mitotic cell cycle | GO:0045087 innate immune response | GO:1904646 cellular response to amyloid-beta |
| | GO:0030099 myeloid cell differentiation | GO:0048024 regulation of mRNA splicing, via spliceosome | GO:0030162 regulation of proteolysis | GO:0002252 immune effector process | GO:0007610 behavior | GO:0008360 regulation of cell shape | GO:0006325 chromatin organization | GO:0035456 response to interferon-beta | GO:0001816 cytokine production |
| | GO:0002521 leukocyte differentiation | GO:0060009 Sertoli cell development | GO:0055094 response to lipoprotein particle | GO:0006959 humoral immune response | GO:0031175 neuron projection development | GO:0030335 positive regulation of cell migration | GO:0051984 positive regulation of chromosome segregation | GO:0097193 intrinsic apoptotic signaling pathway | GO:0006954 inflammatory response |
| | GO:1901214 regulation of neuron death | GO:0048872 homeostasis of number of cells | GO:1905954 positive regulation of lipid localization | GO:1901796 regulation of signal transduction by p53 class mediator | GO:0055082 intracellular chemical homeostasis | GO:0045765 regulation of angiogenesis | GO:0006259 DNA metabolic process | GO:0019884 antigen processing and presentation of exogenous antigen | GO:0042742 defense response to bacterium |
| Marker genes | Jun | Luc7l3 | Cst7 | Spp1 | Sparcl1 | Ly6c1 | Mki67 | Ly6c2 | Mrc1 |
| | Egr1 | Malat1 | Spp1 | Cst7 | Clu | Cldn5 | Top2a | Plac8 | Dab2 |
| | P2ry12 | Prpf4b | Ccl6 | Ccl4 | Slc1a2 | Itm2a | Cenpf | H2.Eb1 | Pf4 |
| | Tmem119 | Luc7l2 | Cxcl13 | Cxcl13 | Ptn | Igfbp7 | Cenpe | Lyz2 | F13a1 |
| | Fos | Rbm5 | Lpl | Apoc1 | Mt2 | Abcb1a | Prc1 | Il1b | Ms4a7 |
| | Gpr34 | Pnn | Apoe | Tyrobp | Atp1a2 | Ptprb | Smc4 | Ms4a4c | Wfdc17 |
| | Ier5 | Arid4a | Cd63 | Ccl3 | Ttyh1 | Bsg | Hmmr | Ifitm3 | Cp |
| | Jund | Golgb1 | Tyrobp | Cd52 | Plpp3 | Adgrf5 | Smc2 | Lgals3 | Wwp1 |
| | Rhob | Cx3cr1 | Cd63.ps | Cd63 | Aldoc | Spock2 | Kif20b | Tgfbi | Maf |
| | Btg2 | Ythdc1 | Ctsd | Ch25h | Ttr | Flt1 | 2810417H13Rik | Ly6a | Stab1 |
| | Zfp36 | Smap2 | Ctsb | Ccl6 | Cpe | Ctla2a | Incenp | S100a11 | Gas6 |
| | Selplg | Akap9 | Creg1 | Cd9 | Scg3 | Slco1a4 | Tpx2 | Crip1 | Fcgr2b |
| | Cx3cr1 | Atrx | Cd9 | Ctsd | Ptprz1 | Slco1c1 | Birc5 | H2.Ab1 | Selenop |
| | Junb | Acin1 | mt.Nd1 | Rpl11 | Gja1 | Pltp | Ezh2 | Srgn | Lyz2 |
| | Siglech | Qk | Ctsz | Ctsz | Mt1 | Cxcl12 | Knl1 | Fxyd5 | Igf1 |
| | Cst3 | Arglu1 | mt.Cytb | Lpl | Aqp4 | Sptbn1 | Kif15 | AW112010 | Ehd4 |
| | Srgap2 | Ivns1abp | Mpeg1 | Ctsb | Slc1a3 | Slc2a1 | Nusap1 | Mndal | Pla2g7 |
| | Abca9 | Ncor1 | Npc2 | Ccl12 | Slc4a4 | Ly6a | Cdca8 | Isg15 | Cybb |
| | Il6ra | Prrc2c | Trem2 | Grn | Cald1 | Gng11 | Ckap2l | H2.Aa | Ms4a6b |
| | P2ry13 | Jmjd1c | Grn | Ctsl | Fxyd1 | Pcp4l1 | Ube2c | Msrb1 | Snx6 |
| | Olfml3 | Rdx | Ctsl | Rpl37 | Fabp7 | Sdpr | Kif23 | Gbp2 | Ifitm3 |
| | Ptgs1 | Pnisr | Rpl32 | Cd74 | Gpm6b | Esam | Aspm | Ifi204 | Ifi202b |
| | Ubc | Rsrc2 | mt.Nd5 | Cd68 | Mt3 | Ablim1 | Hmgb2 | Cdkn1a | Lgals1 |
| | Tgfbr1 | Phf14 | | Lgals3bp | Dbi | Slc39a10 | Diaph3 | Pim1 | Apoe |
| | Pde3b | | | C1qa | Ednrb | Egfl7 | Pbk | S100a6 | |
| | Ccr5 | | | Rpl31 | Htra1 | Tm4sf1 | Tuba1b | Slfn2 | |
| | Hpgds | | | Rpl13 | Acsl3 | Car4 | Cdk1 | Vim | |
| | Ivns1abp | | | Syngr1 | Gpr37l1 | Tsc22d1 | Cenpa | Cebpb | |
| | Elmo1 | | | Gm11942 | Cspg5 | Adgrl4 | Ccnb2 | Iqgap1 | |
| | Golm1 | | | Gm10260 | Ntrk2 | Kitl | Tubb5 | Cd14 | |
| | Mertk | | | Rpl19 | Prnp | Id3 | Mis18bp1 | Samhd1 | |
| | Zfhx3 | | | Rps19 | Atp1b2 | Ramp2 | Spc25 | Ifi211 | |
| | Rnase4 | | | Cd63.ps | Ndrg2 | Pecam1 | Clspn | Slfn5 | |
| | Arhgap5 | | | H2.D1 | Gstm1 | 9430020K01Rik | Kif11 | Tmsb10 | |
| | Mef2c | | | Rps24 | Nrxn1 | Mal | Lig1 | Ifi27l2a | |
| | Maf | | | Apoe | Syt11 | Col4a1 | Tyms | Ms4a6c | |
| | Scamp2 | | | Rpl13a | Bcan | Palmd | Tacc3 | Prdx5 | |
| | Mef2a | | | C1qc | Asrgl1 | Golim4 | Dek | Ms4a7 | |
| | Csf1r | | | Rps7 | Gria2 | Ccdc141 | Anln | Txn1 | |
| | Marcks | | | Gm14303 | Dclk1 | Itih5 | Ndc80 | Ms4a6b | |
| | Serinc3 | | | Rpl37a | Luzp2 | Edn1 | Cep55 | Calm1 | |
| | Lgmn | | | Rps15a | Scd2 | Slc16a1 | Nuf2 | Lilrb4a | |
| | Srrm2 | | | Ank | Lsamp | Epas1 | Tmpo | Cstb | |
| | Itgam | | | Hexa | S100b | Nes | Ncapg | Cd74 | |
| | Son | | | Rpl39 | Timp3 | Utrn | Gm11223 | Gm6977 | |
| | Malat1 | | | Eef1a1 | Prdx6 | Ppfibp1 | Atad2 | Clec7a | |
| | Hexb | | | Pld3 | Tmem47 | Selenom | Stmn1 | Cybb | |
| | Rbm39 | | | Fcer1g | S1pr1 | Fam13c | Cks1b | Lgals1 | |
| | | | | Mpeg1 | Id3 | Ptprg | Hirip3 | Fth1 | |
| | | | | Ctsh | Ntsr2 | Id1 | Sgol2a | Psmb8 | |
| | | | | Ndufa1 | Tspan7 | Sgms1 | Ccl12 | Ifi207 | |
| | | | | B2m | Fermt2 | Ptn | Nucks1 | Msr1 | |
| | | | | Rpl18a | Pla2g7 | Fn1 | Anp32b | Nampt | |
| | | | | | Fam213a | Timp3 | Bub1 | Sp140 | |
| | | | | | Mfge8 | Slc6a6 | Cbx5 | Tpt1.ps3 | |
| | | | | | Tuba1a | Arhgap29 | Selenoh | Ccr1 | |
| | | | | | Gpm6a | Cd59a | Pttg1 | Bcl2a1b | |
| | | | | | Nedd4 | Fermt2 | Arl6ip1 | Rpl34 | |
| | | | | | Sox2 | Kdr | Hist1h1e | Pla2g7 | |
| | | | | | S100a6 | Ets1 | Pcna | Bst2 | |
| | | | | | Ckb | Crip2 | Usp1 | Ifi202b | |
| | | | | | Tpm1 | Foxq1 | Tuba1c | Ifrd1 | |
| | | | | | Vim | App | Ranbp1 | Emp3 | |
| | | | | | Zbtb20 | Vwa1 | Anp32e | Wfdc17 | |
| | | | | | S100a1 | Abcg2 | Cxcl13 | Cxcl16 | |
| | | | | | Sdc4 | Cp | H2afz | Ifi30 | |
| | | | | | Gm42418 | Maoa | Gm4617 | Lst1 | |
| | | | | | Kif1b | Dynll1 | Whsc1 | Tagln2 | |
| | | | | | mt.Rnr1 | Aplp2 | Rangap1 | Stat1 | |
| | | | | | Myo6 | Ece1 | Hnrnpab | Fcgr2b | |
| | | | | | Tspan3 | Col4a2 | Ran | Shfm1 | |
| | | | | | Sept7 | Ifitm3 | Ckap5 | H2.K1 | |
| | | | | | Ldhb | Apod | Rad21 | Psme2b | |
| | | | | | Slc25a4 | Fnbp1l | Ccdc34 | Rps2 | |
| | | | | | Cbx5 | Alas1 | Tagln2 | Anxa5 | |
| | | | | | Agpat5 | Iqgap1 | Gm8203 | Hspa8 | |
| | | | | | Psip1 | Ifitm2 | Hjurp | Psmb9 | |
| | | | | | Nfia | Osbpl1a | Lgals1 | H2.DMa | |
| | | | | | Add3 | Tceb2 | Nasp | Ms4a6d | |
| | | | | | Ccdc88a | Actn4 | Dnajc9 | Sh3bgrl3 | |
| | | | | | Sptbn1 | Ctnnb1 | Anp32b.ps1 | Btg1 | |
| | | | | | Cfap36 | Tpm1 | Fam111a | Tspo | |
| | | | | | Serpine2 | Prnp | Hpf1 | Sh3bgrl | |
| | | | | | Glud1 | Grap | Tubb4b | Mcl1 | |
| | | | | | Skp1a | Igf1r | Pcna.ps2 | H2.T23 | |
| | | | | | Gm12346 | Wbp5 | H2afv | Rps18 | |
| | | | | | | Tmsb10 | Ifi27l2a | Ccrl2 | |
| | | | | | | Tspan13 | Smchd1 | Taldo1 | |
| | | | | | | Hip1 | Smc1a | Cd52 | |
| | | | | | | Col4a3bp | Hmgn1 | Rpl35a | |
| | | | | | | Afdn | Txn1 | Trim30a | |
| | | | | | | Luzp1 | Lmnb1 | Tpm4 | |
| | | | | | | Sparcl1 | Gm4204 | Cyba | |
| | | | | | | Snx3 | Cpd | Pfn1 | |
| | | | | | | Mt2 | Smc6 | Ldha | |
| | | | | | | Rpl36a.ps2 | Gm9800 | Prdx6 | |
| | | | | | | Anxa2 | Cdkn1a | H2.DMb1 | |
| | | | | | | S100a13 | Adamts1 | Gm9794 | |
| | | | | | | Myl12b | Ncl | Alox5ap | |
| | | | | | | Nedd4 | Psip1 | B2m | |
| | | | | | | Tmem256 | Tpr | Ptprc | |
| | | | | | | Rps6.ps4 | Nop58 | Rps27l | |
| | | | | | | BC002163 | Snrpd1 | Pkm | |
| | | | | | | Calm1 | Larp7 | Rps15a | |
| | | | | | | Igf1 | Xpo1 | Gm15427 | |

| | | | | | | | | |
|---|---|---|---|---|---|---|---|---|
| | | | | | | Myl12a | Aimp1 | Rps24.ps3 |
| | | | | | | Atox1 | Uchl5 | Rps13.ps2 |
| | | | | | | Cuta | Hn1 | Rps3 |
| | | | | | | Dynlt3 | Smarcc1 | Psme2 |
| | | | | | | Rpl38.ps2 | Ssrp1 | Rps16.ps2 |
| | | | | | | Gm8186 | Prdx1 | Myl12a |
| | | | | | | Crip1 | Banf1 | H2.D1 |
| | | | | | | Gm7809 | Rif1 | Psma7 |
| | | | | | | Hes1 | Brd8 | Mif |
| | | | | | | Selenop | Tpm4 | Prdx1 |
| | | | | | | Gm5436 | Ybx1 | Rplp0 |
| | | | | | | Ahnak | Supt16 | Sdcbp |
| | | | | | | Syne1 | Ifi30 | Actr3 |
| | | | | | | Gm8894 | Snrpa1 | Arpc5 |
| | | | | | | Gas6 | Pa2g4 | Psma2 |
| | | | | | | Shfm1 | Ptma | Gm9843 |
| | | | | | | Gm6265 | Fkbp3 | Wdr89 |
| | | | | | | Ctnna1 | Hnrnpu | Cox17 |
| | | | | | | Gba | Pbrm1 | Sub1 |
| | | | | | | Cd34 | Cox5a | Rel |
| | | | | | | Ndufb5 | | Ctsc |
| | | | | | | Ifnar1 | | Rpl39 |
| | | | | | | Cfdp1 | | Gm6472 |
| | | | | | | Glud1 | | Rps9 |
| | | | | | | Lima1 | | Rpl22 |
| | | | | | | Gm4617 | | H3f3b |
| | | | | | | Atp5j | | Lrrfip1 |
| | | | | | | Gm3531 | | Gm5905 |
| | | | | | | Capns1 | | Sec61b |
| | | | | | | Atp5d | | Cd47 |
| | | | | | | Uqcc2 | | Gm9385 |
| | | | | | | Arhgef12 | | Gm12254 |
| | | | | | | Nampt | | Rpl18.ps1 |
| | | | | | | Gm4149 | | Lcp1 |
| | | | | | | | | Chmp4b |
| | | | | | | | | Rps15a.ps5 |
| | | | | | | | | Rps26 |
| | | | | | | | | Fosb |
| | | | | | | | | Rps14 |
| | | | | | | | | Gpx1 |
| | | | | | | | | Gm4332 |
| | | | | | | | | Gm5963 |
| | | | | | | | | Arpc3 |
| | | | | | | | | Aldoa |
| | | | | | | | | Gm15500 |
| | | | | | | | | Rps10.ps2 |
| | | | | | | | | Rpl36al |
| | | | | | | | | Myl12b |
| | | | | | | | | Psme1 |
| | | | | | | | | Rpl37rt |
| | | | | | | | | Gm6030 |
| | | | | | | | | Gm5805 |
| | | | | | | | | Hif1a |
| | | | | | | | | Gm5835 |
| | | | | | | | | Rps3a1 |
| | | | | | | | | Atox1 |
| | | | | | | | | Capg |
| | | | | | | | | Slk |
| | | | | | | | | Rpl38.ps2 |
| | | | | | | | | Ly6e |
| | | | | | | | | Neat1 |
| | | | | | | | | Shisa5 |
| | | | | | | | | Gm10250 |
| | | | | | | | | Psap |
| | | | | | | | | Rps5 |
| | | | | | | | | Nfe2l2 |
| | | | | | | | | Gm8995 |
| | | | | | | | | Gm6204 |
| | | | | | | | | Gm8730 |
| | | | | | | | | Rack1 |
| | | | | | | | | Rpl9.ps6 |
| | | | | | | | | Rps10.ps1 |
| | | | | | | | | Nfkbia |
| | | | | | | | | Rpsa.ps10 |
| | | | | | | | | Rps11 |
| | | | | | | | | Npc2 |
| | | | | | | | | Fcer1g |
| | | | | | | | | Gm14586 |
| | | | | | | | | Rps20 |
| | | | | | | | | Rps19 |
| | | | | | | | | Litaf |
| | | | | | | | | Arpc2 |
| | | | | | | | | Gm4149 |
| | | | | | | | | Tpm3.rs7 |
| | | | | | | | | Rpl8 |
| | | | | | | | | Tax1bp1 |
| | | | | | | | | Rps21 |
| | | | | | | | | Atp5h |
| | | | | | | | | Rpl36a.ps2 |
| | | | | | | | | Gm10275 |
| | | | | | | | | Rpl31.ps8 |
| | | | | | | | | Cox7a2 |
| | | | | | | | | Rpl4 |
| | | | | | | | | Spop |
| | | | | | | | | Rpl14.ps1 |
| | | | | | | | | Gm4204 |
| | | | | | | | | Rps15 |
| | | | | | | | | Chmp2a |
| | | | | | | | | Gm11478 |

**Table S6.** Lists of differentially expressed genes between PDOXs,GL261 and their respective normal brain controls (Nu-NB, B6-NB) per each myeloid cell type.
Differentially expressed genes were defined at threshold: FDR <=0.01 and |log2FC|>=1
Genes are listed from the highest to lowest |log2FC|

| Cell type | Microglia (Mg) | | | | | | Peripheral monocyte-derived cells (Mo) | | | | | | Border associated macrophages (BAMs) | | | | | |
|---|---|---|---|---|---|---|---|---|---|---|---|---|---|---|---|---|---|---|
| comparison | PDOX vs GL261 | | PDOX vs Nu-NB | | GL261 vs B6-NB | | PDOX vs GL261 | | PDOX vs Nu-NB | | GL261 vs B6-NB | | PDOX vs GL261 | | PDOX vs Nu-NB | | GL261 vs B6-NB | |
| Fold change | upregulated | downregulated | upregulated | downregulated | upregulated | downregulated | upregulated | downregulated | upregulated | downregulated | upregulated | downregulated | upregulated | downregulated | upregulated | downregulated | upregulated | downregulated |
| Number of genes | 31 | 39 | 171 | 68 | 108 | 36 | 47 | 68 | 81 | 45 | 0 | 96 | 7 | 53 | 46 | 67 | 0 | 0 |
| | Cxcl13 | H2.Eb1 | Spp1 | Sfrs18 | Cd74 | Ttr | Cxcl13 | H2.Eb1 | Spp1 | Gm26924 | | Ttr | Ifi202b | H2.Eb1 | Selenop | Sepp1 | | |
| | Ttr | Plp1 | Gm42418 | Snord49b | H2.Ab1 | Sparcl1 | Ifi202b | Thbs1 | Gm6977 | Sepp1 | | Cd209a | Gm11942 | Cd14 | Gm14303 | Rps29 | | |
| | Lpl | Rpl34 | Cxcl13 | Gm26924 | H2.Eb1 | P2ry12 | Sparc | Arg1 | Clec7a | Rpl5 | | Ighm | Mef2c | Rsad2 | Gm6977 | Sfrs18 | | |
| | Spp1 | Hspa8 | Clec7a | 4632428N05Rik | H2.Aa | Gpr34 | Ftl1 | Hspa8 | Cxcl13 | Sfrs18 | | Cd244 | Jun | Cdkn1a | Clec7a | Gm26924 | | |
| | Ifi202b | Rps2 | Cd74 | Sepp1 | Il1b | Hpgd | Lpl | Rpl34 | Gm5905 | Rps27a | | Nemp2 | Cst3 | Arg1 | Gm5905 | D4Wsu53e | | |
| | Cwc22 | H2.Q7 | H2.Aa | Epb4.1l2 | Lyz2 | Selplg | Hexb | Cxcl2 | Gm42418 | Jhdm1d | | Tnrc6c | mt.Cytb | Cxcl10 | Gm42418 | AI607873 | | |
| | Fn1 | Fosb | Gm6977 | Gm26339 | Ifi27l2a | Rgs2 | Gm11942 | Rps2 | Gm9794 | Gm17087 | | H2.K2 | Malat1 | Rpl34 | Rpl14.ps1 | RP24.288C12.6 | | |
| | Ank | Rps18 | Gm14303 | Entpd1 | Ccl12 | Tmem119 | Cst7 | Rps18 | Gm14303 | Rps23 | | Gm19815 | | Ubl3 | H2.Aa | Rps12 | | |
| | Ccl6 | Bst2 | Gm5905 | Abca9 | Clec7a | Tgfbr1 | Cwc22 | Vegfa | Rpl37rt | AI607873 | | Myrf | | Ly6a | Gm9794 | Vimp | | |
| | Gm11942 | Cebpb | Ccl4 | D4Wsu53e | H2.D1 | Mertk | Ctsd | Bst2 | Rpl14.ps1 | Rpl35 | | Phtf2 | | Tgfbi | Gm15427 | Cmah | | |
| | Ch25h | Gbp2 | Gm9794 | Gpr34 | Ifitm3 | Rnase4 | Cx3cr1 | Fosl2 | Gm15427 | Rps29 | | Pou2f2 | | Rps2 | Gm11478 | Gm26339 | | |
| | Ftl1 | Nr4a1 | H2.Ab1 | P2ry13 | H2.K1 | Maf | Siglech | Cebpb | Ctsd | Lrrc58 | | Zfp329 | | Rps18 | Cd63.ps | Lilrb4 | | |
| | Apoe | Gbp7 | Ttr | P2ry12 | Apoe | Golm1 | Ttr | Cd14 | Cd63.ps | Rpl19 | | Ccp110 | | Ly6c2 | Gm4149 | Cd163 | | |
| | Itgax | Fos | Lpl | Maf | Xist | Csf1r | Ifi44 | Ptgs2 | Gm11478 | Rps12 | | Pla2g12a | | Etf1 | Rpl37rt | Rps27a | | |
| | Sparcl1 | Atf3 | Rpl14.ps1 | Jund | Vim | Cx3cr1 | Serpine2 | Fosb | Rps16.ps2 | Rpl3 | | Spc25 | | Rpl30 | Gm11942 | Rpl10 | | |
| | Gm10260 | Rpl35a | Lyz2 | Zfhx3 | Gbp2 | Dusp6 | Cst3 | Il1b | Gm4149 | Rps6 | | Tlr7 | | Iigp1 | Gm5963 | Rps23 | | |
| | Arsb | Ldhb | Rpl37rt | Nrip1 | Cd83 | Ifngr1 | Gm10260 | Nr4a2 | Rack1 | Trim30a | | Wiz | | Rpl35a | Rps24.ps3 | Snord49b | | |
| | Ptn | Ly6a | Fth1 | Slco2b1 | Ccl4 | Scoc | Cd81 | H2.Q7 | Rps24.ps3 | Diap2 | | Megf9 | | Il1b | Rplp1 | F13a1 | | |
| | mt.Co1 | Il1a | Gm15427 | Tmem119 | Nr4a1 | P2ry13 | mt.Cytb | S100a11 | Gm9385 | Ppp1r12a | | Siglech | | Bst2 | Gm9385 | Crebrf | | |
| | mt.Cytb | Ccl12 | Ccl3 | Hpgds | H2.Q7 | Fcrls | Trem2 | AA467197 | Gm5963 | Arhgap30 | | Ctnnal1 | | Hspa8 | Tpt1.ps3 | Ccnd2 | | |
| | Cd34 | Cd83 | Gm11478 | Il6ra | Ly6a | Siglech | Mef2c | Clec4n | Ccl4 | Fau | | Hspa1b | | Nrp2 | Wdr89 | Med13 | | |
| | Axl | Fcrls | Cd63.ps | Cx3cr1 | Cd52 | Jun | Rnase4 | Nr4a3 | Trem2 | Erbb2ip | | Ate1 | | Ifi209 | Rps16.ps2 | Myl6 | | |
| | Cadm1 | H2.Ab1 | Rack1 | Mertk | Il1a | Rhob | mt.Co1 | Rpl29 | Tpt1.ps3 | Rps27 | | Stap1 | | Ccl2 | Gm5805 | Fau | | |
| | Scd2 | Ccl4 | Gm16580 | Smap2 | Ifi204 | Cst3 | Golm1 | Nlrp3 | Gm11942 | Rpl17 | | Myo18a | | Cebpb | Pnisr | Zc3h7a | | |
| | Xist | Dusp1 | Gm11942 | Ppp1r9a | Fth1 | Ivns1abp | Ccl3 | Ifrd1 | Lilrb4a | Rpl21 | | Zfp207 | | Lgals3 | Rpl3.ps1 | Stard8 | | |
| | Ctsd | C1qb | Gm5963 | Arid4a | Bst2 | Serpine2 | Creg1 | Tgfbi | Gm5805 | Rab8b | | Mef2c | | Prdx5 | Gm10073 | Ptpn6 | | |
| | mt.Nd4 | Ifrd1 | Cst7 | Son | Ifrd1 | Il6ra | Spp1 | Ly6a | Rpl3.ps1 | Rpl27 | | Dmxl1 | | Aldoa | Gm10288 | Ralgps2 | | |
| | mt.Nd2 | Cd14 | Apoe | Tanc2 | Rpl32 | Elmo1 | mt.Nd1 | Ccr5 | Gm16580 | Gnb2l1 | | Mum1 | | Lyz2 | Rps13.ps2 | Abi1 | | |
| | Gnas | Ifi204 | Selenop | Chd9 | Txn1 | Pip4k2a | mt.Nd4 | Ccr1 | Rps13.ps2 | Rpl13 | | Cuedc1 | | Ckap5 | Lilrb4a | Ppia | | |
| | mt.Nd5 | Nfkbia | Gm4149 | Srgap2 | Rpl39 | Sirpa | Cadm1 | Dab2 | Gm6204 | Rpl18a | | Ankrd49 | | Pim1 | Rsrp1 | Kmt2c | | |
| | mt.Nd1 | C1qc | Rps24.ps3 | Ogfrl1 | Stat1 | Plxdc2 | Jun | Gpnmb | Fth1 | Tpt1 | | Plxdc2 | | Mndal | Cd52 | Tab2 | | |
| | | Il1b | Gm9385 | Scamp2 | Nfkbia | Fam105a | Rpl13a | Osm | Rpl36a.ps2 | Mrps25 | | Clock | | Fosb | Rack1 | Iqgap2 | | |
| | | Mcl1 | Rps16.ps2 | Elmo1 | Tmsb10 | Rbm5 | Cd9 | Gm9493 | Gm4332 | Rpl13a | | Faap100 | | Vim | Rpl18.ps1 | Cwc22 | | |
| | | Txnip | Gm5805 | Mycbp2 | Rps20 | Vsir | mt.Nd2 | Mndal | Cst7 | Sarnp | | Slco2b1 | | Psap | Mt1 | Nrp1 | | |
| | | H2.DMa | Rpl3.ps1 | Lpar6 | Plp1 | Hpgds | Egr1 | Rpl3 | Gm8995 | Rpl10 | | Tcf3 | | Srgn | Rpl36a.ps2 | Map4k4 | | |
| | | Ly86 | Pnisr | Rbm5 | Cstb | Hexb | Ctsl | Crem | Rps10.ps1 | Ptp4a2 | | Znrf2 | | Ccl7 | Ttr | Abca9 | | |
| | | Ubc | Ccl6 | Ssh2 | Nfkbiz | | Hpgds | Lyz2 | Rps26 | Rpl7a | | Acss2 | | Txn1 | Gm6204 | Sbds | | |
| | | Rpl3 | Fn1 | Marcks | Rps2 | | Gnas | Tnfaip2 | Gm10073 | Ifi203 | | Dhx33 | | Gm16580 | Slc25a5 | Gm17087 | | |
| | | Socs3 | Rps13.ps2 | Qk | Lgals3bp | | Abhd12 | Nr4a1 | Gm14586 | Rpl15 | | Tmem183a | | Samd9l | Rps5 | Rps6 | | |
| | | | Il1b | Macf1 | Slfn2 | | Mtdh | Cdkn1a | Gm4617 | Prrc2c | | Sp3 | | Ccr1 | Fth1 | Fam111a | | |
| | | | Gm6204 | Arl4c | Ccl3 | | Apbb1ip | Plaur | Rgs1 | Rpl23a | | Eif3s6.ps2 | | Nfe2l2 | Gm9843 | Emp3 | | |
| | | | Ifi27l2a | Ythdc1 | Rps26 | | Man2b1 | Rpl30 | Lpl | Ptma | | Trp53inp1 | | Rnasel | Npc2 | Aldh2 | | |
| | | | Gm8995 | Cwc22 | Atf3 | | Ccl4 | Slamf7 | Rpl10a.ps1 | D4Wsu53e | | Plxna4os1 | | Cox5a | H2.D1 | Mcfd2 | | |
| | | | Cd52 | Frmd4a | Anp32b | | Cd63 | Trim25 | Rpl18.ps1 | Lyz1 | | Cc2d1b | | Psmb9 | B2m | Cfh | | |
| | | | Kctd12 | Dusp6 | B2m | | Rps4x | Tnfaip3 | Gm15500 | Clec12a | | Trerf1 | | Fth1 | Fcer1g | Ctage5 | | |
| | | | H2.K1 | Olfml3 | Mif | | Apoe | Fgl2 | Cd63 | | | Ttf1 | | H2.Ab1 | Actb | Hmgb1 | | |
| | | | Rpl36a.ps2 | Prrc2c | Rpl9.ps6 | | Rgs1 | Tiparp | mt.Nd1 | | | Pik3cd | | Pomp | | Sgpl1 | | |
| | | | Epb41l2 | Srrm2 | Rpl37 | | | Pim1 | Gm6030 | | | Gm6564 | | C1qb | | Rabep1 | | |
| | | | Ch25h | Ifngr1 | Ncl | | | Mxd1 | Rps25.ps1 | | | 1110051M20Rik | | Sh3bgrl3 | | Dusp6 | | |
| | | | Rpl10a.ps1 | Ivns1abp | Rpl41 | | | Pde4b | Rpsa.ps10 | | | Igip | | H2.K1 | | Cx3cr1 | | |
| | | | Rps5 | Mtdh | Rplp1 | | | Il1rn | mt.Cytb | | | Rfx5 | | Calm1 | | Ppp1r9a | | |
| | | | Rsrp1 | Acin1 | Rack1 | | | Ets2 | Rps5 | | | Hpgd | | Stat1 | | Tlr7 | | |
| | | | Gm14586 | Arhgap5 | Gm10288 | | | Nfil3 | Gm5835 | | | Gm15207 | | B2m | | Hnrnpl | | |
| | | | Gm4617 | Ccr5 | Rpl22 | | | Cytip | Ctsl | | | Slc12a2 | | | | Plekho1 | | |
| | | | Ctsd | Selplg | Rpl14 | | | Tgm2 | Rsrp1 | | | Zfp770 | | | | Mrc1 | | |
| | | | Gm4332 | Malat1 | Tspo | | | Acp5 | Rpl31.ps8 | | | Fam149a | | | | Gpr183 | | |
| | | | Ifi202b | Pnn | Srgn | | | H2.Ab1 | Rps15a.ps5 | | | Nfix | | | | Maf | | |
| | | | Rpsa.ps10 | Pde3b | Cd14 | | | Cyp4f18 | Cd81 | | | Nupl1 | | | | Rpl13a | | |
| | | | Rps26 | Tgfbr1 | Rpl31.ps8 | | | Dusp1 | Sparc | | | Slc1a3 | | | | Acin1 | | |
| | | | Mt1 | Rbm25 | Psmb8 | | | Clic4 | Gm6863 | | | Poc1b | | | | Mndal | | |
| | | | H2.D1 | Pmepa1 | Rps10.ps1 | | | Rps24.ps2 | Rplp0 | | | Il16 | | | | Zfp36l2 | | |
| | | | Rps25.ps1 | Anp32a | Rps8 | | | Tmsb10 | H2.K1 | | | Enpp4 | | | | Rin2 | | |
| | | | Gm10288 | Basp1 | Rps3 | | | Klf4 | Gm10275 | | | Syne2 | | | | Fus | | |
| | | | Calr.ps | Serinc3 | Rps10 | | | Rpl35a | Gnas | | | Strip1 | | | | Hspa8 | | |
| | | | Gm6030 | Srrm1 | Rps25.ps1 | | | Bcl2a1b | Rps15 | | | Ipo9 | | | | Pdia3 | | |
| | | | Cybb | Ywhah | Gm9794 | | | Vim | Apoe | | | Vps52 | | | | D8Ertd738e | | |
| | | | Wdr89 | Jmjd1c | Mcl1 | | | Emb | mt.Nd4 | | | Slc44a2 | | | | C1qa | | |
| | | | Ctsb | Ddx5 | Rps19 | | | Lgals3 | Rplp1 | | | Dleu2 | | | | | | |
| | | | Rpl31.ps8 | | Wdr89 | | | | Ctsb | | | Rbm25 | | | | | | |
| | | | AU020206 | | Ccrl2 | | | | Slc25a5 | | | Gm10064 | | | | | | |
| | | | Tyrobp | | Cebpb | | | | Gm9843 | | | D17H6S53E | | | | | | |
| | | | Ccrl2 | | Rps24 | | | | Atox1 | | | Rabggta | | | | | | |
| | | | Gm15500 | | Gm4332 | | | | B2m | | | Cfh | | | | | | |
| | | | Gm9843 | | Hint1 | | | | Npc2 | | | Map4k5 | | | | | | |
| | | | Gm5835 | | Gm14303 | | | | Rps3 | | | Rrad | | | | | | |
| | | | Tpt1.ps3 | | Gm5963 | | | | Rpl32 | | | Dopey1 | | | | | | |
| | | | Gm10073 | | Rplp0 | | | | Tyrobp | | | Spata13 | | | | | | |
| | | | Rpl18.ps1 | | Rpl8 | | | | Mpeg1 | | | Rpl5 | | | | | | |
| | | | Gm8276 | | Rpl37rt | | | | Wdr89 | | | Tesk1 | | | | | | |
| | | | Rpl32 | | Rps15 | | | | Aif1 | | | Fam120b | | | | | | |
| | | | Vsir | | Rps26.ps1 | | | | Ctsz | | | Usp32 | | | | | | |
| | | | Axl | | Prdx1 | | | | | | | Prnp | | | | | | |
| | | | Rps20 | | Rps15a | | | | | | | Bbc3 | | | | | | |
| | | | Rplp1 | | Gbp7 | | | | | | | Zfp120 | | | | | | |
| | | | Itgax | | Rps24.ps3 | | | | | | | Gm37494 | | | | | | |
| | | | Id2 | | Tubb5 | | | | | | | Ube3b | | | | | | |
| | | | Rps10.ps1 | | Egr1 | | | | | | | Gpm6b | | | | | | |
| | | | mt.Cytb | | Fosb | | | | | | | Dhx57 | | | | | | |
| | | | mt.Nd1 | | Gm16580 | | | | | | | Wdr60 | | | | | | |
| | | | Gm14681 | | Rps21 | | | | | | | Tppp3 | | | | | | |
| | | | Gm6863 | | Gm6977 | | | | | | | RP23.394G23.2 | | | | | | |
| | | | Ms4a6c | | Ifi30 | | | | | | | Gm7589 | | | | | | |
| | | | Cd83 | | Rpl23 | | | | | | | Sort1 | | | | | | |
| | | | Gnas | | Bcl2a1b | | | | | | | Hspa1a | | | | | | |
| | | | Gm10275 | | Eef2 | | | | | | | E330020D12Rik | | | | | | |
| | | | Ank | | Gm6030 | | | | | | | Rufy2 | | | | | | |
| | | | Cd63 | | Rps5 | | | | | | | | | | | | | |
| | | | Atox1 | | Npm1 | | | | | | | | | | | | | |
| | | | Ctsz | | Pkm | | | | | | | | | | | | | |
| | | | Atf3 | | Pabpc1 | | | | | | | | | | | | | |
| | | | B2m | | Rpl34 | | | | | | | | | | | | | |
| | | | Nfkbia | | Plek | | | | | | | | | | | | | |
| | | | Atp5e | | Rpl36a.ps2 | | | | | | | | | | | | | |
| | | | Srgn | | Rps13.ps2 | | | | | | | | | | | | | |
| | | | Calm1 | | H2.T23 | | | | | | | | | | | | | |
| | | | Cox6a1 | | Gatm | | | | | | | | | | | | | |
| | | | Rps26.ps1 | | Rpl4 | | | | | | | | | | | | | |
| | | | Cstb | | Eef1b2 | | | | | | | | | | | | | |
| | | | Npc2 | | | | | | | | | | | | | | | |
| | | | Dbi | | | | | | | | | | | | | | | |
| | | | Rpl39 | | | | | | | | | | | | | | | |
| | | | Cox4i1 | | | | | | | | | | | | | | | |
| | | | Ctsl | | | | | | | | | | | | | | | |
| | | | Rgs1 | | | | | | | | | | | | | | | |
| | | | Grn | | | | | | | | | | | | | | | |
| | | | Prdx1 | | | | | | | | | | | | | | | |
| | | | Rps15 | | | | | | | | | | | | | | | |
| | | | Csf2ra | | | | | | | | | | | | | | | |
| | | | Cox6b1 | | | | | | | | | | | | | | | |
| | | | mt.Nd4 | | | | | | | | | | | | | | | |
| | | | Gusb | | | | | | | | | | | | | | | |
| | | | Lgals3bp | | | | | | | | | | | | | | | |
| | | | Pabpc1 | | | | | | | | | | | | | | | |
| | | | Pld3 | | | | | | | | | | | | | | | |
| | | | Ppp1r15a | | | | | | | | | | | | | | | |
| | | | Rplp0 | | | | | | | | | | | | | | | |
| | | | Tmsb10 | | | | | | | | | | | | | | | |
| | | | Rps19 | | | | | | | | | | | | | | | |
| | | | Gpx1 | | | | | | | | | | | | | | | |
| | | | Cox8a | | | | | | | | | | | | | | | |
| | | | Fcer1g | | | | | | | | | | | | | | | |
| | | | Rps9 | | | | | | | | | | | | | | | |
| | | | Cd9 | | | | | | | | | | | | | | | |
| | | | Slc25a5 | | | | | | | | | | | | | | | |
| | | | Nfkbiz | | | | | | | | | | | | | | | |
| | | | Gm8730 | | | | | | | | | | | | | | | |
| | | | Rps3 | | | | | | | | | | | | | | | |
| | | | Rps11 | | | | | | | | | | | | | | | |
| | | | Akr1a1 | | | | | | | | | | | | | | | |
| | | | Cotl1 | | | | | | | | | | | | | | | |
| | | | Sh3bgrl3 | | | | | | | | | | | | | | | |
| | | | Uqcrq | | | | | | | | | | | | | | | |
| | | | Ier2 | | | | | | | | | | | | | | | |
| | | | Gm6472 | | | | | | | | | | | | | | | |
| | | | Ndufa2 | | | | | | | | | | | | | | | |
| | | | Ssr4 | | | | | | | | | | | | | | | |
| | | | Shfm1 | | | | | | | | | | | | | | | |
| | | | Ctsh | | | | | | | | | | | | | | | |
| | | | Psme2b | | | | | | | | | | | | | | | |
| | | | Eif3f | | | | | | | | | | | | | | | |
| | | | Rpl22l1 | | | | | | | | | | | | | | | |
| | | | Myo5a | | | | | | | | | | | | | | | |
| | | | Rps21 | | | | | | | | | | | | | | | |
| | | | Rps14 | | | | | | | | | | | | | | | |
| | | | Rpl8 | | | | | | | | | | | | | | | |
| | | | 2010107E04Rik | | | | | | | | | | | | | | | |
| | | | Gm10076 | | | | | | | | | | | | | | | |
| | | | Arpc1b | | | | | | | | | | | | | | | |
| | | | Atp6v1f | | | | | | | | | | | | | | | |
| | | | Cox6c | | | | | | | | | | | | | | | |
| | | | Eef2 | | | | | | | | | | | | | | | |
| | | | Pfdn5 | | | | | | | | | | | | | | | |
| | | | Cd68 | | | | | | | | | | | | | | | |
| | | | Myeov2 | | | | | | | | | | | | | | | |
| | | | Mpeg1 | | | | | | | | | | | | | | | |
| | | | Itgb2 | | | | | | | | | | | | | | | |
| | | | Cox7a2 | | | | | | | | | | | | | | | |
| | | | Ndufa1 | | | | | | | | | | | | | | | |
| | | | Tmem256 | | | | | | | | | | | | | | | |
| | | | Rpl41 | | | | | | | | | | | | | | | |
| | | | Atp5j2 | | | | | | | | | | | | | | | |

**Table S7.** List of gene signatures applied for scRNA-seq analysis.

| Signature | Human Mg | Human Mo | Enzymatic activation | Homeostatic Mg | Sensome | Sensome w/o homeostatic Mg | Migration | APC | Phagocytosis |
|---|---|---|---|---|---|---|---|---|---|
| Genes | 12 | 13 | 25 | 35 | 140 | 127 | 464 | 106 | 219 |
| | HEXB | CCR2 | Rgs1 | Bhlhe41 | 4632428N05Rik | Abcc3 | Abl1 | Abcb9 | 4932438A13Rik |
| | P2RY12 | PLAC8 | Hist2h2aa1 | Crybb1 | Abcc3 | Adora3 | Acvrl1 | Ap3b1 | 4933434E20Rik |
| | P2RY13 | CLEC12A | Hist1h4i | Csf1r | Adora3 | Adrb2 | Adam17 | Ap3d1 | Abca1 |
| | TMEM119 | FCN1 | Nfkbiz | Ctsd | Adrb2 | Bin2 | Adam8 | Arl8b | Abca7 |
| | GPR34 | VCAN | Klf2 | Ctsf | AF251705 | C3ar1 | Adam9 | Atg5 | Abl1 |
| | CXCR1 | LYZ | Junb | Cttnbp2 | Bin2 | C5ar1 | Adamts9 | B2m | Abl2 |
| | TGFBR1 | LGALS3 | Dusp1 | Cttnbp2nl | C3ar1 | Ccr5 | Adgra2 | Bag6 | Abr |
| | CST3 | CRIP1 | Ccl3 | Cx3cr1 | C5ar1 | Ccrl2 | Adgrb1 | Calr | Adgrb1 |
| | OLFML3 | S100A4 | Hspa1a | Fcrls | C5ar2 | Cd14 | Adgrg1 | Ccr7 | Aif1 |
| | SALL1 | S100A6 | Hsp90aa1 | Golm1 | Ccr5 | Cd180 | Adipor1 | Cd1d1 | Ano6 |
| | SELPLG | S100A8 | Fos | Gpr34 | Ccrl2 | Cd22 | Adora2b | Cd68 | Anxa1 |
| | TREM2 | S100A9 | Hspa1b | Gpr84 | Cd101 | Cd300a | Agt | Cd74 | Anxa3 |
| | | CD44 | Jun | Hexb | Cd14 | Cd33 | Aif1 | Clec4a1 | Appl1 |
| | | | Jund | Hpgds | Cd180 | Cd37 | Akirin1 | Clec4a2 | Appl2 |
| | | | Nfkbid | Hspa1a | Cd22 | Cd48 | Akt1 | Clec4a3 | Arhgap12 |
| | | | Gem | Lgmn | Cd300a | Cd52 | Akt3 | Ctsl | Arhgap25 |
| | | | Ccl4 | Lrrc3 | Cd33 | Cd53 | Amot | Ctss | Arl8b |
| | | | Ier5 | Ltc4s | Cd37 | Cd68 | Amotl1 | Erap1 | Atg3 |
| | | | Txnip | Mafb | Cd48 | Cd74 | Angpt1 | Ext1 | Atg5 |
| | | | Hist1h2bc | Nuak1 | Cd52 | Cd83 | Angpt2 | Fcer1g | Atg7 |
| | | | Zfp36 | Olfml3 | Cd53 | Cd84 | Anln | Fcgr1 | Axl |
| | | | Hist1h1c | P2ry12 | Cd68 | Cd86 | Ano6 | Fcgr2b | Bcr |
| | | | Egr1 | P2ry13 | Cd74 | Clec4a2 | Anxa1 | Fcgr3 | Becn1 |
| | | | Atf3 | Plxdc2 | Cd79b | Clec4a3 | Anxa3 | Fcgr4 | Bin2 |
| | | | Rhob | Rab3il1 | Cd83 | Clec5a | Apoe | Fgl2 | C2 |
| | | | | Sall1 | Cd84 | Clec7a | App | Gba | C3 |
| | | | | Scoc | Cd86 | Cmklr1 | Aqp1 | Gm8909 | C4b |
| | | | | Serpine2 | Clec4a2 | Cmtm6 | Arf6 | H2.Aa | Calr |
| | | | | Siglech | Clec4a3 | Cmtm7 | Arl13b | H2.Ab1 | Camk1d |
| | | | | Slc2a5 | Clec4b1 | Csf2rb2 | Arsb | H2.D1 | Ccl2 |
| | | | | Tgfbr1 | Clec5a | Csf3r | Atoh8 | H2.DMa | Ccr2 |
| | | | | Tmem119 | Clec7a | Cxcl16 | Atp2b4 | H2.DMb1 | Ccr7 |
| | | | | Atp8a2 | Cmklr1 | Cysltr1 | Atp5a1 | H2.DMb2 | Cd300a |
| | | | | Lag3 | Cmtm6 | Daglb | Atp5b | H2.Eb1 | Cd300lf |
| | | | | Slco4a1 | Cmtm7 | Ecscr | Axl | H2.K1 | Cd302 |
| | | | | | Csf1r | Entpd1 | Bcar1 | H2.M3 | Cd36 |
| | | | | | Csf2rb2 | Fcer1g | Bcas3 | H2.Oa | Cd47 |
| | | | | | Csf3r | Fcgr1 | Bmpr2 | H2.Q10 | Cdc42 |
| | | | | | Cx3cr1 | Fcgr2b | Bsg | H2.Q4 | Cdc42se1 |
| | | | | | Cxcl16 | Fcgr3 | Bst1 | H2.Q6 | Cdc42se2 |
| | | | | | Cysltr1 | Fcgr4 | C1qbp | H2.Q7 | Cfp |
| | | | | | Daglb | Gpr160 | C3ar1 | H2.T22 | Clcn3 |
| | | | | | Ecscr | Gpr183 | C5ar1 | H2.T23 | Clec7a |
| | | | | | Emr1 | Gsdmd | Calr | H2.T24 | Cln3 |
| | | | | | Entpd1 | H2.Oa | Camk1d | H2.T3 | Cnn2 |
| | | | | | Fcer1g | H2.Ob | Capn7 | Hfe | Colec12 |
| | | | | | Fcgr1 | H2.T23 | Card10 | Icam1 | Coro1a |
| | | | | | Fcgr2b | Havcr2 | Ccdc141 | Ide | Coro1c |
| | | | | | Fcgr3 | Hfe | Ccl12 | Ifi30 | Csk |
| | | | | | Fcgr4 | I830077J02Rik | Ccl17 | Ighm | Cyba |
| | | | | | Fcrl1 | Icam1 | Ccl2 | Mfsd6 | Dnm2 |
| | | | | | Fcrls | Icosl | Ccl24 | Mr1 | Dock1 |
| | | | | | Gpr160 | Ifitm6 | Ccl25 | Nod1 | Dock2 |
| | | | | | Gpr183 | Ifngr1 | Ccl3 | Nod2 | Eif2ak1 |
| | | | | | Gpr34 | Igsf6 | Ccl4 | Pdia3 | Elmo1 |
| | | | | | Gpr56 | Il10ra | Ccl5 | Pikfyve | Elmo2 |
| | | | | | Gpr84 | Il21r | Ccl6 | Psap | Elmo3 |
| | | | | | Gsdmd | Il4ra | Ccl7 | Psmb8 | Fcer1g |
| | | | | | H2-Oa | Il6ra | Ccl8 | Psmb9 | Fcgr1 |
| | | | | | H2-Ob | Itgam | Ccl9 | Psme1 | Fcgr2b |
| | | | | | H2-T23 | Itgb2 | Ccr1 | Psme2 | Fcgr3 |
| | | | | | Havcr2 | Itgb5 | Ccr2 | Ptpn22 | Fgr |
| | | | | | Hfe | Kctd12 | Ccr7 | Pycard | Gas6 |
| | | | | | I830077J02Rik | Lair1 | Cd200 | Rab10 | Gata2 |
| | | | | | Icam1 | Laptm5 | Cd200r1 | Rab27a | Gsn |
| | | | | | Icam4 | Lat2 | Cd40 | Rab32 | Gulp1 |
| | | | | | Icosl | Lgals9 | Cd63 | Rab33a | Hck |
| | | | | | Ifitm6 | Lilra5 | Cd74 | Rab34 | Hmgb1 |
| | | | | | Ifngr1 | Ly86 | Cd81 | Rab35 | Hspa8 |
| | | | | | Igsf6 | Nckap1l | Cd9 | Rab3b | Ighm |
| | | | | | Il10ra | Nfam1 | Cdh13 | Rab3c | Il15 |
| | | | | | Il21r | Ninj1 | Cdh5 | Rab4a | Il15ra |
| | | | | | Il4ra | Numb | Cdk5 | Rab5b | Il1b |
| | | | | | Il6ra | P2ry6 | Cdk5r1 | Rab6a | Il2rg |
| | | | | | Itgam | Pilra | Ceacam1 | Rab8b | Irf8 |
| | | | | | Itgb2 | Plekho1 | Cib1 | Relb | Itga2 |
| | | | | | Itgb5 | Pmepa1 | Cklf | Rftn1 | Itgal |
| | | | | | Kctd12 | Pmp22 | Clasp1 | Slc11a1 | Itgam |
| | | | | | Lag3 | Ptafr | Clasp2 | Tap1 | Itgav |
| | | | | | Lair1 | Ptprc | Clec14a | Tap2 | Itgb1 |
| | | | | | Laptm5 | Selplg | Cmklr1 | Tapbp | Itgb2 |
| | | | | | Lat2 | Sirpa | Cntn2 | Tapbpl | Itgb3 |
| | | | | | Lgals9 | Slamf9 | Col3a1 | Thbs1 | Jmjd6 |
| | | | | | Lilra5 | Slc11a1 | Coro1b | Traf6 | Lbp |
| | | | | | Liph | Slc16a3 | Coro1c | Trem2 | Ldlr |
| | | | | | Lpar5 | Slc44a2 | Creb3 | Unc93b1 | Lepr |
| | | | | | Ltf | Slc46a1 | Csf1 | Was | Letmd1 |
| | | | | | Ly86 | Slc7a7 | Csf3r | Wdfy4 | Lman2 |
| | | | | | Nckap1l | Slc7a8 | Csnk2b | Ythdf1 | Lrp1 |
| | | | | | Nfam1 | Slco2b1 | Ctnnb1 | H2.M5 | Lyar |
| | | | | | Ninj1 | Stab1 | Ctsh | Kdm5d | Lyst |
| | | | | | Numb | Tcirg1 | Cul5 | Flt3 | Masp2 |
| | | | | | P2ry12 | Tgfbr2 | Cx3cl1 | Gm11127 | Megf10 |
| | | | | | P2ry13 | Tlr13 | Cxadr | H2.Ob | Mertk |
| | | | | | P2ry6 | Tlr2 | Cxcl1 | H2.Eb2 | Met |
| | | | | | Pilra | Tlr4 | Cxcl10 | H2.M2 | Mex3b |
| | | | | | Plekho1 | Tlr6 | Cxcl12 | H2.Q1 | Mfge8 |
| | | | | | Plxdc2 | Tlr7 | Cxcl13 | H2.Q2 | Msr1 |
| | | | | | Pmepa1 | Tmem173 | Cxcl2 | Treml4 | Myd88 |
| | | | | | Pmp22 | Tmem206 | Cxcl9 | Ifng | Myh9 |
| | | | | | Ptafr | Tmem37 | Cxcr4 | Clec4b1 | Myo18a |
| | | | | | Ptprc | Tnfrsf13b | Cyp1b1 | Ctse | Myo1g |
| | | | | | Selplg | Tnfrsf1a | Daam2 | H2.M9 | Myo7a |
| | | | | | Siglec5 | Tnfrsf1b | Dab1 | Azgp1 | Ncf2 |
| | | | | | Siglece | Trem2 | Dab2ip | Gm7030 | Ncf4 |
| | | | | | Siglech | Trpv2 | Dcn | Ccl21a | Nckap1l |
| | | | | | Sirpa | Tyrobp | Dicer1 | | Nod2 |
| | | | | | Slamf9 | Icam4 | Disc1 | | Nr1h3 |
| | | | | | Slc11a1 | Cd79b | Dixdc1 | | P2rx7 |
| | | | | | Slc16a3 | Siglece | Dnaja4 | | P2ry6 |
| | | | | | Slc2a5 | Tlr1 | Dnm1l | | Pear1 |
| | | | | | Slc44a2 | C5ar2 | Dock1 | | Pecam1 |
| | | | | | Slc46a1 | Cd101 | Dock5 | | Pik3ca |
| | | | | | Slc7a7 | Liph | Dpp4 | | Pikfyve |
| | | | | | Slc7a8 | Tlr12 | Dusp1 | | Pip5k1c |
| | | | | | Slco2b1 | Tnfrsf17 | Dusp10 | | Pla2g5 |
| | | | | | Stab1 | Upk1b | Edn1 | | Plcg2 |
| | | | | | Tcirg1 | Fcrl1 | Edn3 | | Pld4 |
| | | | | | Tgfbr1 | Tmem8c | Ednra | | Pot1b |
| | | | | | Tgfbr2 | Clec4b1 | Ednrb | | Prkcg |
| | | | | | Tlr1 | Lpar5 | Efhc1 | | Pros1 |
| | | | | | Tlr12 | 4632428N05Rik | Efna1 | | Pten |
| | | | | | Tlr13 | AF251705 | Efnb2 | | Ptk2 |
| | | | | | Tlr2 | Emr1 | Egf | | Ptprc |
| | | | | | Tlr4 | Gpr56 | Egfr | | Ptprj |
| | | | | | Tlr6 | Siglec5 | Emc10 | | Ptx3 |
| | | | | | Tlr7 | Ltf | Emilin1 | | Pycard |
| | | | | | Tmem119 | | Emp2 | | Rab11fip2 |
| | | | | | Tmem173 | | Emx2 | | Rab14 |
| | | | | | Tmem206 | | Enpp1 | | Rab20 |
| | | | | | Tmem37 | | Enpp2 | | Rab27a |
| | | | | | Tmem8c | | Epb41l4b | | Rab31 |
| | | | | | Tnfrsf13b | | Epb41l5 | | Rab34 |
| | | | | | Tnfrsf17 | | Epha2 | | Rab5a |
| | | | | | Tnfrsf1a | | Ephb4 | | Rab7 |
| | | | | | Tnfrsf1b | | Ets1 | | Rab7b |
| | | | | | Trem2 | | Evl | | Rac1 |
| | | | | | Trpv2 | | Fap | | Rac2 |
| | | | | | Tyrobp | | Fbxo45 | | Rack1 |
| | | | | | Upk1b | | Fbxw7 | | Rap1a |
| | | | | | | | Fcer1g | | Rap1gap |
| | | | | | | | Fcgr3 | | Rapgef1 |
| | | | | | | | Fgf1 | | Rara |
| | | | | | | | Fgf2 | | Rhobtb1 |

| | | | | | | | | | |
|---|---|---|---|---|---|---|---|---|---|
| | | | | | | | Fgf7 | | Rhobtb2 |
| | | | | | | | Fgfr1 | | Rhog |
| | | | | | | | Flt1 | | Rhoh |
| | | | | | | | Flt4 | | Scarb1 |
| | | | | | | | Foxc2 | | Sh3bp1 |
| | | | | | | | Foxg1 | | Sirpa |
| | | | | | | | Foxp1 | | Slc11a1 |
| | | | | | | | Fstl1 | | Snx3 |
| | | | | | | | Fut10 | | Sod1 |
| | | | | | | | Gab2 | | Spg11 |
| | | | | | | | Gadd45a | | Sphk1 |
| | | | | | | | Gata2 | | Stap1 |
| | | | | | | | Gbf1 | | Syk |
| | | | | | | | Gipc1 | | Syt11 |
| | | | | | | | Gli3 | | Syt7 |
| | | | | | | | Glipr2 | | Tgfb1 |
| | | | | | | | Glul | | Tgm2 |
| | | | | | | | Gpi1 | | Thbs1 |
| | | | | | | | Gpld1 | | Tlr2 |
| | | | | | | | Gpr35 | | Tlr4 |
| | | | | | | | Gpx1 | | Tm9sf4 |
| | | | | | | | Grn | | Tmem175 |
| | | | | | | | Has2 | | Tnf |
| | | | | | | | Hbegf | | Trbc2 |
| | | | | | | | Hc | | Trem2 |
| | | | | | | | Hdac5 | | Tub |
| | | | | | | | Hdac6 | | Tusc2 |
| | | | | | | | Hdac7 | | Tyro3 |
| | | | | | | | Hdac9 | | Tyrobp |
| | | | | | | | Hif1a | | Unc13d |
| | | | | | | | Hmgb1 | | Vamp7 |
| | | | | | | | Hmox1 | | Vav1 |
| | | | | | | | Hspb1 | | Xkr5 |
| | | | | | | | Hyal1 | | Xkr6 |
| | | | | | | | Igf1 | | Xkr8 |
| | | | | | | | Igf2 | | Fer1l5 |
| | | | | | | | Il1b | | Icam5 |
| | | | | | | | Il34 | | Trbc1 |
| | | | | | | | Iqsec1 | | Apoa2 |
| | | | | | | | Itga1 | | Il2rb |
| | | | | | | | Itga2 | | Mbl2 |
| | | | | | | | Itga3 | | Cfb |
| | | | | | | | Itga9 | | Siglece |
| | | | | | | | Itgam | | Sirpb1b |
| | | | | | | | Itgav | | Sirpb1c |
| | | | | | | | Itgb1 | | Ticam2 |
| | | | | | | | Itgb1bp1 | | Xkr4 |
| | | | | | | | Itgb2 | | Gm9733 |
| | | | | | | | Itgb3 | | Dysf |
| | | | | | | | Jam3 | | Gm5150 |
| | | | | | | | Jun | | Ighd |
| | | | | | | | Jup | | Igkc |
| | | | | | | | Kank1 | | Iglc3 |
| | | | | | | | Kank2 | | Rab39 |
| | | | | | | | Kdr | | Sirpb1a |
| | | | | | | | Kit | | Slamf1 |
| | | | | | | | Klf4 | | Treml4 |
| | | | | | | | Krit1 | | F2rl1 |
| | | | | | | | Lamb1 | | Ifng |
| | | | | | | | Lbp | | Marco |
| | | | | | | | Lemd3 | | Prtn3 |
| | | | | | | | Lgals3 | | Ager |
| | | | | | | | Lgals8 | | Apoa1 |
| | | | | | | | Loxl2 | | Cd209b |
| | | | | | | | Lpxn | | Iglc1 |
| | | | | | | | Lrp6 | | Iglc2 |
| | | | | | | | Lrp8 | | Masp1 |
| | | | | | | | Lrrk2 | | Spon2 |
| | | | | | | | Lyn | | Srpx |
| | | | | | | | Macf1 | | Tulp1 |
| | | | | | | | Map2k3 | | Alox15 |
| | | | | | | | Map2k5 | | Havcr1 |
| | | | | | | | Map3k3 | | Timd4 |
| | | | | | | | Map4k4 | | Trdc |
| | | | | | | | Mapk1 | | Colec11 |
| | | | | | | | Mapk3 | | |
| | | | | | | | Mapre2 | | |
| | | | | | | | Mboat7 | | |
| | | | | | | | Mcc | | |
| | | | | | | | Mcu | | |
| | | | | | | | Mdk | | |
| | | | | | | | Mecp2 | | |
| | | | | | | | Mef2c | | |
| | | | | | | | Met | | |
| | | | | | | | Mia3 | | |
| | | | | | | | Mif | | |
| | | | | | | | Mmp14 | | |
| | | | | | | | Mmp28 | | |
| | | | | | | | Mmrn2 | | |
| | | | | | | | Mospd2 | | |
| | | | | | | | Mpp1 | | |
| | | | | | | | Mtor | | |
| | | | | | | | Mtus1 | | |
| | | | | | | | Myh9 | | |
| | | | | | | | Myo9b | | |
| | | | | | | | Nanos1 | | |
| | | | | | | | Nbl1 | | |
| | | | | | | | Nckap1l | | |
| | | | | | | | Ndel1 | | |
| | | | | | | | Nf1 | | |
| | | | | | | | Nfe2l2 | | |
| | | | | | | | Ninj1 | | |
| | | | | | | | Nod2 | | |
| | | | | | | | Nos3 | | |
| | | | | | | | Notch1 | | |
| | | | | | | | Nr2e1 | | |
| | | | | | | | Nr2f2 | | |
| | | | | | | | Nr4a1 | | |
| | | | | | | | Nrp1 | | |
| | | | | | | | Nup85 | | |
| | | | | | | | Nus1 | | |
| | | | | | | | P2rx4 | | |
| | | | | | | | Pafah1b1 | | |
| | | | | | | | Patz1 | | |
| | | | | | | | Paxip1 | | |
| | | | | | | | Pdcd10 | | |
| | | | | | | | Pdcd6 | | |
| | | | | | | | Pde4b | | |
| | | | | | | | Pde4d | | |
| | | | | | | | Pdgfb | | |
| | | | | | | | Pdpk1 | | |
| | | | | | | | Pecam1 | | |
| | | | | | | | Pex13 | | |
| | | | | | | | Pex5 | | |
| | | | | | | | Pf4 | | |
| | | | | | | | Pfn1 | | |
| | | | | | | | Pfn2 | | |
| | | | | | | | Pik3c2a | | |
| | | | | | | | Pik3cb | | |
| | | | | | | | Pik3cd | | |
| | | | | | | | Pik3cg | | |
| | | | | | | | Pik3r2 | | |
| | | | | | | | Pikfyve | | |
| | | | | | | | Pkn1 | | |
| | | | | | | | Pkn2 | | |
| | | | | | | | Pkn3 | | |
| | | | | | | | Pla2g7 | | |
| | | | | | | | Plcg1 | | |
| | | | | | | | Plcg2 | | |
| | | | | | | | Plekhg5 | | |
| | | | | | | | Plk2 | | |
| | | | | | | | Plpp3 | | |
| | | | | | | | Plxnd1 | | |
| | | | | | | | Pou3f2 | | |
| | | | | | | | Pou3f3 | | |
| | | | | | | | Ppard | | |
| | | | | | | | Ppia | | |
| | | | | | | | Ppib | | |
| | | | | | | | Ppm1f | | |
| | | | | | | | Prcp | | |

| | | | | | | | |
|---|---|---|---|---|---|---|---|
| | | | | | | | Prex1 |
| | | | | | | | Prkca |
| | | | | | | | Prkce |
| | | | | | | | Prkd1 |
| | | | | | | | Prkd2 |
| | | | | | | | Prox1 |
| | | | | | | | Psen1 |
| | | | | | | | Pten |
| | | | | | | | Ptgs2 |
| | | | | | | | Ptk2 |
| | | | | | | | Ptk2b |
| | | | | | | | Ptp4a3 |
| | | | | | | | Ptpn11 |
| | | | | | | | Ptpn23 |
| | | | | | | | Ptprg |
| | | | | | | | Ptprj |
| | | | | | | | Ptprm |
| | | | | | | | Ptpro |
| | | | | | | | Pxn |
| | | | | | | | Rab11a |
| | | | | | | | Rab13 |
| | | | | | | | Rac1 |
| | | | | | | | Rac2 |
| | | | | | | | Rarres2 |
| | | | | | | | Reln |
| | | | | | | | Rgcc |
| | | | | | | | Rhoa |
| | | | | | | | Rhob |
| | | | | | | | Rhoj |
| | | | | | | | Rin2 |
| | | | | | | | Rnf7 |
| | | | | | | | Robo1 |
| | | | | | | | Rock1 |
| | | | | | | | Rock2 |
| | | | | | | | Rpl13a |
| | | | | | | | Rps19 |
| | | | | | | | Rras |
| | | | | | | | Rreb1 |
| | | | | | | | Rtn4 |
| | | | | | | | Sash1 |
| | | | | | | | Scarb1 |
| | | | | | | | Sell |
| | | | | | | | Sema3a |
| | | | | | | | Sema4a |
| | | | | | | | Sema5a |
| | | | | | | | Serpine1 |
| | | | | | | | Serpinf1 |
| | | | | | | | Sh3bp1 |
| | | | | | | | Sirt1 |
| | | | | | | | Slc37a4 |
| | | | | | | | Slit2 |
| | | | | | | | Smoc2 |
| | | | | | | | Snai2 |
| | | | | | | | Socs7 |
| | | | | | | | Sox9 |
| | | | | | | | Sp1 |
| | | | | | | | Sp100 |
| | | | | | | | Sparc |
| | | | | | | | Spp1 |
| | | | | | | | Spred1 |
| | | | | | | | Src |
| | | | | | | | Srf |
| | | | | | | | Srgap2 |
| | | | | | | | Srpx2 |
| | | | | | | | Stap1 |
| | | | | | | | Stard13 |
| | | | | | | | Stat5a |
| | | | | | | | Sun1 |
| | | | | | | | Sun2 |
| | | | | | | | Svbp |
| | | | | | | | Syk |
| | | | | | | | Syne2 |
| | | | | | | | Synj2bp |
| | | | | | | | Tbxa2r |
| | | | | | | | Tek |
| | | | | | | | Tesk1 |
| | | | | | | | Tgfb1 |
| | | | | | | | Tgfb2 |
| | | | | | | | Tgfbr2 |
| | | | | | | | Tgfbr3 |
| | | | | | | | Thbs1 |
| | | | | | | | Tirap |
| | | | | | | | Tnf |
| | | | | | | | Tnfaip6 |
| | | | | | | | Tnr |
| | | | | | | | Trem1 |
| | | | | | | | Trem2 |
| | | | | | | | Trpv4 |
| | | | | | | | Tyro3 |
| | | | | | | | Vash1 |
| | | | | | | | Vav1 |
| | | | | | | | Vav3 |
| | | | | | | | Vegfa |
| | | | | | | | Vhl |
| | | | | | | | Vil1 |
| | | | | | | | Wdpcp |
| | | | | | | | Wnt5a |
| | | | | | | | Zc3h12a |
| | | | | | | | Zeb2 |
| | | | | | | | Zfp580 |
| | | | | | | | Zmiz1 |
| | | | | | | | Il23a |
| | | | | | | | Mst1 |
| | | | | | | | Ccl22 |
| | | | | | | | Xcl1 |
| | | | | | | | Apoh |
| | | | | | | | Cxcl3 |
| | | | | | | | Gata3 |
| | | | | | | | Gdf2 |
| | | | | | | | S100a14 |
| | | | | | | | Arx |
| | | | | | | | Bmper |
| | | | | | | | Ccl11 |
| | | | | | | | Dcx |
| | | | | | | | Dll4 |
| | | | | | | | Fezf2 |
| | | | | | | | Fgf13 |
| | | | | | | | Lcn2 |
| | | | | | | | Ptprr |
| | | | | | | | Stc1 |
| | | | | | | | Thbs4 |
| | | | | | | | Wnt7a |
| | | | | | | | Cdk5r2 |
| | | | | | | | Bmp4 |
| | | | | | | | C5ar2 |
| | | | | | | | Ccr6 |
| | | | | | | | Dysf |
| | | | | | | | Nrg1 |
| | | | | | | | S100a8 |
| | | | | | | | Slamf1 |
| | | | | | | | Slamf8 |
| | | | | | | | Trem3 |
| | | | | | | | Vegfc |
| | | | | | | | Vstm4 |
| | | | | | | | Mcoln2 |
| | | | | | | | Dapk2 |
| | | | | | | | Ccl1 |
| | | | | | | | Ifng |
| | | | | | | | Tac1 |
| | | | | | | | Tnfsf11 |
| | | | | | | | Ager |
| | | | | | | | Alox12 |
| | | | | | | | Apoa1 |
| | | | | | | | Cxcl5 |
| | | | | | | | Defb25 |
| | | | | | | | Fgf18 |
| | | | | | | | Fgfbp1 |
| | | | | | | | Foxb1 |
| | | | | | | | Lgals12 |
| | | | | | | | Mdga1 |
| | | | | | | | Slit1 |
| | | | | | | | Adtrp |
| | | | | | | | Ccbe1 |
| | | | | | | | Drd1 |
| | | | | | | | Fgf10 |
| | | | | | | | Lhx6 |
| | | | | | | | Lrg1 |
| | | | | | | | Meox2 |
| | | | | | | | Mmp9 |
| | | | | | | | Msmp |
| | | | | | | | Mstn |
| | | | | | | | Ror2 |
| | | | | | | | S100a9 |
| | | | | | | | Tacr1 |
| | | | | | | | Cxcr2 |
| | | | | | | | Egr3 |
| | | | | | | | Trim55 |
| | | | | | | | Nrg3 |
| | | | | | | | Fermt1 |
| | | | | | | | Ccl21a |

**Table S8.** Lists of differentially expressed genes between TMZ treated and control P3 PDOXs per each major cell type analysed.
Differentially expressed genes were defined at threshold: FDR <=0.01 and |log2FC|>=0.5
Genes are listed from the highest to lowest $|\log_2FC|$

| Cell origin | mouse TME | | | | | | human GBM | |
|---|---|---|---|---|---|---|---|---|
| Cell type | Astrocytes | | Endothelial cells | | Myeloid cells | | Tumor cells | |
| Fold change | upregulated | downregulated | upregulated | downregulated | upregulated | downregulated | upregulated | downregulated |
| Number of genes | 2 | 1 | 41 | 47 | 127 | 28 | 90 | 78 |
| | Scg2 | Gm12222 | RP23.269H21.1 | Ttr | Rnaset2b | Rpl26 | GDF15 | BCAN |
| | Aldoc | | Rnaset2b | Rpl26 | Gm14513 | Rpl35a | CDKN1A | ID3 |
| | | | mt.Tc | Rpl35a | Cxcl13 | Igfbp7 | SCG2 | C1orf61 |
| | | | Gm7638 | Hexb | Gm7638 | Ttr | CHI3L1 | HES5 |
| | | | Abcb1a | Apln | Pde3b | Ptn | NEAT1 | CSPG5 |
| | | | Cxcl12 | Rpl39.ps | Ptgs1 | Gng5 | GADD45A | FABP7 |
| | | | Pcp4l1 | Ctsd | Jund | Rps15a.ps5 | RPS27L | HMGCS1 |
| | | | Paqr5 | Cwc22 | Slco2b1 | Gm10076 | FTL | CST3 |
| | | | Timp3 | Lrp8 | Spp1 | Rpl22.ps1 | MDM2 | HSPA1A |
| | | | Slc6a6 | Myh10 | Zfhx3 | Enpp2 | DDB2 | HSPA1B |
| | | | H2.D1 | Ctss | Rpsa | Gm12338 | GAP43 | NCAN |
| | | | Hsp25.ps1 | Tyrobp | Apoe | Rps15a.ps7 | OCIAD2 | MSMO1 |
| | | | Degs2 | Ndufs5 | P2ry12 | Gm5644 | NRP2 | TTYH1 |
| | | | Fam32a | Rpl36a.ps2 | Tmem119 | Gas5 | S100A6 | ACAT2 |
| | | | Ly6a | Dok4 | Cfl1 | Gm10269 | BTG1 | HEY1 |
| | | | Ppil4 | Hopx | Cnpy2 | Cd52 | PHLDA1 | FDPS |
| | | | Ucp2 | Hspa5 | Cx3cr1 | Rpl36a.ps2 | SPP1 | DBI |
| | | | Hspb1 | Slc7a1 | Rpl9.ps6 | Ndufa4 | SQSTM1 | IDH1 |
| | | | Ly6c1 | Lyz2 | Cited2 | Rpl37rt | TIMP1 | METRN |
| | | | Serinc3 | Gm7266 | Tubgcp5 | Gm4332 | SLC3A2 | LMO4 |
| | | | Sparcl1 | Wwtr1 | Mpc1 | Rplp2 | ZMAT3 | PEA15 |
| | | | Lrg1 | Ccny | Fscn1 | Rpl23 | C6orf141 | MT3 |
| | | | Maoa | Rpl38.ps2 | Ltc4s | Gm4149 | TUBA1C | EDNRB |
| | | | Aldh2 | Rps15a.ps5 | Gm10263 | Rps16.ps2 | IGFBP3 | SCRG1 |
| | | | mt.Rnr1 | Slc38a2 | Mylip | Rps21 | PHPT1 | NDRG2 |
| | | | Tspo | Rps11.ps3 | Ftl1 | Ndufs5 | PHLDA3 | CYP51A1 |
| | | | U2af1 | Gas5 | Aimp1 | Rpl32 | IDS | CNN3 |
| | | | Pltp | Gm10076 | Dtnbp1 | Rps15 | GAS5 | HIST1H4C |
| | | | mt.Rnr2 | Gm15148 | C1qa | | VGF | RPS10 |
| | | | Cd59a | RP23.288C18.3 | Ucp2 | | DDIT3 | YWHAE |
| | | | Gm694 | Cst3 | Pwwp2a | | ARL4C | FDFT1 |
| | | | Mal | Pmepa1 | Rhob | | MALAT1 | FAM181B |
| | | | Gm14513 | Gm10269 | Coro1a | | FDXR | ARC |
| | | | Ly6e | Rflnb | Pik3cg | | RPS19 | ATP1A2 |
| | | | Bsg | Rpl37rt | Zfp90 | | BAX | IDI1 |
| | | | Malat1 | Nostrin | Rpl13 | | LMNA | MARCKS |
| | | | Ifitm2 | Trf | Gpr34 | | GABPB1-AS1 | ASCL1 |
| | | | Utrn | Calr | Frmd4a | | CAMK2D | FOS |
| | | | Ubb | Rps11.ps2 | Ifngr1 | | RCAN1 | GNAI2 |
| | | | Gm11560 | Gm4149 | Scamp2 | | PMEPA1 | SAMD1 |
| | | | Cldn5 | Sparc | Rpl18.ps1 | | IGFBP5 | HES4 |
| | | | | Tmsb4x | Lrp1 | | LONRF2 | SDC3 |
| | | | | Hsp90b1 | Sox4 | | MDK | STMN1 |
| | | | | Gm9794 | Selplg | | ENC1 | IGFBP2 |
| | | | | Tmsb10 | Qpct | | AEN | LRRC17 |
| | | | | Rpl32 | Sltm | | PGM2L1 | TUBB2B |
| | | | | Rplp1 | Rpl10.ps3 | | TNFRSF12A | TMSB4X |
| | | | | | Gpsm3 | | CEBPB | TUBA1B |
| | | | | | Lamtor1 | | ZFAS1 | ITM2C |
| | | | | | Ier3 | | SRPX | NRARP |
| | | | | | Tmem100 | | NUPR1 | MGST3 |
| | | | | | Sdhd | | HBEGF | SQLE |
| | | | | | Commd2 | | YBX3 | SLC1A3 |
| | | | | | Eif4g2 | | ANXA2 | CHMP4B |
| | | | | | Arl4c | | PMAIP1 | ID1 |
| | | | | | Mknk1 | | VMP1 | GFAP |
| | | | | | Dnm2 | | ASCC3 | TMSB15A |
| | | | | | Kctd12 | | RBP1 | PLPP3 |
| | | | | | Scd2 | | PLK3 | CAMTA1 |
| | | | | | Fam76b | | TRIAP1 | GATM |
| | | | | | Ankle2 | | TMSB10 | MDFI |
| | | | | | Scarb2 | | APLP1 | HSPA8 |
| | | | | | Ly6e | | PLXNB2 | HNRNPA2B1 |
| | | | | | Gm15536 | | SLC25A37 | QKI |
| | | | | | Rab10 | | AC010198.2 | SLC6A11 |
| | | | | | Gm10443 | | TMEM158 | ARL6IP6 |
| | | | | | Hpgds | | KCNF1 | TUBB2A |
| | | | | | Mxi1 | | SESN2 | OLIG1 |
| | | | | | Zfand6 | | AL353138.1 | MT1M |
| | | | | | Prdx5 | | CMBL | APCDD1 |
| | | | | | H2.T.ps | | CCND1 | TSPAN3 |
| | | | | | Naa50 | | CIRBP | FGFBP3 |
| | | | | | Tmem86a | | FBXO22 | AC004540.2 |
| | | | | | Slc29a3 | | S100A16 | MT2A |
| | | | | | Abca1 | | SMIM3 | ID2 |
| | | | | | Atp6v0a1 | | JAG1 | MARCKSL1 |
| | | | | | Ninj1 | | LGALS3 | PAFAH1B3 |
| | | | | | Limd2 | | ARHGEF2 | SAPCD2 |
| | | | | | Olfml3 | | PDLIM4 | |
| | | | | | Elf1 | | XPC | |
| | | | | | Prune2 | | TIGAR | |
| | | | | | Arpc4 | | BLOC1S2 | |
| | | | | | Zfp36l1 | | TXNIP | |
| | | | | | Timp2 | | CEBPG | |
| | | | | | Msr1 | | METTL7B | |
| | | | | | Bin1 | | PRKCQ-AS1 | |
| | | | | | Parvg | | QDPR | |
| | | | | | Marcks | | SCG5 | |
| | | | | | Gm8979 | | SH3BGRL3 | |
| | | | | | Ywhae | | PDGFC | |
| | | | | | Phyh | | | |
| | | | | | Rgs19 | | | |
| | | | | | Sp3 | | | |
| | | | | | Arl6ip4 | | | |
| | | | | | Tuba1b | | | |
| | | | | | Ndufs7 | | | |
| | | | | | Cd37 | | | |
| | | | | | Comt | | | |
| | | | | | Brk1 | | | |
| | | | | | Rpl10a | | | |
| | | | | | Sun2 | | | |
| | | | | | Scoc | | | |
| | | | | | Rtn4rl1 | | | |
| | | | | | Colgalt1 | | | |
| | | | | | Crebrf | | | |
| | | | | | Pla2g7 | | | |
| | | | | | H2.D1 | | | |
| | | | | | Abcg1 | | | |
| | | | | | Srgap2 | | | |
| | | | | | Gla | | | |
| | | | | | Gm8399 | | | |
| | | | | | Ube2d3 | | | |
| | | | | | Lgmn | | | |
| | | | | | mt.Co1 | | | |
| | | | | | Bmyc | | | |
| | | | | | Csf1r | | | |
| | | | | | Gm8606 | | | |
| | | | | | Id2 | | | |
| | | | | | Cndp2 | | | |
| | | | | | Zfp62 | | | |
| | | | | | Lyl1 | | | |
| | | | | | Iah1 | | | |
| | | | | | Dock10 | | | |
| | | | | | Fam49b | | | |
| | | | | | Kdm7a | | | |
| | | | | | Psmc4 | | | |
| | | | | | Selenop | | | |
